# Single-cell imaging of the lytic phage life cycle in bacteria

**DOI:** 10.1101/2024.04.11.588870

**Authors:** Charlie Wedd, Ruizhe Li, Temur Yunusov, Aaron Smith, Georgeos Hardo, Michael Hunter, Aparna Kaaraal Mohan, Racha Majed, Diana Fusco, Somenath Bakshi

## Abstract

When a lytic bacteriophage infects a bacterial cell, it commandeers the cell’s resources to replicate, ultimately causing cell lysis and the release of new virions. As phages function as obligate parasites, each stage of the infection process depends on the physiological parameters of the host cell. Given the inherent physiological variability within a population of genetically identical bacterial cells, we ask how the phage infection dynamic reflects such heterogeneity. Here, we introduce a timelapse imaging assay for investigating the dynamics of individual infection steps by a single T7 phage on a single bacterium. This high-throughput, time-resolved assay enables us to monitor the infection progression simultaneously in multiple cells, uncovering substantial heterogeneity at each step and revealing correlations between infection dynamics and the physiological state of the infected cell. Simulations of competing phage populations with different lysis time distributions reveal that heterogeneity in infection dynamics can significantly impact phage fitness, highlighting it as a potential evolutionary driver of phage-bacteria interactions.

## Introduction

Bacteriophages, viruses that infect bacteria, play pivotal roles in shaping bacterial communities in nature and hold significant promise in medicine and biotechnology as biocontrol agents^1,2^. Among them, lytic bacteriophages stand out for their potential in combating antibiotic-resistant infections^2^. As these viruses rely on the host molecular machinery and precursors to proliferate, the infection-to-lysis process intricately depends on the physiological characteristics of the target cell: for instance, factors such as lipopolysaccharide (LPS) composition and receptor density influence the adsorption rate^3,4^; the resources and machinery of the host cell affect the rate of virion replication^5,6^ and the production of lytic agents necessary for cell lysis^7^. Recent single-cell studies have unveiled significant physiological variability among genetically identical bacterial cells^8,9^, raising questions about how this diversity impacts the dynamics of phage infection^10^ and how the variability in infection dynamics, in turn, influences the overall effectiveness of phages in eliminating their target bacteria at a population level^11,12^.

Most of the current methods for studying the dynamics of phage infection steps in bacteria rely on bulk culture approaches and omics analysis^13–15^, which lack the necessary single-cell resolution to analyse cell-to-cell heterogeneity. Innovative adaptations of such approaches can achieve single-cell resolution by using very high dilution factors^16^, but cannot link the measured infection parameter to the physiology of the host cell. Alternatively, cryo-EM imaging enables high-resolution investigation of structural aspects, but cannot offer time-resolved information to monitor the progress of individual infection steps and their interrelations^17,18^. Recent single-cell studies have revealed unprecedented insights into the mechanisms underlying the lytic-lysogenic switch of temperate phages^19^. However, few studies have attempted to analyse lytic phage infection at single-cell level and to understand how the physiological diversity of host cells influences the infection cycle^12,20,21^. To achieve this, a method is needed that can: (i) track each stage of infection initiated by a single phage targeting a solitary living bacterium in a time-resolved fashion, (ii) maintain a spatiotemporally homogeneous environment to isolate the impact of intrinsic variations in host cell physiology on infection parameters, and (iii) be sufficiently high throughput for quantifying the detailed distribution across these target bacterial cells.

Here, we present a novel approach tailored to address these challenges. Our method harnesses a microfluidic platform engineered to maintain isolated populations of target cells under uniform growth conditions, to enable the tracking of infection dynamics as individual cells become infected and lysed by individual phages. Using high-speed scanning time-resolved microscopy^22^, and a combination of fluorescent markers on the model system of phage T7, we are able to follow individual infection events from phage adsorption to cell lysis on individual cells of *Escherichia coli*. The T7–E. coli system serves as an established quantitative model for lytic phage infection, combining extensive genetic and biochemical knowledge with a short and well-characterized infection cycle that facilitates precise temporal measurements. This makes it particularly suitable for dissecting infection dynamics and benchmarking methods applicable to other phage-host pairs. Altogether, the method provides the first quantification of the timing and variability in the dynamics of lytic phage infection steps within the infected host. Moreover, employing this method allows us to correlate the observed fluctuations in infection parameters with the physiological parameters of the infected host, thereby elucidating the source of such variations.

Analysis of these single-cell time-resolved datasets has yielded unprecedented insights into the temporal dynamics and variability of each infection stage, revealing their detailed distributions, interrelationships, and broader implications in terms of selective pressure on phage populations. Results from our simulations show that the details of the distribution of the key parameters of infection kinetics are crucial in determining the competitive fitness of a lytic phage, suggesting that variability in the phage life history parameters could constitute an evolutionary trait that is currently under-explored. Looking forward, we anticipate that our method will offer the opportunity to quantify the distribution of infection parameters, revealing an understanding of phage-bacteria interactions and their evolution previously unattainable.

## Results

### Single-cell imaging of the T7 phage infection cycle

To identify and monitor the timing of the different steps in the T7 life cycle using fluorescence microscopy, we introduced two fluorescent labels in the phage (Fig. 1a). First, we modified the wild-type T7 genome to include a fluorescent reporter of capsid gene expression (gp10A-B, Methods, Supplementary note 1). The capsid genes are among the most highly expressed T7 genes^23^, making them an excellent target to obtain a strong fluorescent signal for phage transcription. mVenus NB, a yellow fluorescent protein (YFP) with a very fast maturation time (4 min)^24^, was selected as suitable fluorescent reporter given the short phage life cycle (15-20 min)^25^. This modified T7 phage (referred to as T7*) was then stained with a DNA binding dye, SYTOX Orange (Fig. 1a, Methods, Supplementary note 2), to visualise phage adsorption to the host cell (*t*_0_) and subsequent genome injection. SYTOX Orange is spectrally compatible with the YFP reporter, does not affect cell growth^26^, and is amongst SYTOX dyes which have previously been used to label the lambda, T7 and Sf6 phage genomes^20,27,28^.

**Fig. 1:**
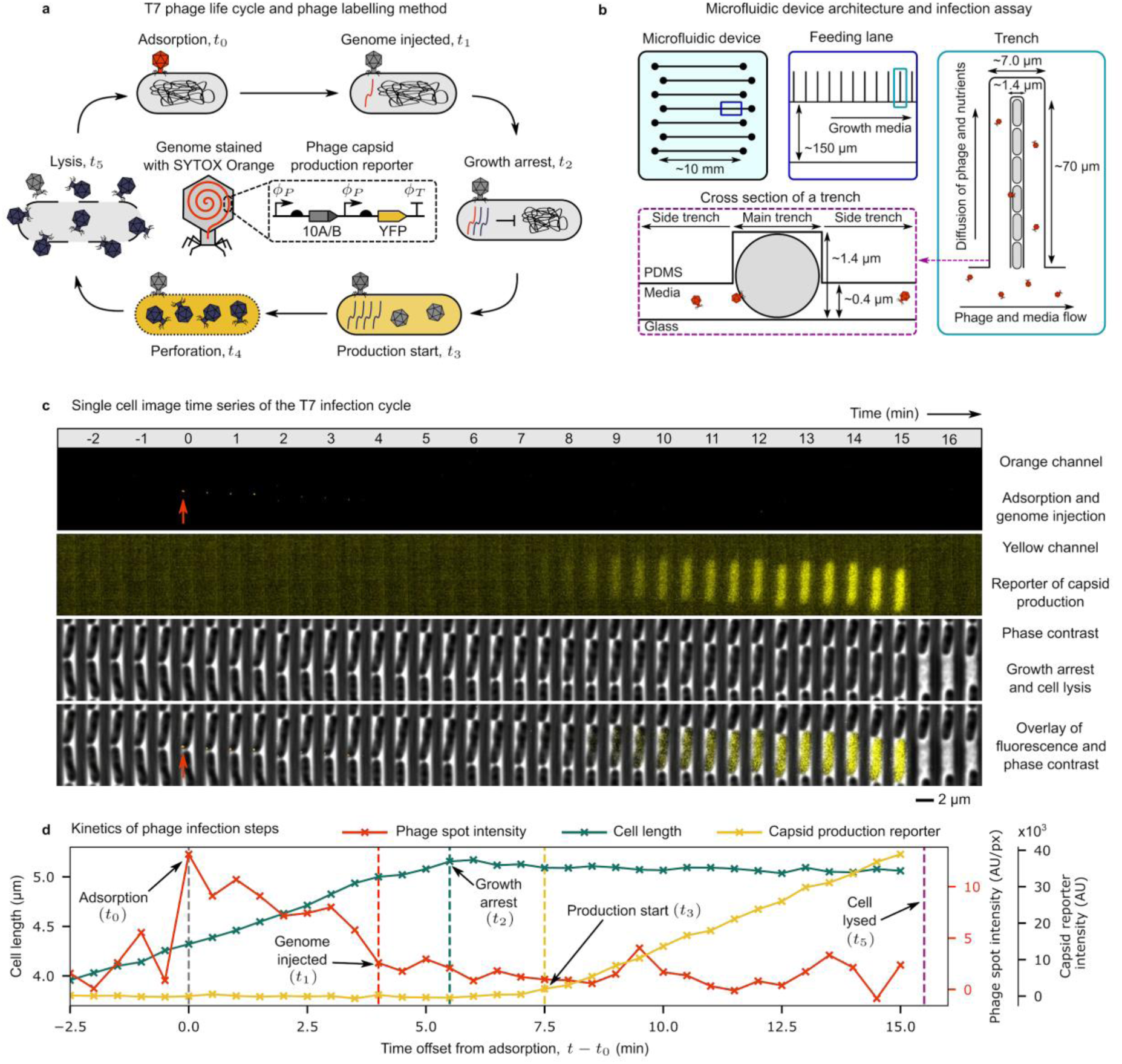
An assay for imaging the T7 phage infection cycle at the single-cell level. **a.** An overview of six key time points in the T7 life cycle, which we use throughout the study to analyse the kinetics of infection steps in the phage life cycle. The DNA staining method and genomic location of the capsid production reporter are indicated in the centre of the loop. **b.** A description of the microfluidic device and infection assay used in this study. **c.** Kymographs showing a T7* phage (indicated by the red arrow) infecting a single bacterial cell. The orange channel image has been bandpass filtered to remove bleedthrough from the YFP capsid production reporter (Supplementary note 6). **d.** Time series data corresponding to the phage infection images presented in (**c**) demonstrate the typical progression of signals during T7* infection. With the exception of perforation, which happens very close to lysis, all time points from Fig. 1a are labelled on the time series. See Supplementary note 7 for a separated panel view of part (**d**).

Individual phage infection cycles in single bacterial cells (*E. coli* MG1655 7740 ΔmotA) were monitored using a modified version of the ‘mother machine’ microfluidic device^29^. In this device, cells are cultivated in linear colonies within narrow (1.4 μm wide) trenches, receiving nutrients diffusively from the media flowing through the orthogonal flow channel (Fig. 1b). In contrast to the regular mother machine design^30^, the narrow trenches are flanked by shallow side trenches that facilitate the diffusion of both nutrients and phage along the length of the trench^29^. We found that the presence of side trenches is essential for phages to infect cells deeper in the trench, so that infection events can be monitored over time all the way to lysis before the corresponding infected cell is pushed out of the trench by the replicating cells above (Supplementary note 3). As individual lineages are isolated in their own trenches and the media continuously flows throughout the experiment, the device maintains cells in exponential growth in a spatiotemporally uniform environment^29^. This uniformity is key to minimise potential sources of heterogeneity arising from a variable environment and accurately quantify the stochasticity of individual steps in the infection process across a bacterial population experiencing identical external conditions. Additionally, as we operate at very low multiplicity of infection, we can ensure that the first lysis events in each trench are truly originating from the infection of one bacterium by one single phage, as evidenced by the rare occurrences of such events (Supplementary movie 1).

The cells are loaded into the device and grown in LB Miller with pluronic for a minimum of three hours to allow them to reach a steady-state exponential growth phase. Subsequently, the media is switched to media with added phage (Methods). High-speed time-resolved scanning microscopy^22^ was used to collect multichannel data at high time-resolution (2 frames min^−1^) during the infection events (Supplementary movie 2). The multichannel timelapse imagedata is then processed using a machine-learning model trained with synthetic micrographs^31^ (Methods, Supplementary note 4), to quantify cell physiology (size, shape, division time, and growth-rate) and infection markers (adsorption, genome injection, host take over, capsid expression, and lysis) over time. Individual infected cells were tracked across frames using a custom-designed lineage-tracking algorithm, which accommodates the disappearance of a subset of cells due to phage-induced lysis (Supplementary note 5). An example of a resultant multichannel kymograph of a single infection event is shown in Fig. 1c and its corresponding time-series data in Fig. 1d. In the orange channel, the adsorption of the SYTOX Orange stained phage to a cell is seen as an orange dot (*t*_0_), which fades and disappears over time as the genome is injected (*t*_1_). In the yellow channel, the YFP signal of the capsid production reporter can be seen increasing in intensity (*t*_3_) after genome injection is completed and up to cell lysis. The phase contrast channel shows the cell growing in length up until a point, post-phage-adsorption, when the growth stops abruptly (*t*_2_), and the cell eventually lyses (*t*_5_). In the following sections, we analyse the dynamics and heterogeneity of each of these steps across different infection events.

### Genome injection dynamics show multiple entry modalities

The molecular mechanisms that lead to T7 genome entry have been extensively studied^32–36^ and result in a three-step process: (i) up to the first 850 base-pairs^37^ enter the cell as the phage tail penetrates the cell wall and membrane, (ii) the host RNA polymerase (RNAP) recognises a series of binding sites on the genome and translocates it while transcribing the early genes, including the T7 RNAP, (iii) once expressed, the T7 RNAP takes over the process and pulls in the remaining 85% of the genome.

Studies in bulk cultures have shown that the whole process takes approximately 10 min in LB medium at 30 °C^38^. Since most of these studies take place at 30 °C, we have used available data^37,38^ to estimate genome entry to take between 5 min and 7 min at 37 °C (Supplementary note 10). However, as these studies used bulk cultures, the variability of its dynamics within a population is unknown. In our setup, we can track such dynamics across multiple infection events, by quantifying the fluorescence signal coming from the stained phage DNA over time (Fig. 2, Methods, Supplementary note 8). When the phage binds to a cell, a bright spot suddenly appears in the orange channel due to the immobilisation of the phage upon adsorption (Fig. 2c). The fluorescence intensity of the spot then decreases over time as the genome gradually leaves the viral capsid and enters the cell (Supplementary movie 3). We have also quantified the photobleaching kinetics of these stained phages in the absence of cells, allowing us to differentiate between photobleaching and genuine genome injection events (Supplementary note 9).

**Fig. 2:**
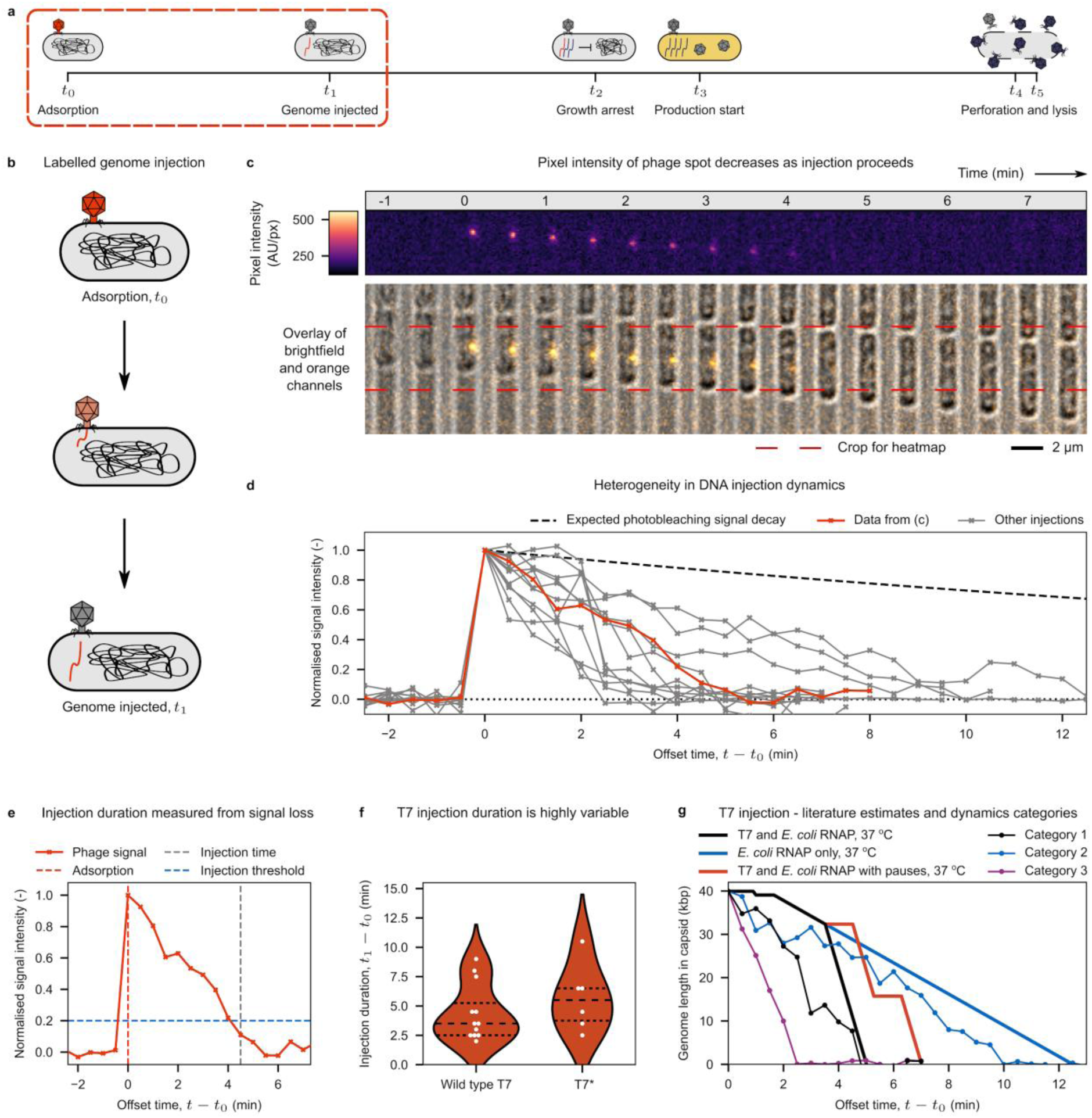
Heterogeneity in phage genome injection dynamics. **a.** A timeline showing the temporal location of genome injection in the T7 life cycle. **b.** A schematic of T7 genome injection. **c.** A kymograph of genome injection shown as a heat map, along with a corresponding overlay of the brightfield and orange channel images. Both the brightfield and orange channels have been uniformly contrast adjusted to enhance visibility. The phage spot moves vertically downwards in sequential frames due to cell growth, but remains bound at the same location on the cell surface. **d.** A comparison of phage signal intensities over time for several genome injection events. Signals have been background corrected and normalised. The injection shown in (**c**) is displayed in orange and examples of 11 other injections are displayed in grey. **e.** An example plot showing how the injection duration is calculated. **f.** Violin plots showing the distribution of genome injection durations (*t*_0_ to *t*_1_) for T7 and T7*. The central dashed line represents the median and the outer dashed lines the first and third quartiles. The mean injection duration is 4.4 min (n = 12, CV = 55%) for T7 and 5.7 min (n = 6, CV = 50%) for T7*. Details of the event selection criteria are found in Supplementary note 11. **g.** Estimates of T7 genome injection dynamics constructed from literature sources^37,38^, shown by the thick solid lines without markers, (Supplementary note 10) have similarities with the measured injection dynamics on single cells. This can be seen by comparison to the example events, representing three of the five categories of genome entry dynamics (Supplementary note 10), which are plotted using solid lines with filled circles. For the example events, the genome length remaining in the capsid is calculated by applying a correction to the normalised fluorescence signal to account for the expected photobleaching (Supplementary note 10).

Across multiple adsorption events, we observed significant heterogeneity in the progression of genome injection (Fig. 2d, 2f, Supplementary note 10). The intensity trends reveal five broad categories of genome injection dynamics (Fig. 2g, Supplementary note 10). In the first category, genome injection starts at approximately 4 kbp min^−1^ for the first 1.5 min to 2.0 min, at which point it transitions to a faster rate. The injection rate in the faster phase fluctuates, but averages at 7 kbp min^−1^ to 12 kbp min^−1^. The second category includes genome injection events which begin similarly to the first category and then continue at an approximately constant rate of 4 kbp min^−1^, representing the slowest of the five categories. In the third and fourth categories, the initial entry rate is considerably higher (Fig. 2g, Supplementary note 10). In the fifth category, most of the genome enters in a single sharp drop at a rate of approximately 60 kbp min^−1^ (Supplementary note 10). The distribution of the injection time duration (Fig. 2f) displays a mean duration time of 4.4 min for T7 and 5.7 min for T7*, consistent with our estimates for genome injection time from previous bulk experiments, with a large variability across infection events (coefficient of variation (CV) = 55%, n = 12 for T7, CV = 50%, n = 6 for T7*). This large variability is likely due to the different kinetic categories we observe, which are considered in detail in the Discussion section

### Dynamics of host cell shutdown and viral takeover are remarkably consistent across infection events

T7 early (class I) genes, transcribed by the host RNAP at the start of infection, encode proteins such as Gp0.4 and Gp0.7 that inhibit cell division and host transcription, while some class II genes contribute to host genome degradation and further inhibit host transcription; together, these activities rapidly shut down host functions^39–43^. We therefore expect cell growth arrest to be among the first signs that phage proteins are being produced. Our fluorescent transcriptional reporter for the capsid proteins, in addition, pinpoints the onset of expression of late (class III) genes from the phage genome. Together, these two markers (growth arrest and capsid expression reporter) allow us to analyse the dynamics and variability in phage protein production during the infection process (Fig. 3a).

**Fig. 3:**
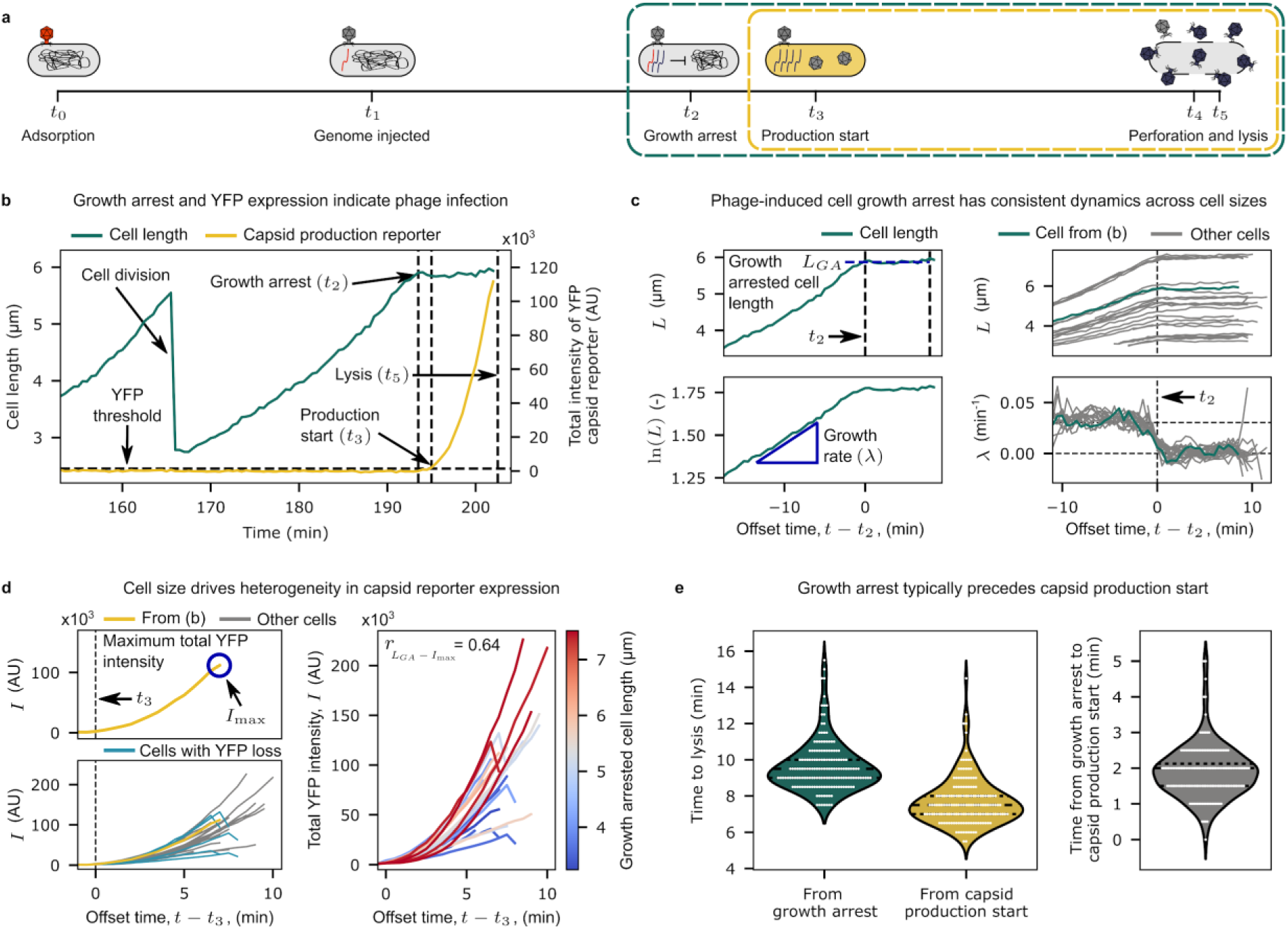
Dynamics of host shutdown and phage gene expression. **a.** A timeline showing the temporal location of growth arrest and capsid production start in the T7 life cycle. **b.** Example time series data showing representative cell length and capsid reporter expression data of a cell which becomes infected. Three time points, the growth arrest (*t*_2_), capsid production start (*t*_3_) and lysis (*t*_5_), are indicated. The threshold intensity used to define production start is also indicated. **c.** We use the growth arrested cell length as a measure of cell size after host takeover (left, top panel). The cell growth rate is calculated as the instantaneous slope of the natural log-transformed cell length (left, bottom panel). Comparison of the cell length (right, top panel) and growth rate (right, bottom panel) between the cell in panel (**b**) (green lines) and 22 other infected cells (grey lines). The growth arrest dynamic is highly consistent between cells. The upper and lower horizontal dashed lines in the bottom right panel show the mean growth rate and zero respectively. **d.** Capsid reporter production is highly variable between different infection events. We use total YFP intensity summed over the cell, *I*, as a measure of capsid reporter production, and hence the maximum total YFP intensity, *I*_*max*_, (left, top panel) as a proxy for the total number of capsid proteins produced in a cell. Example data from 23 infection events (left, bottom panel) shows the variability in capsid reporter production dynamics, comparing the example from panel (**b**) (yellow line) to other events (grey lines). Five events are highlighted in green; these cells show a sharp drop in YFP signal in the final observation before lysis due to the perforation of the cell envelope (Supplementary note 13). The variability in production kinetics is linked to differences in cell size (right). The lines are coloured by the growth arrested cell length of each cell and the maximum total YFP intensity is positively correlated with the growth arrested cell length (*r* = 0.64). **e.** Growth arrest typically precedes production start by a mean time delay of 1.9 min. The violin plots show kernel density estimates of the timings from growth arrest (green) and production start (yellow) to lysis, along with the timings between growth arrest and production start (grey). The central dashed line represents the median and the outer dashed lines the first and third quartiles. The mean time from growth arrest to lysis (*t*_2_ to *t*_5_) is 9.6 min (n = 166, CV = 15%). The mean time from production start to lysis (*t*_3_ to *t*_5_) is 7.7 min (n = 160, CV = 17%). The mean time from growth arrest to production start (*t*_2_ to *t*_3_) is 1.9 min (n = 160, CV = 39%). The slight discrepancy in sample size arises because a small subset of infected cells shifted towards the exit of the trench where segmentation was unreliable due to phase contrast artefacts, making it impossible to accurately determine the timing of production onset (Supplementary note 14).

A representative time series from a single infection event is illustrated in Fig. 3b. The green line depicts cell length, *L*(*t*), where the abrupt periodic drops prior to infection correspond to cell division events. The instantaneous growth rate for each cell, *λ*, is calculated from the local slope of the *ln* (*L*(*t*)) time series (Fig. 3c, Supplementary note 12). The precise moment of growth arrest (*t*_2_) is defined as the point at which the instantaneous growth rate falls below a given threshold (Supplementary note 12). Soon after growth arrest is detected, we observe the level of capsid expression (*I*(*t*), yellow line) to increase rapidly until lysis occurs (Fig. 3b).

Growth arrest dynamics are found to be remarkably robust across different infection events (Fig. 3c). Cells transition from pre-infection growth rates to complete cessation within 3-4 min and in a consistent fashion, independently of their size or position in the cell-cycle. By contrast, expression of the capsid reporter displays considerable variability (Fig. 3d), with the maximum intensity of the reporter, *I*_*max*_, varying by almost an order of magnitude across infection events. We found the variability in *I*_*max*_ to be strongly correlated with the size of the growth arrested cell (*L*_*GA*_) (Fig. 3d, right). Larger production rates in larger cells would be consistent with the presence of more ribosomes, which is likely the limiting factor in phage protein production. We investigate this in the next section.

In five of the 23 example infections presented in Fig. 3d, the YFP intensity sharply decreases in the final observation before lysis (highlighted in green, Fig. 3d). This decline in signal coincides with the perforation of the cell envelope (*t*_4_, detailed in Fig. 5 and Supplementary note 13). Unlike the genome injection process discussed earlier (*t*_0_ to *t*_1_), the time intervals between growth arrest and subsequent lysis (*t*_2_ to *t*_5_) and between start of capsid production and lysis (*t*_3_ to *t*_5_) are narrowly distributed (respectively, CV of 15% and 17%, Fig. 3e), with the latter delayed on average by 1.9 min compared to the first. We note here that the folding and maturation of the reporter proteins can take minutes^24^, implying that actual expression of the capsid genes might start at the same time, if not earlier than the growth arrest of the host cell. Taken together, these results suggest that once the phage has taken control over the cell, the subsequent timing of events follows a highly reproducible sequence with minimal stochastic variation.

### Phage protein production kinetics are strongly coupled with the physiological variability of the host

The observed high variability in phage protein production, despite the high reproducibility of the timing of events, hints to other sources of heterogeneity beyond production time. To understand the source of this heterogeneity and its correlation with the cell size at the moment of growth arrest, *L*_*GA*_ (Fig. 3d), we developed a mathematical model that takes into account viral genome replication, transcription, and translation (Fig. 4a, details in Supplementary note 15). In brief, the model assumes that the phage genome replication is autocatalytic, and the genome replicates exponentially with a rate *λ*_1_. If the number of phage genome copies, rather than T7 RNAP availability, is the limiting factor in transcription, the total amount of mRNA encoding YFP is expected to increase exponentially as the phage genome replicates. The translation rate, and thus YFP production, depends on the available number of mRNA copies, *m*, and the number of ribosomes present in the cell according to a simple Hill equation (Fig. 4a, Supplementary note 15). The maximum translation rate, *λ*_2_, is then a proxy for the total number of active ribosomes in the cell.

**Fig. 4:**
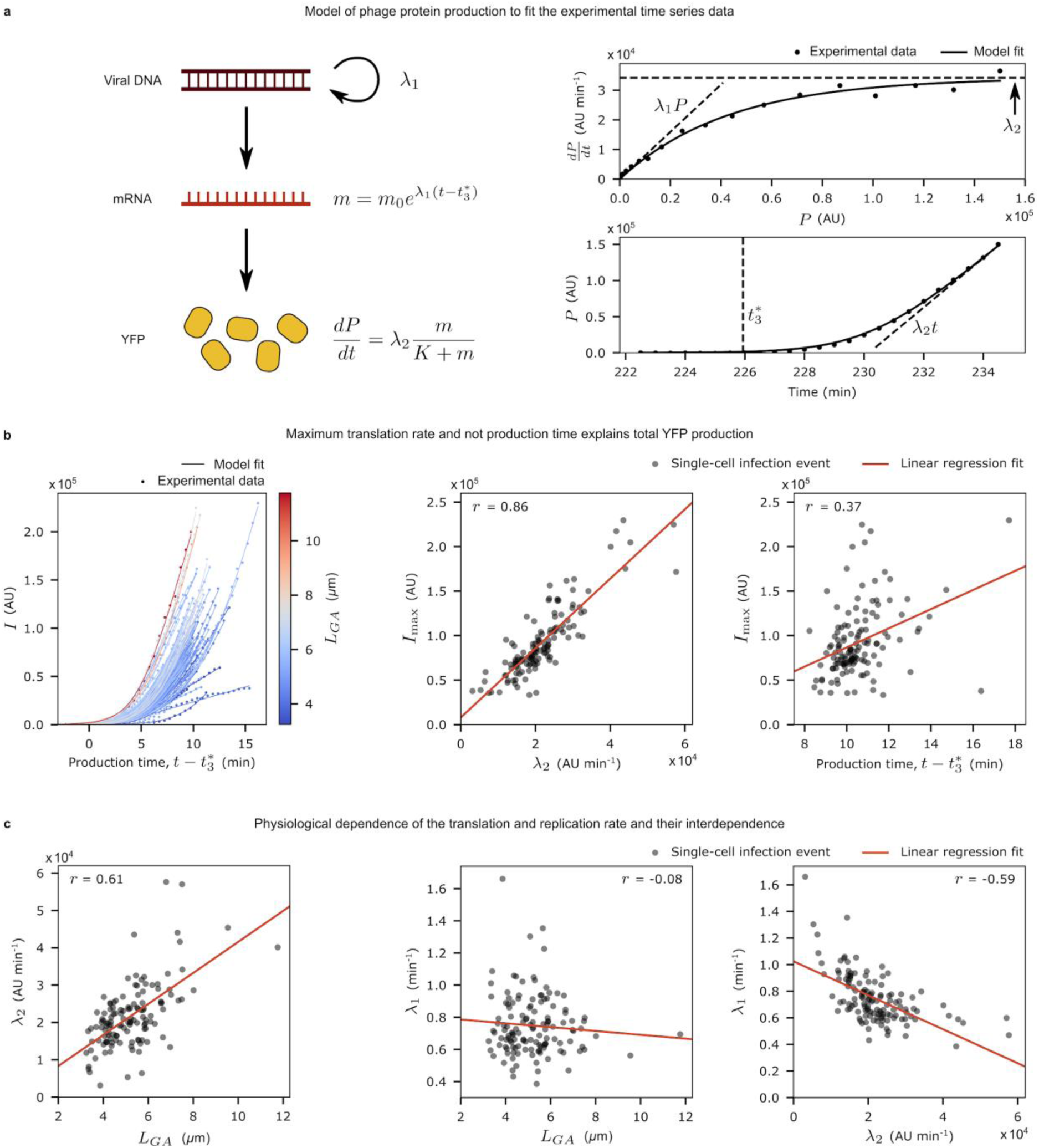
Maximum translation capacity explains phage protein production. **a.** Schematic diagram of the mathematical model for phage protein production from viral genome duplication, through gene transcription to protein translation. The two analytical functions, *dP*/*dt* as a function of *P* (eq. 1) and *P* as a function of time (eq. 2) are sequentially used to fit the experimental data and determine *λ*_1_, *λ*_2_, and production start time, *t*_3_ ∗. **b.** Time-series data of YFP expression are shifted to align production start time (*t*_3_ ∗), with traces coloured according to cell length at growth arrest. The maximum YFP signal observed, *I*_*max*_, strongly correlates with the inferred maximum translation rate *λ*_2_, and only moderately with inferred production time (*t*_3_ ∗ to *t*_5_). **c.** Maximum translation rate correlates with cell length at growth arrest (*L*_*GA*_), linking cell physiology with this parameter. By contrast, phage genome duplication rate, *λ*_1_, shows no correlation with host cell features (Supplementary note 15). Intriguingly, *λ*_1_ and *λ*_2_ show a negative correlation. For parts (**b**) and (**c**), all plots show 130 lysis events, pooled from three independent experiments (Supplementary note 14).

The kinetics of YFP can be solved analytically, providing simple functional forms for *dYFP*/*dt* as a function of YFP and of YFP as a function of time:

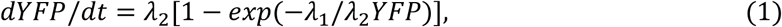

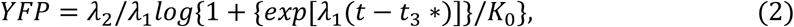

where *K*_0_ = *K*/*m*_0_ (Fig. 4a and Supplementary note 15).

These equations allow us to estimate the model parameters from the experimental fluorescence intensity time series (Fig. 4a, Supplementary note 15). By fitting each cell’s time series, we can extract values for *λ*_1_, *λ*_2_ (from eq. 1), and the time, *t*_3_ ∗ (from eq. 2) at which protein production starts (Fig. 4b). In contrast to *t*_3_, which estimates the start of protein production based on the fluorescent signal increasing above a given threshold and is therefore dependent on the experimental setup sensitivity, *t*_3_ ∗ is robust to variations at low fluorescence levels and can thus provide a more accurate estimate of the true start of protein production. We find that the maximum translation rate, *λ*_2_, strongly correlates with maximum YFP values, pointing at the number of ribosomes in the infected cell as the key variable controlling phage protein production (Fig. 4b). By contrast, production time, defined as *t*_5_ − *t*_3_ ∗, shows only a weak correlation with maximum YFP, in line with the idea that the major source of variability in phage protein production lies in the number of ribosomes of the infected cell and, therefore, the cell’s physiological state at the time of viral takeover, and not the latent period of infection, as previously thought^44–46^.

We first examined how translation kinetics relate to host physiology. Unsurprisingly, we find that *λ*_2_ positively correlates with cell size at the point of growth arrest (Fig. 4c), which explains the empirical correlation between maximum YFP and cell size in the experimental data (Fig. 3d). Importantly, our YFP marker was designed to track capsid protein expression and, as such, is our best proxy for the number of viral capsids produced by the infected cell, i.e., the burst size. Our results therefore suggest that burst size could, in principle, vary by almost one order of magnitude even in perfectly homogeneous conditions, simply because of physiological differences across infected cells.

Intriguingly, we observe a significant negative correlation between *λ*_1_ and *λ*_2_, suggesting the presence of a negative feedback between viral genome replication and translation. A possible explanation is that, if translation is fast, viral capsids might assemble rapidly, spooling in viral DNA and thus depleting the pool of viral genomes that could replicate. This inverse coupling could thus represent a self-limiting mechanism that balances genome amplification with protein synthesis. Testing this hypothesis will require direct time-resolved measurements of genome and capsid dynamics in single cells.

Finally, while cell length at growth arrest partially explains the variability observed in YFP production, we find that maximum YFP concentration, *I*_*max*_/*L*_*GA*_, still exhibits a residual CV of 33%. If holin production was to track YFP production, this degree of variability would be incompatible with the tight distribution of time between growth arrest and lysis (CV of 15%). These results imply that holin accumulation is buffered from global protein synthesis noise. A fluorescent reporter for the holins would be necessary to test this hypothesis.

### Gradual perforation of the cell membrane precedes the abrupt lysis of the infected cell

The concluding stage of the infection process involves the lysis of the host cell. We collected high-frame rate images (100 Hz) to visualise the events preceding cell lysis (Methods, Supplementary movie 4). For this specific analysis, we used secondary infections to achieve sufficient throughput with high-speed imaging. These events likely occur with a multiplicity of infection (MOI) greater than one, however, we assume the timing of the final lysis stage is independent of MOI (Methods). The process of lysis unfolds in two distinct phases. The initial phase, here termed as perforation, involves a gradual leakage of material from the cell, resulting in a subtle increase in phase brightness (Fig. 5b), accompanied by a gradual decrease in the intensity of the surrounding environment. This phase is slower, lasting several seconds, and typically begins with a small but sharp increase in phase brightness (Fig. 5d), marking the onset of perforation (*t*_4_). Material is then steadily lost from the cell until the commencement of the second phase at *t*_5_. The subsequent phase, referred to as lysis, represents the final structural breakdown of the cell envelope. This is indicated by the rapid increase in the phase contrast intensity (purple arrow in Fig. 5d). We observed perforation in all lysing cells (Supplementary note 16), and we have estimated the mean duration between the onset of perforation and the start of lysis (*t*_4_ to *t*_5_) to be 6.56 s.

**Fig. 5:**
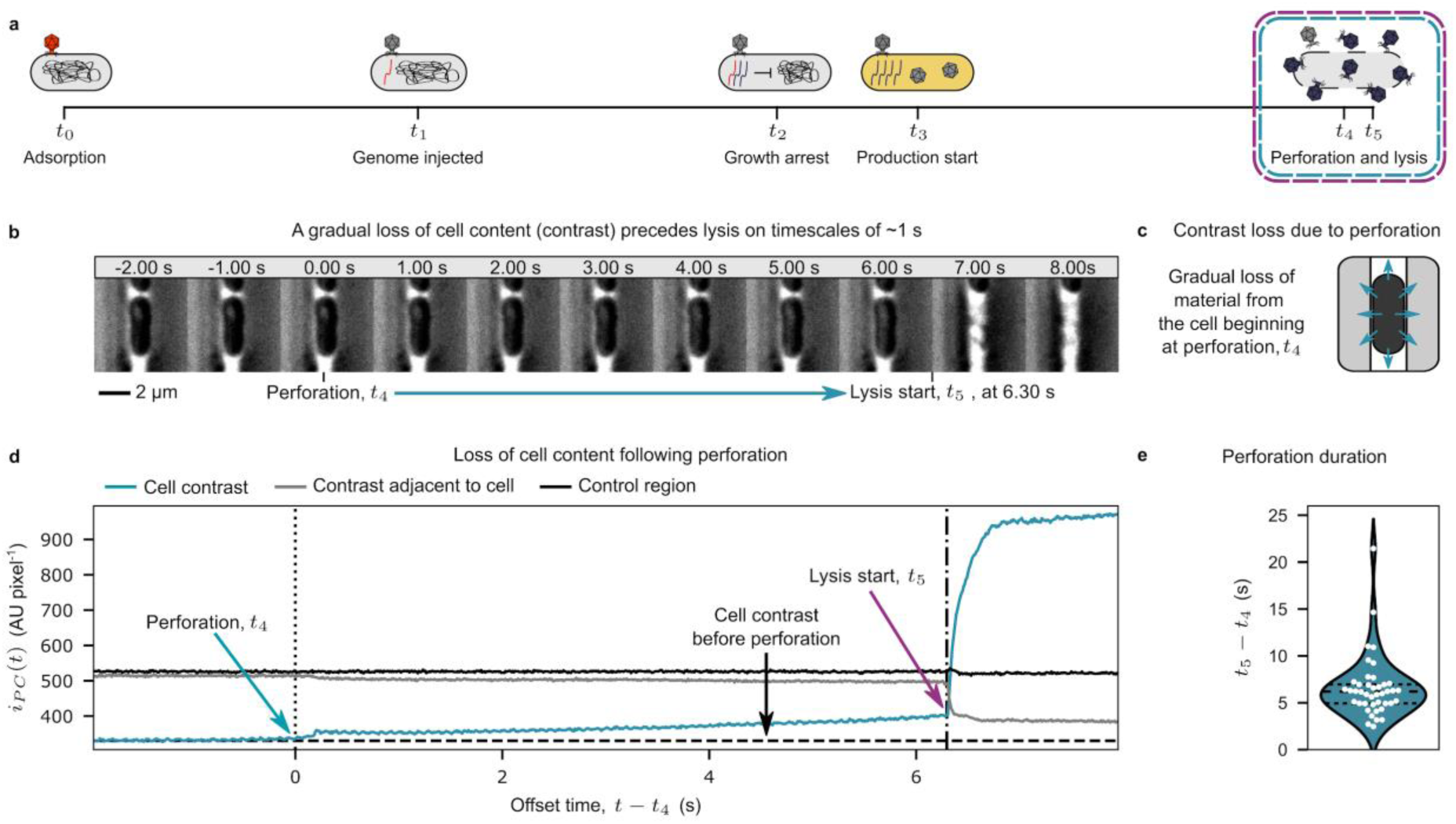
A gradual loss of cell contents precedes cell lysis. **a.** A timeline of the T7 phage life cycle, indicating the timing of perforation (*t*_4_) and lysis (*t*_5_). **b.** A kymograph showing the cell lysis on timescales of 1.00 s. The time axis is offset such that perforation (*t*_4_) starts at time zero. **c.** A schematic illustrating the gradual loss of material from the cell which follows perforation and gives a loss of contrast in the images. **d.** Time series data showing the phase contrast intensity within and outside the cell shown in the timelapse shown in (**b**). **e.** Violin plot showing the distribution of perforation duration across different infection events (perforation start to lysis start, *t*_4_ to *t*_5_). The mean perforation duration is 6.56 s (n = 42, CV = 50%). Details of event selection are provided in Supplementary note 17.

In Fig. 3d, we observe that in some infections, the YFP signal drops in the final observation prior to lysis. This phenomenon is only observed in a fraction of the infection events due to the short interval between perforation and lysis compared to the frame rate of the images taken. With a mean perforation to lysis time of 6.56 s, we only expect to observe the drop in YFP in 21.9% of infections when imaging at 2 frames min^−1^, which is consistent with the results shown in Fig. 3d and the observed increase in phase contrast signal of those cells (indicating a loss of material from the cell) as shown in Fig. S11 (Supplementary note 13).

### Heterogeneity in lysis time is driven by variability in the early stages of infection

Having measured the timing of several points in the infection cycle, we set out to construct a full timeline of the typical T7 infection cycle, alongside a comparison of the relative contributions of each individual infection step to the overall variability in lysis time.

Fig. 6a illustrates a comprehensive timeline of the typical infection cycle of T7* on *E. coli* cells, from phage adsorption onto the cell to the mean of four key time points in the infection cycle: genome injection (*t*_1_), host cell shutdown (*t*_2_), capsid production (*t*_3_), and cell lysis (*t*_5_). Error bars accompany each time point, representing the range of variability (95% confidence interval). The perforation step (*t*_4_) is not represented here due to its negligible duration compared to the other stages (Fig. 5).

**Fig. 6:**
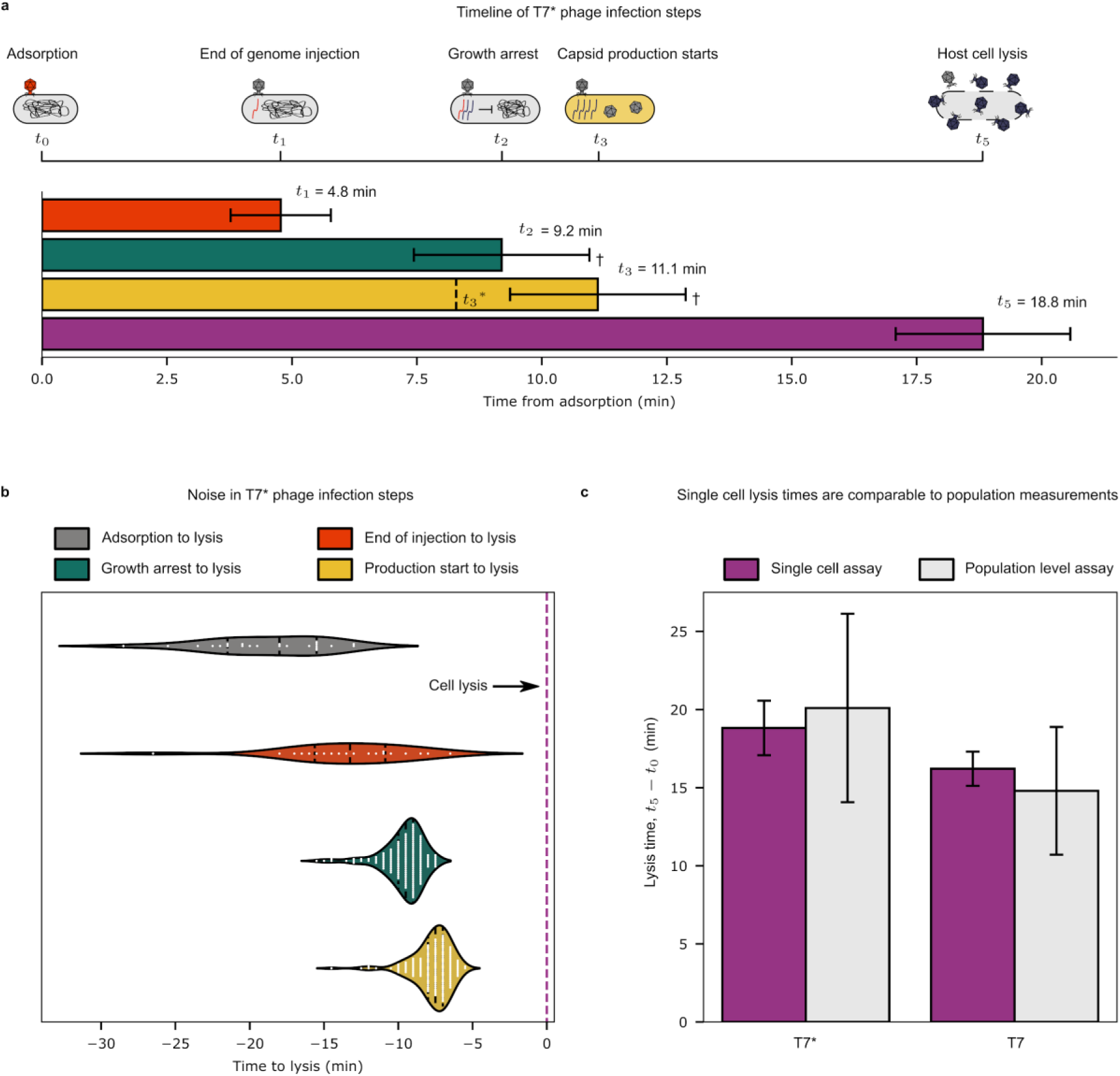
The timing of individual steps of infection and the associated variability across events. **a.** Timeline of phage infection steps. Each bar represents the mean time from adsorption to the indicated time point in the T7* phage life cycle. The corrected production start time from the model results in Fig. 4, *t*_3_ ∗, is indicated as a dashed line on the yellow bar. The data is pooled from three single-cell experiments, giving total sample sizes for the measurement of *t*_0_ → *t*_1_ (injection end) and *t*_0_ → *t*_5_ (lysis) of 20 and 23 respectively. Measurements of *t*_2_ → *t*_5_ (growth arrest to lysis, n = 166) and *t*_3_ → *t*_5_ (production start to lysis, n = 160) were subtracted from the mean lysis time to estimate the mean time for *t*_0_ → *t*_2_ (growth arrest) and *t*_0_ → *t*_3_ (production start). All error bars represent a 95% confidence interval. Confidence intervals for error bars marked with a † are calculated by propagating the standard error from both samples used for the calculation and assuming both samples were independent. Event selection is summarised in Supplementary notes 11 and 14. **b.** Kernel density estimates of the timing distributions from different time points in the T7* phage life cycle to lysis. Each single-cell infection event measured is shown as a white dot. Inner and outer dashed lines in each violin represent the median and lower and upper quartiles respectively. Adsorption to lysis (grey violin) has mean 18.8 min, CV = 21%, n = 23. Injection end to lysis (orange violin) has mean 13.4 min, CV = 33%, n = 20. Growth arrest to lysis (green violin) has mean 9.6 min, CV = 15%, n = 166. Production start to lysis (yellow violin) has mean 7.7 min, CV = 17%, n = 160. The perforation to lysis time is not included, as relative to the timescales of the other steps, the contributed noise from this step is negligible (the mean is 6.56 s, CV = 50%, n = 42). **c.** Lysis times measured with single-cell and population level methods are comparable. Purple bars represent the mean of single-cell lysis times pooled from three experiments, and grey bars represent the mean of three population level lysis time measurements (three biological replicates). The error bars represent the 95% confidence interval. The mean lysis times of T7* obtained from single-cell and population level measurements (18.8 min, n = 23 and 20.1 min, n = 3 respectively) are not significantly different when compared with a t-test. The mean lysis times of T7 obtained from single-cell and population level measurements (16.2 min, n = 7 and 14.8 min, n = 3 respectively) are also not significantly different when compared with a t-test. For all data in Fig. 6, events with an adsorption to lysis time of 30 min or more are treated as outliers and excluded from the distribution. More detail on the single-cell selection criteria is given in Supplementary note 11.

Analysis of this timeline reveals that genome injection typically concludes 4.8 ± 1.0 min after phage adsorption, accounting for just over one quarter of the overall lysis time (18.8 ± 1.7 min). Approximately 9.2 ± 1.8 min into the infection cycle, the host cell ceases growth, then capsid production starts at 11.1 ± 1.8 min after adsorption. However, the corrected production start time inferred from the model (*t*_3_ ∗) is 8.3 ± 1.8 min after adsorption, suggesting that capsid production actually starts slightly before host takeover. Approximately 9.6 ± 0.2 min after the host takeover (measured, Fig. 3e), the cell undergoes lysis, releasing the phage copies into the surrounding environment. The average duration from adsorption to lysis, the lysis time, is 18.8 ± 1.7 min for T7*, consistent with the value obtained from bulk experiments (20.1 ± 6.0 min, n = 3, Fig. 6c, Supplementary note 18). For wild type T7 phage without the genomically integrated capsid expression reporter, we estimated the average time between adsorption and lysis to be 16.2 ± 1.1 min, also consistent with the corresponding bulk average (14.8 ± 4.1 min, n = 3, Fig. 6c, Supplementary note 18). The observed difference in lysis time between the two strains could be related to the additional burden imposed by the expression of the capsid reporter.

Our results show that the lysis time exhibits a CV of 21% across infection events. Fig. 6b illustrates the contribution of each step in the infection cycle towards this variability. Here, all time points are directly measured with reference to cell lysis, as this is the most sudden and clearly identifiable of the events. We find that the variability in the overall lysis time is comparable to the variability in the time interval between the end of injection (*t*_1_) and lysis (*t*_5_). Conversely, both the durations between growth arrest and lysis (*t*_2_ to *t*_5_), and capsid production start and lysis (*t*_3_ and *t*_5_), are very consistent, with interquartile ranges spanning just 1.0 min (<10% of the lysis time). These data suggest that the primary sources of variability in lysis time stem from the initial stages of infection up to the point in which cell growth stops. Further variability has potential to arise from pre-adsorption steps, such as the phage searching for the appropriate receptor. However, our data show that injecting phage typically remain in their initial position, suggesting that such searches may be uncommon. After growth arrest, which delineates the time when the phage has likely taken control of the host cellular machinery, the timing of events is remarkably consistent across infection events.

### Lysis time variability can provide fitness advantages to phage populations

Lysis time represents a key life history parameter for lytic phages and is known to be under strong selective pressure in laboratory experiments^47–51^ and potentially in the wild. Its fitness effect on phage populations in different environments is theoretically well studied, however, due to the lack of experimental data regarding its level of stochasticity, it is typically assumed to be either deterministic, exponentially distributed, or erlang distributed for modelling convenience^52–55^. Our data provide the unprecedented opportunity to quantify lysis time variability, raising the question of whether it can, in principle, represent an evolutionary trait conferring a competitive fitness advantage to a phage population.

To investigate this question, we used stochastic agent-based simulations of a serial passage experiment, in which two phage populations, denoted as “wild type” and “mutant”, are initially mixed in equal proportion and then passaged through several population bottlenecks until one phage approaches fixation (Fig. 7a, Methods, Supplementary note 19). To specifically quantify how distinct distributions in lysis time increase or decrease the ability of a phage to outcompete another, we set the wild type and mutant phage so that they share the same burst size distribution, but have distinct and inheritable lysis time distributions. Some examples of these distributions are shown in Fig. 7c. Mutant 1 has the same mean lysis time as the wild type, but different standard deviation. Mutant 2 has the same mean and standard deviation in lysis time as the wild type, but different skew.

**Fig. 7:**
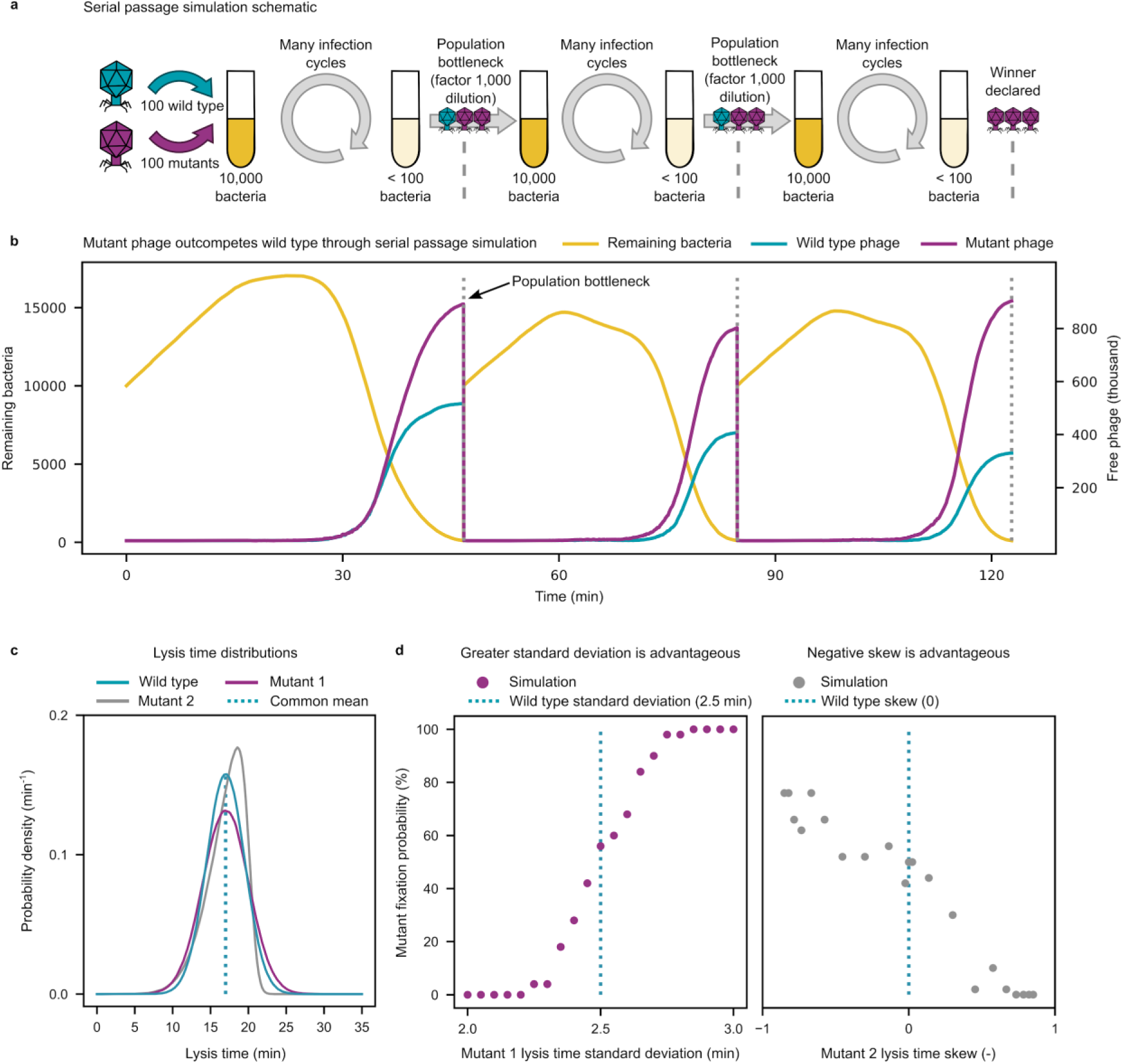
Simulations predict lysis time variance and skewness impact phage fitness. **a.** A schematic explaining the serial passage simulation, where a wild type and mutant phage with corresponding lysis times drawn from different distributions (Fig. 7c) compete and undergo several rounds of dilution into fresh bacteria (Methods, Supplementary note 19). **b.** Example time series data from the simulations, demonstrating the growth and lysis of the bacteria (yellow line), and the proliferation of the wild type and mutant phage (blue and purple lines respectively). At the population bottleneck, a 1,000 fold dilution of the phage into fresh bacteria is simulated. If, after a bottleneck, one phage accounts for more than 70% of the total phage population, it is declared the winner. **c.** Example lysis time distributions of wild type and mutant phage. Mutant 1 (purple) has the same mean as the wild type (blue), but the standard deviation in lysis time is varied (Fig. 7d, left panel). Mutant 2 (grey) has the same mean and standard deviation as the wild type, but the skew in lysis time is varied (Fig. 7d, right panel). **d.** When competing against a wild type phage, mutants with lysis times drawn from a distribution with equal mean but greater variance are predicted to have a fitness advantage (left panel). Mutants with lysis times draw from a distribution with equal mean and variance to the wild type but with negative skew are also predicted to have a fitness advantage (right panel). Each data point is computed from 25 simulations.

Fig. 7d shows the probability of mutant fixation as a function of standard deviation (mutant 1, left panel) and skew (mutant 2, right panel) determined over 25 independent simulations. A proportion of 50% of simulations leading to fixation corresponds to a neutral competition between mutant and wild-type (no fitness advantage). A higher proportion of simulations leading to fixation corresponds to the mutant possessing a competitive advantage and, vice versa, a lower proportion indicates the wild-type possessing a competitive advantage. The results show that a larger standard deviation confers a competitive fitness advantage if mean and skew are the same, while a negative skew in lysis time is advantageous when the mean and standard deviation are the same. Overall, our results clearly indicate that the mean lysis time alone is not sufficient to predict whether a phage can outcompete another, and the higher order moments of the distribution can significantly alter a phage’s competitive advantage. Although here we investigate the effect of variation in lysis time alone, while keeping the other phage life history parameters constant, so as to isolate its specific effect on phage fitness, in reality, these parameters are likely dependent on each other, giving rise to a range of tradeoffs. In recent work, we have explored in more detail how such inter-dependencies in variability can shape phage fitness^56^. Moreover, mutants of phage lambda have been reported to display a broad range of variance in lysis time^57^, supporting the idea that noise in infection dynamics can be shaped by the phage’s genetic makeup and constitute an inheritable trait.

## Discussion

Here, we have reported the first study that quantifies the kinetics of individual steps in the lytic cycle of phage T7 at single-phage-single-cell resolution. Our assay provides a new way to quantify phage-bacteria interactions that is orthogonal to bulk analyses, which provide dynamic but averaged phenotypes, and structural investigation, which assesses variability but is based on static observations. Our approach enables kinetic measurements across many infection events in a precisely controlled environment, while maintaining the individuality of each of them in order to assess variability and correlations across the phenotypes of the phage and the corresponding infected cell.

We find that the major source of variability in lysis time originates from the early stages of infection, from viral DNA entry up to cell growth arrest, while the second part of the infection process, between host take-over and cell lysis, proceeds with remarkable consistency. Analysis of single-cell trajectories revealed five distinct categories of genome entry dynamics. Four of these display rapid DNA translocation, while one proceeds at a slower, nearly constant rate. This slow-entry category is consistent with a process in which the host RNAP alone drives genome internalisation; notably, the rate we observe (∼3.6 kbp min⁻¹ at 37 °C) matches the known speed of *E. coli* RNAP-mediated translocation^37^.

One of the four fast-entry categories aligns with the established three-step genome entry mechanism, in which the first ∼6 kbp of the T7 genome are translocated by the *E. coli* RNAP and the remainder by the much faster T7 RNAP. Two of the remaining fast-entry categories show an initially high rate of genome entry, distinct from this established pattern. Occasionally, we observe stained phage genomes entering cells without subsequent infection, which may represent cases where *E. coli* successfully repels a T7 infection. Therefore, events of these categories could potentially be explained by the presence of the T7 RNAP enzyme remaining in the cell from a previous unstained genome injection and driving rapid genome entry^38^, or even by a new, as yet unknown genome entry mechanism. The final fast-entry category shows injection events where the majority of the genome enters at speeds of up to 60 kbp min^−1^, which could be due to some malfunction of the T7 injection machinery causing uncontrolled genome translocation.

Importantly, our method provides sufficient precision to distinguish between these different categories of dynamics, even when their overall injection durations are similar. The small number of molecules involved in these initial steps (one phage genome and few T7 RNAPs) likely underlies the high level of noise observed in this process^58^. Nevertheless, we cannot exclude that variability in the structural properties of the capsid^59^ and consequent attachment to the host cell receptors might also contribute. Future studies perturbing the genome injection process with antibiotics and genomic modifications, or quantifying the proportion of unsuccessful adsorptions and the intracellular dynamic of the T7 RNAP, will help clarify the mechanistic basis of these distinct entry modes.

Following host takeover, the infection enters a productive phase characterised by rapid synthesis of phage proteins. During this phase, in which capsid proteins and, arguably, viable phage particles are produced within the host cell, we observe a surprisingly large variability in expression kinetics and total production, despite the robustness in the timing of the events. The strong correlation between the total phage protein production and the size of the growth-arrested host suggests that this variability may originate in physiological differences across infected cells. Indeed, using our mathematical model, we find that the phage protein production rate strongly depends on the translational resources of the cell, which scale with cell size, providing a strong mechanistic link between host cell physiology and phage burst size.

Following the productive phase, the infection culminates in host-cell lysis, releasing the newly assembled virions. We found that material loss from T7 infected cells begins approximately 6.6 s before the complete breakdown of the cell envelope - a period we term the perforation phase. Three non-exclusive mechanisms could explain this short delay between the onset of perforation and complete lysis, each primarily affecting a different layer of the cell envelope. One possibility is that aggregations of holins^60^ form initial ruptures in the inner membrane^61^ which then grow, progressively increasing envelope permeability until envelope integrity fails. Alternatively, large inner membrane lesions may already be present when perforation becomes visible, and the ensuing delay reflects the time required for endolysin^62^ (gp3.5), accumulated earlier in the cytoplasm, to degrade the peptidoglycan. A third possibility is that the delay represents the time required for spanins^63^ (gp18.5 and gp18.7) to create a large enough hole to trigger a widespread collapse of the structurally important^7^ outer membrane. Our data support one or both of the latter two scenarios. The abrupt drop in cell contrast (seen from the rise in phase intensity) at the onset of perforation suggests that membrane rupture occurs suddenly. While it is unclear if this initial surge corresponds to a rupture of the inner or outer membrane, we favour the hypothesis of it reflecting the formation of an outer membrane perforation. The ensuing ∼6 s interval likely represents the enzymatic degradation and mechanical weakening of the cell wall or outer membrane, or both, preceding the catastrophic failure of the cell envelope.

We therefore interpret lysis as comprising three coordinated phases: an initial rupture phase (inner membrane breach by holins), a fracture phase (cell wall degradation by endolysins), and a perforation phase (outer membrane breach by spanins). Further studies are necessary to establish if these phases are entirely sequential, or if the second and third phases occur in tandem. It is likely that at some critical threshold of overall cell envelope strength, crack propagation^64^ will take over and govern the rapid breakdown (Supplementary note 16) of the cell envelope. Further work using fluorescently-tagged lysis proteins could help to elucidate the mechanisms and kinetics of T7-induced lysis.

The ordered sequence of infection events, from genome entry to cell lysis, nonetheless exhibits measurable variability at each step. Accurate quantification of the sources of their relative variability and their relative correlations across infection events is not only important to understand the underlying molecular mechanisms controlling phage infection outcomes, but can also have significant evolutionary consequences. Our simulation results clearly show that mean phenotypic values, such as average lysis time, are insufficient to predict the fitness advantage of a phage population and that higher moments of the distribution can have a significant impact. These findings open the intriguing and currently under-explored possibility that variability in phage infection phenotype could be under strong selective pressure^65,66^, raising fascinating questions regarding how evolution shapes it in different scenarios.

Throughout this work, we have focused on T7-*E. coli* to benchmark the assay using a well-studied model system. In particular, since T7 encodes its own DNA and RNA polymerases, it is likely less affected by the host’s intrinsic variability, which simplifies the modelling assumptions for its gene expression dynamics. We expect our experimental approach, however, to be readily applicable to any natural, evolved, or engineered phage. The high-throughput and scalable nature of the platform can be harnessed for multiplexity, to benchmark a variety of sequence variants of phages against specific target bacteria, or multiple mutants of a target bacteria against a particular phage. Precise characterisation of properties associated with infection steps (such as adsorption, production, and lysis) can generate a multi-phenotypic profile for each phage-bacteria pair, enabling detailed analysis of mechanisms underlying response, resistance, and phage-bacteria co-evolution. Similarly, a collection of natural or engineered phages can be evaluated for their efficacy in eradicating a target strain, with implications for medical or biotechnological applications. In summary, this assay promises to open up new avenues for the systems analysis of phage-bacteria interactions and their practical applications.

## Online Methods

### Bacterial strains and growth conditions

The following bacterial strains were used in this study.

**Table 1:**
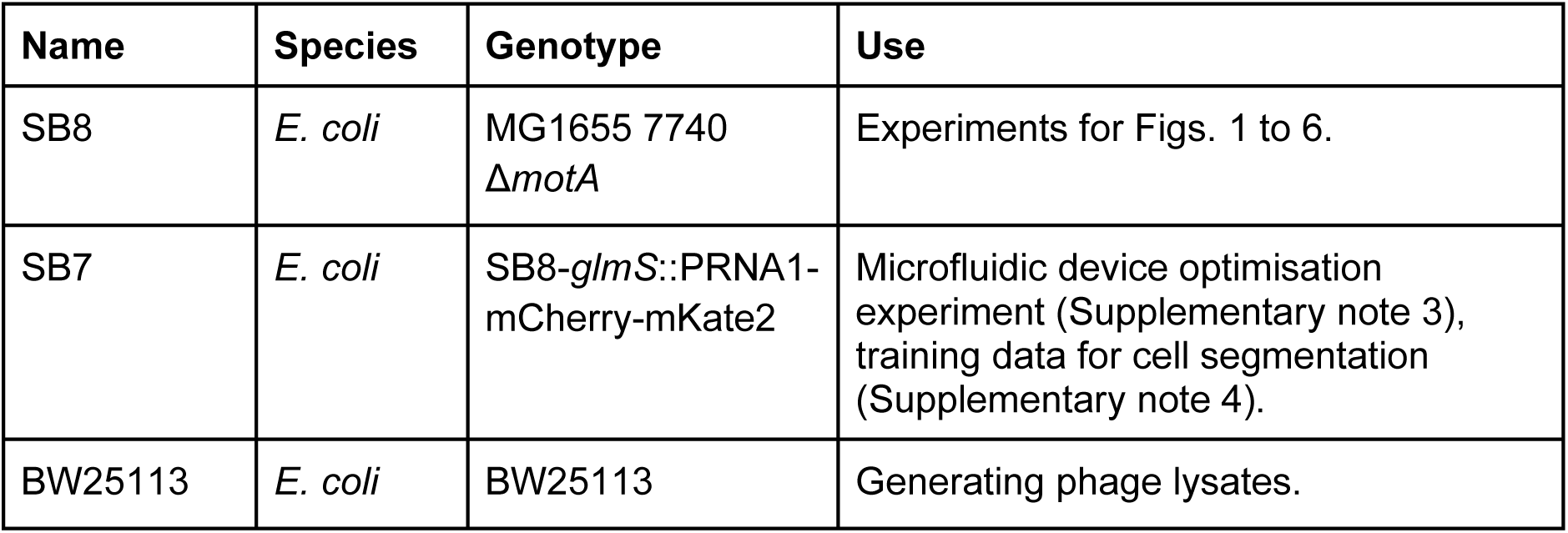
List of *E. coli* strains used in this study.

The motility knockout in SB8 and SB7 prevents cells leaving the trenches of the microfluidic device. Prior to experiments in the microfluidic device, cells were grown overnight in a shaking incubator at 250 rpm and 37 °C in LB Miller (Invitrogen) containing 0.8 g L^−1^ of pluronic F-108 (Sigma-Aldrich, 542342). Cultures were started directly from a frozen stock to maintain a consistent genetic diversity across the cells used in experiments across different days. The LB Miller was sterilised by autoclaving. The pluronic F-108 was first prepared as a 100 g L^−1^ solution and filter sterilised, and then diluted 0.8 % v/v into the LB Miller.

### Phage lysate preparation

*E. coli* BW25113 strain cells were grown overnight in a shaking incubator at 250 rpm and 37 °C in LB Miller (Invitrogen). 500 μL of the overnight liquid culture was used to inoculate 20 mL volume of LB Miller and left to grow in a shaking incubator at 37 °C for 1 h 40 min. Once the culture reached OD 0.6-0.7, 500 μL of stock phage lysate was added and left in the shaking incubator for 7 min. The phage-inoculated cells were centrifuged in a pre-chilled (4 °C) centrifuge at 5000 rpm for 5 min. The supernatant was discarded and the pelleted cells were resuspended in 2 mL of fresh LB Miller. The resuspended culture was left in the shaking incubator for 1 h at 37 °C for the infected cells to fully lyse. The lysate was transferred into 1.5 mL Eppendorf tubes and centrifuged at 14000 rpm for 10 min. The resulting supernatant was passed through 0.22 μm filters to remove any traces of cell debris and unlysed cells.

### PFU estimation

The number of plaque forming units (PFU) in the filtered lysate was estimated using a plaque assay. For this, serial dilutions of the filtered lysate were set up ranging from dilution factor 10^5^ to 10^8^. 20 μL of each diluted lysate was mixed with 100 μL of overnight BW25113 cells in 5 mL of 0.7% agar LB, kept at 50 °C. The mixture was briefly vortexed and poured as a thin layer of agar on 9 cm-diameter plates and incubated at 37 °C for 4 h. The formed plaques were counted and the number divided by 20 to estimate the number of PFU per μL of each dilution factor. Measurements across three dilution factors were used to estimate the concentration of PFUs per μL of the filtered lysate. The titres of lysates obtained using the above method are listed in Table 2. Filtered lysate was used either directly or stained as per protocol below. Note that lysate titres listed in Table 2 are diluted into growth media for microfluidic experiments.

### Genetic engineering of phage: construction of T7*

Transgenic T7 strain T7* was created by assembling PCR-cloned fragments of WT T7 genome along with the fragment encoding T7 phi10 promoter followed by *E. coli* codon-optimised mVenus NB (SYFP2) into a circular plasmid. This circularised transgenic genome was then electroporated into BW25113 cells to produce the transgenic phage lysate. Virions from individual plaques were isolated and sequenced to establish the isogenic strain of T7*. A full description of the PCR protocol to clone the required fragments, the Gibson Assembly of the transgenic genome, the electroporation protocol and the isolation of transgenic strains is available in Supplementary note 1.

### Staining the DNA of the phage genome

The phage lysate was treated with DNAse I-XT to remove any residual bacterial DNA and then stained with SYTOX Orange at the final concentration of 25 uM. Details of the DNAse I-XT treatment and SYTOX Orange staining protocol are available in Supplementary note 2.

### Population level measurement of lysis time

Population level measurements of the mean lysis time of wild type T7 and T7* were carried out according to the ‘one-step growth curve’ or ‘lysis curve’ protocol^67^, a full account of which is available as Supplementary note 18. Measurements were taken in LB using SB8 (Table 1) as the host bacteria. Each phage’s lysis time was measured over three biological replicates, and we find these values to be 14.8 ± 1.3 min for wild type T7, and 20.1 ± 2.0 min for T7* (mean ± 1 standard error of the mean).

**Table 2:** List of T7 strains and typical corresponding lysate PFUs obtained.

| Experiment | Phage strain | Lysate PFU |
| --- | --- | --- |
| Figs. 1, 3, 6 | T7*, SYTOX Orange stained | $1.0 \times 10^7 \mu\text{L}^{-1}$ |
| Fig. 2 | Wild type (WT) T7, SYTOX Orange stained | $4.4 \times 10^6 \mu\text{L}^{-1}$ |
| Fig. 5 | WT T7, unstained | $4.0 \times 10^7 \mu\text{L}^{-1}$ |

### Microfluidic device fabrication

The microfluidic devices were fabricated using soft lithography, by casting a silicone elastomer onto a silicon wafer. We received this wafer as a gift from Dr. Matthew Cabeen of Oklahoma State University. It was fabricated by the Searle Clean Room at the University of Chicago (https://searle-cleanroom.uchicago.edu/) according to the specifications provided by Dr. Cabeen and his colleague, Dr. Jin Park. These specifications were based on the design presented by Norman *et al.*^29^. The silicone elastomer was prepared by mixing polydimethylsiloxane (PDMS) and curing agent from the Sylgard 184 kit (Dow) in a 5:1 ratio and degassing for 30 min in a vacuum chamber. The elastomer was then poured onto the silicon wafer and degassed in a vacuum chamber for a further 1 h. The elastomer was then cured for 1 h at 95 °C. The appropriate devices were then cut out and inlet and outlet holes were punched with a 0.75 mm biopsy punch (WPI). Devices were then cleaned with Scotch Magic tape before being sonicated in isopropanol for 30 min, blow dried with compressed air and then sonicated in distilled water for 20 min. 22×50 mm glass coverslips (Fisherbrand) were sonicated for 20 minutes in 1 M potassium hydroxide, rinsed and then sonicated for 20 min in distilled water before blow drying with compressed air. The devices and coverslips were then dried for 30 min at 95 °C. Devices and coverslips were plasma bonded using a Diener Electronic Zepto plasma cleaner, by first pulling a vacuum to 0.1 mbar, and then powering on the plasma generator at 35% and admitting atmospheric air to a chamber pressure of 0.7 mbar for 2 min. The device and coverslip were then bonded and heated on a hotplate at 95 °C for 5 min, before transferring to an oven at 95 °C for 1 h to produce the finished microfluidic devices.

### Single-cell infection assay

On the day of the experiment, sterile growth media containing LB Miller (Invitrogen) with 0.8 g L^−1^ pluronic F-108 (Sigma-Aldrich) was loaded into a syringe. The pluronic is added as a surfactant to prevent cell clumping in the overnight culture, to improve cell loading, and to prevent cells clumping at the outlet of trenches. It is added to the media at sub-inhibitory concentrations^22^. The lane of the microfluidic device to be loaded was first passivated by adding the above described growth media into the lane with a gel loading tip, and allowing it to rest for 10 min. A 1 mL volume of the cells grown overnight were transferred into a 1.5 mL tube and spun gently at 1000 g for 3 min to sediment the cells. The supernatant was poured away and the cells resuspended in the residual volume. A small volume of this dense cell culture was then pushed into the passivated lane using a gel loading tip and left to rest for 10 min. During this time, the small stationary phase cells will diffuse into the trenches.

While the cells are diffusing into the trenches, growth media from the syringe is pushed through a silicone tubing path to purge the tubing of air. The tubing has a forked path, and the flow is directed down a given fork using a 3-way solenoid pinch valve (Cole Parmer). One fork supplies the microfluidic device with growth media, while the other leads directly to the waste bottle.

The media flow is then connected to the lane of the device containing the cells using 0.83 mm outer diameter needles. The outlet flow goes to a waste bottle. Inlet flow from the media syringe is driven by a syringe pump, and initial flow is set to 100 μL min^−1^ for 10 min to clear excess cells from the lane. Media flow is then reduced to 5 μL min^−1^. Following this, 365 nm illumination light is shone onto the inlet of the device such that each part of the inlet receives at least 7 min of illumination. This kills any cells not removed by the high flow rate, and helps to prevent biofilm formation in the device inlet.

The cells are grown in the device for a minimum period of 3 h from the introduction of fresh growth media into the lane, to allow the cells to reach a steady state, exponential growth phase. After this wake up period, the media is switched to media containing phage to begin the infection imaging. A typical phage media composition is described in Table 3, which would result in a final phage titre of 10^6^ PFU μL^−1^. Note that the phage lysate is also washed and resuspended in LB Miller (Supplementary note 2), so the exact lysate volume used is unlikely to significantly change the nutritional composition of the media.

**Table 3:** Phage treatment media composition.

| Component | Volume ($\mu\text{L}$) | Volume fraction (-) | Concentration in media |
| --- | --- | --- | --- |
| LB Miller | 4460 | 0.892 | N/A |
| Pluronic F-108, 100 g $\text{L}^{-1}$ | 40 | 0.008 | 0.8 g $\text{L}^{-1}$ pluronic F-108 |
| Phage lysate, PFU $10^7 \mu\text{L}^{-1}$ | 500 | 0.100 | $10^6$ PFU $\mu\text{L}^{-1}$ |
| **Total** | 5000 | 1.000 | N/A |

The phage titre must be sufficiently high to ensure at least some infections occur in each given trench, but the exact titre is unimportant in the ranges used, as we operate at very low multiplicity of infection in order to ensure that all first round infections result from just one phage binding to a cell. The phage titres used in each experiment are listed in Supplementary note 20.

To change the media to phage media, the solenoid pinch valve is activated to block flow to the microfluidic device. The flow to the microfluidic device is blocked for a maximum of 10 min. While it is blocked, the growth media syringe is changed to a syringe containing growth media with phage, and then the tubing is flushed with the phage media at high flow rates, such that the growth media without phage is completely cleared from the tubing. Then, the flow rate is returned to 5 μL min^−1^ and the flow is switched to introduce phage media to the cells. Purging the initial section of tubing with phage media reduces the time between the switch and the phages reaching the cells, without having to expose the cells to high flow rates which could cause mechanical stress. Additionally, it purges bubbles which can sometimes be introduced when the syringes are switched. Once the media is switched, time-resolved image acquisition begins.

The experiments used for Fig. 2 were conducted similarly, with some differences outlined in Supplementary note 21.

### Time-resolved image acquisition

Images were acquired using a Nikon Eclipse Ti2 inverted microscope with a Hamamatsu C14440-20UP camera. The microscope has an automated stage and a perfect focus system, which automatically maintains focus over time. The microscope contains two multiband filter cubes, each of which contains a multi-bandpass dichroic mirror and corresponding multiband excitation and emission filters. There is an additional emission filter which can be quickly switched to select the correct emission wavelength band. Together the multiband cubes and the emission filter wheel allow for fast imaging in multiple colour channels. All captured images are initially saved using Nikon’s ND2 file format. For the experiments in Figs. 1, 3 and 6, imaging began before the phage media reached the cells, and continued at a regular frequency for the duration of the experiments. The microscope settings used for each channel are listed in Table 4 below.

**Table 4:**
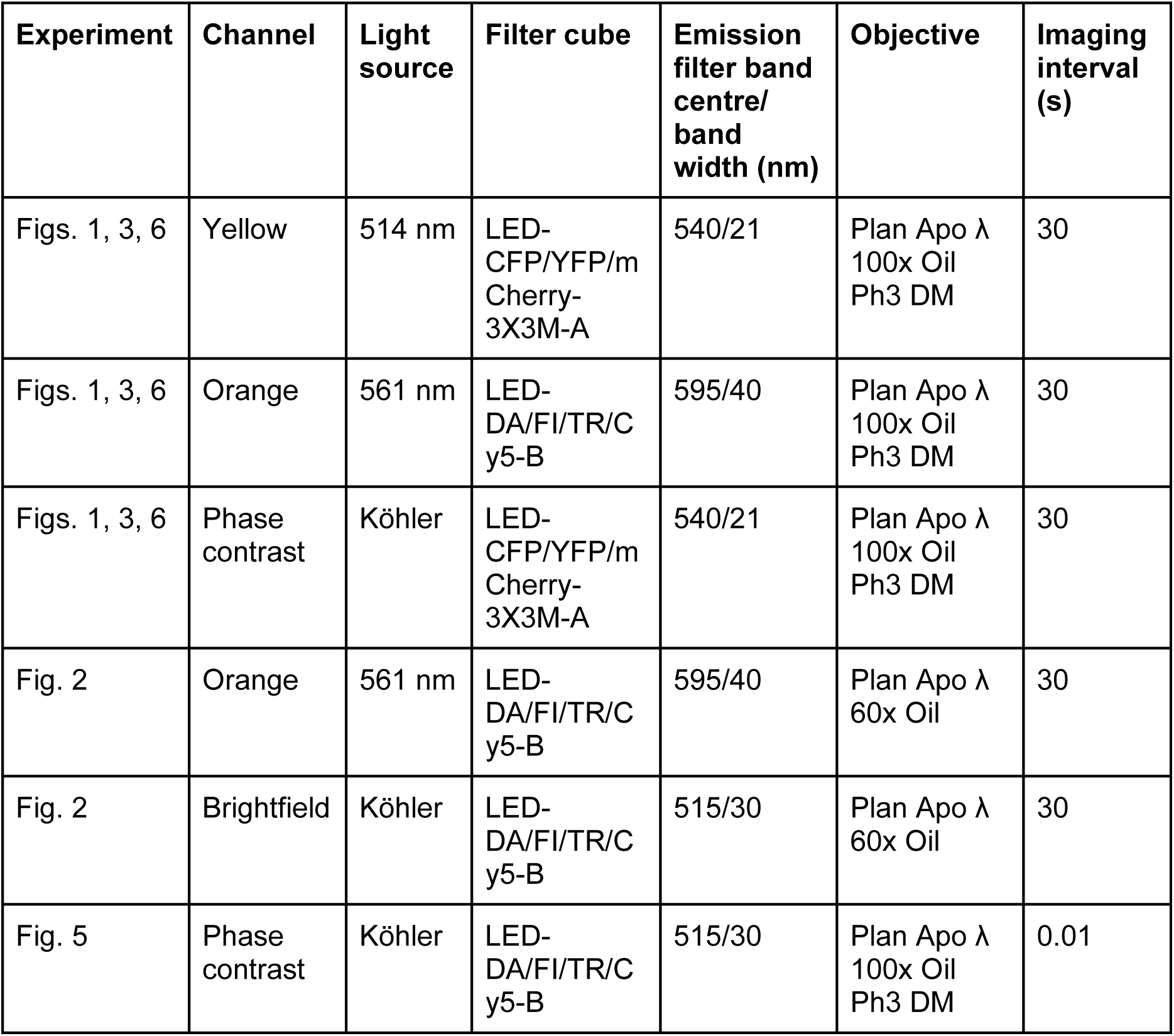
Imaging settings for different experiments used in this paper.

We have used high-speed timelapse imaging (100 frames s^−1^) to capture the events preceding the lysis, as our initial attempts using 1 frame s^−1^ imaging revealed that the structural changes occurring during lysis unfold on sub-second timescales. The high frame rate, while conducive to observing rapid dynamics, renders fluorescence imaging unsuitable due to photobleaching and potential phototoxicity. Nonetheless, phase-contrast imaging is sufficient to gain a detailed insight into the material loss from the cell to its surroundings in the fleeting moments preceding lysis (Fig. 5). A short exposure time (3 ms) and a small region of interest (ROI) around individual trenches filled with cells close to lysis (Supplementary note 22) enabled us to achieve an imaging interval of 10.2 ms (Supplementary movie 4).

### Image preprocessing

All captured images were pre-processed before feature extraction. First, individual frames from the Nikon ND2 format were extracted and saved as PNG files using custom Python code which makes use of the nd2 module (https://pypi.org/project/nd2/). Using custom Python code (https://github.com/georgeoshardo/PyMMM), these frames were then registered to correct for any stage drift and rotated to ensure the trenches were vertical in the images. We then use automated methods to find the position of each trench in the images and crop out the trenches for further processing, as described below.

### Cell segmentation

The phase contrast images of cells in the extracted trench images from our experiments in Figs. 1, 3 and 6 were segmented using a custom trained Omnipose machine learning model^68^. The model was trained on images (taken on a different day) of SB7 *E. coli* (Table 1) growing in our device where both fluorescence and phase contrast images were acquired using the same objective as for the experiments. These images will be referred to as training images, and are separate from the experiment image data.

To train the model, our approach was to first train an Omnipose model to segment fluorescence images. To generate a high volume of training data and corresponding ground truth masks for fluorescence images of cells in the mother machine, we use a virtual microscopy platform called SyMBac^31^. Using this fluorescence model, we segmented fluorescence images of cells and generated cell masks for the fluorescence channel of the training images.

The cell masks for the fluorescence channel were then checked against the phase contrast training images, and pairs of fluorescence masks and phase contrast training images which matched well were manually curated into a training data set. This training data was used to train an Omnipose model for the segmentation of phase contrast images. The phase contrast model was then used to generate cell masks for the experiment image data. This pipeline is further described in Supplementary note 4.

### Feature extraction

Basic cell properties, such as cell position, length, area, and YFP intensity, were extracted from regions of the images corresponding to the cell masks produced by segmentation. The cell properties were extracted using custom Python code (https://github.com/CharlieW313/MM_regionprops) utilising the scikit-image regionprops function^69^. For the data in Fig. 3, further properties are calculated from the basic cell properties. Mathematical descriptions are found in Supplementary note 23.

### Single-cell lineage tracking

The single-cell growth and lysis traces were tracked over time using features extracted at each frame, including cell position, area, orientation, and Zernike moments. This process was done using a custom Python script (https://github.com/erezli/MMLineageTracking). The algorithm predicts many potential states of these features for each cell at subsequent time steps. It then finds the best match to the feature states in the following frame to determine the tracking outcome. The results are stored in tree-structured Python objects containing detailed cell properties such as YFP mean intensity. The tracking results are manually checked by visualising them in kymographs. Further information about the algorithm can be found in Supplementary note 5.

### Single-phage tracking

To track the injection of the phage genome, we monitored the intensity of the bright spot indicating the phage location over consecutive frames until injection was complete. Spot intensity was measured as the intensity of a fraction of the brightest pixels in a rectangular box centred on the spot, with a control box alongside for background subtraction. Genome injection duration was estimated as the time between adsorption and the spot intensity falling to a specified threshold. Further details concerning the tracking of individual phage spots and the calculation of genome injection time for Figs. 1, 2 and 6 are presented in Supplementary note 8.

### Analysis of capsid production data

The fluorescence intensity in the yellow channel was analysed to determine the start of capsid production, as the time point where the signal from the YFP reporter of capsid production increases above the baseline. We subtract the background intensity from the raw total intensity of the YFP reporter to give a total intensity, *I*(*t*), as described in Supplementary note 22. The production start time, *t*_3_, is calculated as the first time point when the total intensity reaches a threshold value (chosen to be 1420 AU based on inspection of the intensity time-series), and then subsequently remains above that threshold for a total of four consecutive time points. This start is later adjusted to *t*_3_ ∗ by fitting the model, as explained in Supplementary note 15.

### Analysis of perforation and lysis

For the high time resolution imaging of phage induced lysis, a machine learning based approach for cell segmentation was unsuitable. This was because we wished to monitor the phase contrast intensity of the cell before, during and after lysis, so any attempt to segment the cells using features of the image would begin to fail as those features markedly changed through the lysis process. We therefore used a hand-drawn manual segmentation of a static region at the location of each cell in Fiji (ImageJ)^70^, as the cells did not move significantly over the short timescales of lysis. The mean phase contrast pixel intensity of this region, *i*_*PC*_ (*t*), was then measured in each frame. By translating the static region by 1.43 μm to the left and right of the cell (along the short axis of the trench), the phase contrast intensities of the regions adjacent to the cell were also measured. By further translating the region on the right of the cell an additional 0.72 μm to the right and 4.29 μm along the long axis of the trench towards its closed end, a region in the side trench far away from the lysis was used as a control region for the phase contrast intensity.

To determine the start of perforation (*t*_4_), the mean and standard deviation of the phase contrast intensity over 200 time points (a window ending a few seconds prior to the perforation start) were computed. The perforation start was declared when the phase contrast intensity first exceeds this calculated mean plus three standard deviations, for a minimum of five consecutive time points. The start of lysis (*t*_5_), indicated by a purple arrow in Fig. 5d, was determined as the point where the derivative of phase contrast intensity increases above the mean plus three standard deviations of the phase contrast intensity derivative calculated over a 1 s window (between 1.5 s and 0.5 s before the point where the phase contrast intensity derivative reaches a maximum) during perforation. We refer to the time interval between these two timepoints (*t*_5_ − *t*_4_) as the perforation duration.

### Serial passage simulations

The simulation extends infection kinetic ODEs^52^ to a stochastic, agent-based model (for a full description of the implementation see Supplementary note 19). The simulation begins with 2 pools of 100 phage and 10,000 susceptible bacteria in a simulated well-mixed volume of 10^−5^ ml. All bacteria begin the simulation in the ‘uninfected’ state, at random points in their cell cycle. In each simulation time-step, ‘bacterial growth’, ‘adsorption’, ‘infection’, ‘lysis’, and ‘decay’ substeps occur.

Once the number of cells has dropped below 100, we simulate a ‘bottleneck’: a 1,000 fold dilution of the phage and remaining cells, and addition of 10,000 new susceptible bacteria. The simulation continues, with bottlenecks occurring every time the bacterial population falls below 100, until either one phage pool outnumbers the other 70:30 and is declared the winner, or until a preset timeout, at which point we declare a tie.

## Acknowledgements

We are thankful to the members of the Bakshi Lab and Fusco lab for their support and feedback on this work.

## Declarations

### Authors’ contributions

S.B. and D.F conceived the study and were in charge of the overall direction and planning. C.W., T.Y., A.S., M.H., D.F., and S.B. designed the experiments and simulations. C.W. and R.L. carried out the microfluidics microscopy experiments, A.S. and M.H. carried out the bulk experiments, T.Y. carried out the genetic engineering of phages, and T.Y. and A.K.M. carried out the phage staining. C.W., R.L., G.H., and S.B designed the data analysis pipeline and carried out the data analysis. M.H., A.S., and D.F. carried out the simulations and associated analysis. D.F. developed the mathematical model and associated analysis. C.W., D.F., and S.B., lead the manuscript writing. R.M. contributed experimental material and methods. All the other authors provided critical feedback and contributed to the manuscript.

### Data availability

Data for this paper is available from the Zenodo repository associated with this paper, which can be found at 10.5281/zenodo.13227935.

### Code availability

Microscopy images were registered using the custom-built python script: https://github.com/georgeoshardo/PyMMM. Registered images were segmented using an Omnipose model trained with synthetic image data generated using the SyMBac pipeline: https://github.com/georgeoshardo/SyMBac. Single-cell features from the segmented images were extracted using https://github.com/CharlieW313/MM_regionprop. The custom-built Python script for tracking individual cell lineages in time series data is available at: https://github.com/erezli/MMLineageTracking. Python scripts for phage diffusion simulations, phage competition simulations, and Matlab script for fitting gene-expression data to mathematical models are available at: https://github.com/FuscoLab/single-cell-phage-tracking

## Supplementary Information

### Single-cell imaging of the lytic phage life cycle in bacteria

#### Supplementary note 1 Integration of capsid expression fluorescent reporter into the T7 genome

To create T7*, the phi10:mVenusNB fragment was inserted directly after the stop codon of T7 gene 10A. To this end, wild type (WT) T7 genomic DNA was extracted from a fresh lysate preparation (Methods, Phage lysate preparation) using Norgen Genomic DNA extraction Kit (Norgen, SKU 24770), following manufacturer’s instructions. The WT genomic DNA was used as a template for Q5® High-Fidelity 2X Master Mix (NEB, M0492L) PCR reactions to clone three fragments of the T7 genome, as well as the phi10:mVenusNB insert using the oligos listed in Supplementary table 1. The template for the *E. coli* codon-optimised mVenusNB insert was ordered as a gene block from IDT. The circularisation fragment was ordered as a pair of oligos from NEB and was annealed prior to their use in the DNA assembly reaction.

**Supplementary table 1:**
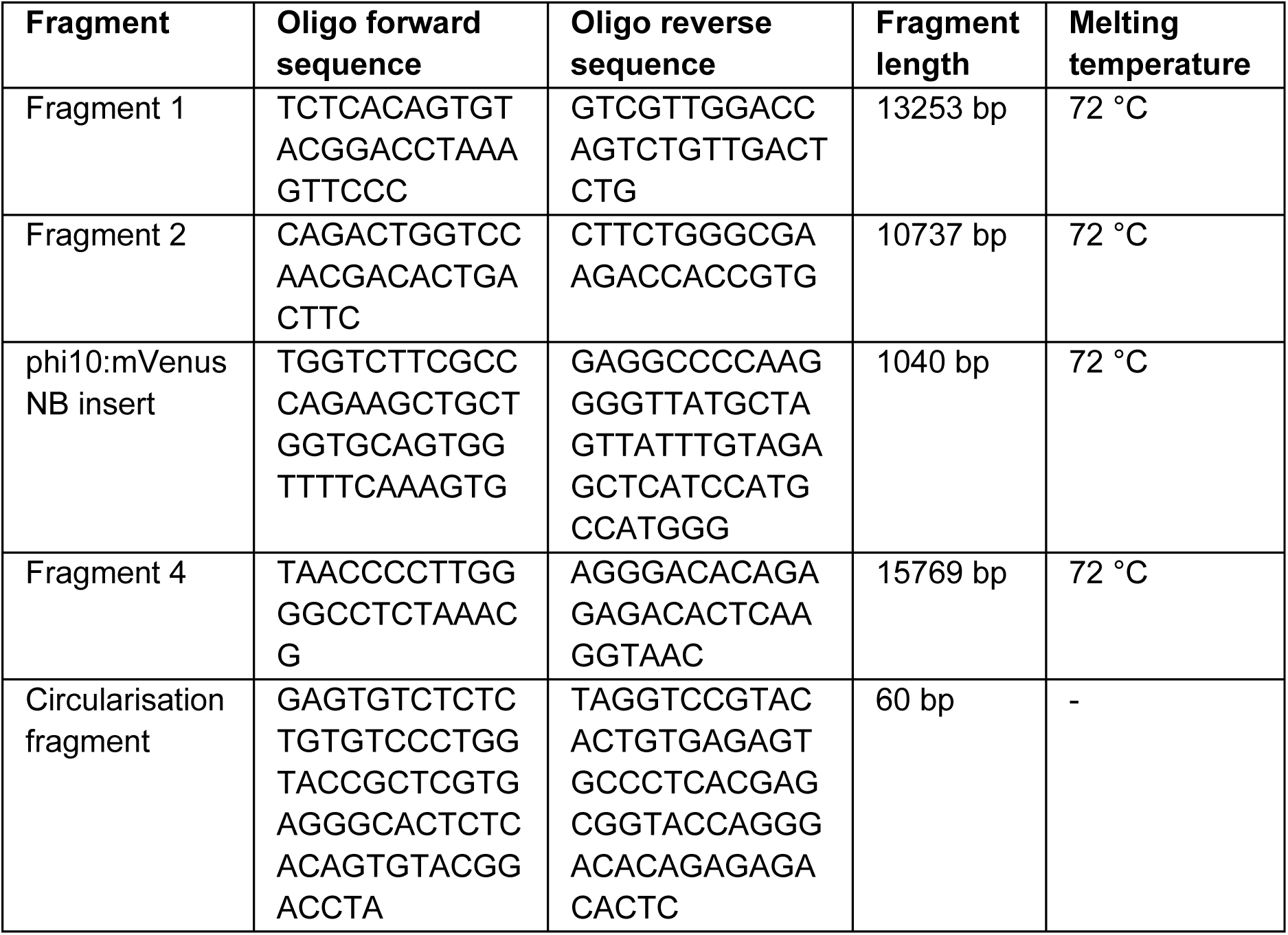
List of oligo sequences.

The Q5 Master Mix was supplemented with 0.5 μL of 10 μM DNTP’s solution and the extension time was set to 13 min. The cloned fragments were purified using the Monarch® PCR & DNA Cleanup Kit (5 μg) (NEB, T1030L) and the DNA concentration was measured using Qubit 4 Fluorometer (Invitrogen). The fragments were assembled in a NEBuilder® HiFi DNA Assembly Master Mix (NEB, E2621L), as per manufacturer’s recommended ratios and incubated at 50 °C for 1.5 h. 10 μL of the reaction product was used to transform an aliquot of electro-competent cells using the MicroPulser Electroporator (BioRad). The cells were pulsed 3 times to ensure the integration of the circularised phage genome within the cells (although alternative methods for transformation were also tested and found to be similarly efficient, Supplementary table 2). Overall, we found that the exact transformation efficiency could vary over orders of magnitude from batch to batch, as a function of number of competent cells and amount of genomic DNA. The latter was tested more systematically using wild type T7 genomic DNA, with order of magnitude of number of plaques reported in Supplementary table 3.

**Supplementary table 2:** Efficiency of transformation of phage genome into *E. coli*.

| Transformation method | Number of plaques per 20 uL of transformed lysate at 10<sup>-4</sup> dilution |
| --- | --- |
| Electro-competent cells with linear synthetic genome | 107 |
| Electro-competent cells with circularised synthetic genome | 83 |
| Chemically-competent cells with linear synthetic genome | 18 |
| Chemically-competent cells with circularised synthetic genome | 65 |

**Supplementary table 3:** Efficiency of transformation as a function of phage genome amount.

| *E. coli* strain | Amount of transformant linear phage gDNA |  |  |  |
| --- | --- | --- | --- | --- |
|  | 300ng | 100ng | 50ng | 30ng |
| BW25113 (chemically-competent) | 100 PFUs | 100 PFUs | 0 PFUs | 0 PFUs |
| DH5a (chemically-competent) | 100 PFUs | 0 PFUs | 0 PFUs | 0 PFUs |

The electroporated cells were immediately supplemented with 1 mL of room temperature LB Miller and left in a shaking incubator at 37 °C for 1 h. The resulting lysate was diluted serially up to dilution factor of 10^−4^ and all dilutions were used in a plaque assay. The plaques were screened under a microscope and YFP-expressing plaques were individually cored out and dissolved in 50 μL of fresh LB. These phage suspensions were used to prepare concentrated lysates, from which genomic DNA was extracted. DNA extracted from candidate plaques was fully sequenced by Plasmidsaurus to confirm correct scar-free integration of the phi10:mVenusNB insert and absence of any mutations from the template WT T7 genome.

#### Supplementary note 2 Staining of T7 genome with SYTOX orange

The genomic DNA of the T7 phage was stained with the DNA binding dye SYTOX Orange (Invitrogen, S11368). To reduce the bacterial DNA background, the concentrated phage lysate was incubated at 37 °C with DNase I-XT (NEB M0570L), for 1 h. Such treatment did not significantly reduce the titre of PFUs (Fig. S1), but significantly reduced the background fluorescence from dyes bound to DNA fragments from lysed cells. The treated lysate was centrifuged across a 100 kDa (Amicon® Ultra, UFC5100) filter and rinsed across the filter membrane with fresh LB five times. The concentrated stained phages were resuspended in 1 mL of fresh LB and then stained with SYTOX Orange at a final concentration of 25 μM and left overnight at 4 °C. Excess dye was removed by centrifuging the stained lysate across a 100 kDa (Amicon® Ultra, UFC5100) filter and rinsing the dye away across the filter membrane with fresh LB five times. The concentrated stained phages were resuspended in 1 mL of fresh LB and stored at 4 °C.

**Fig. S1:**
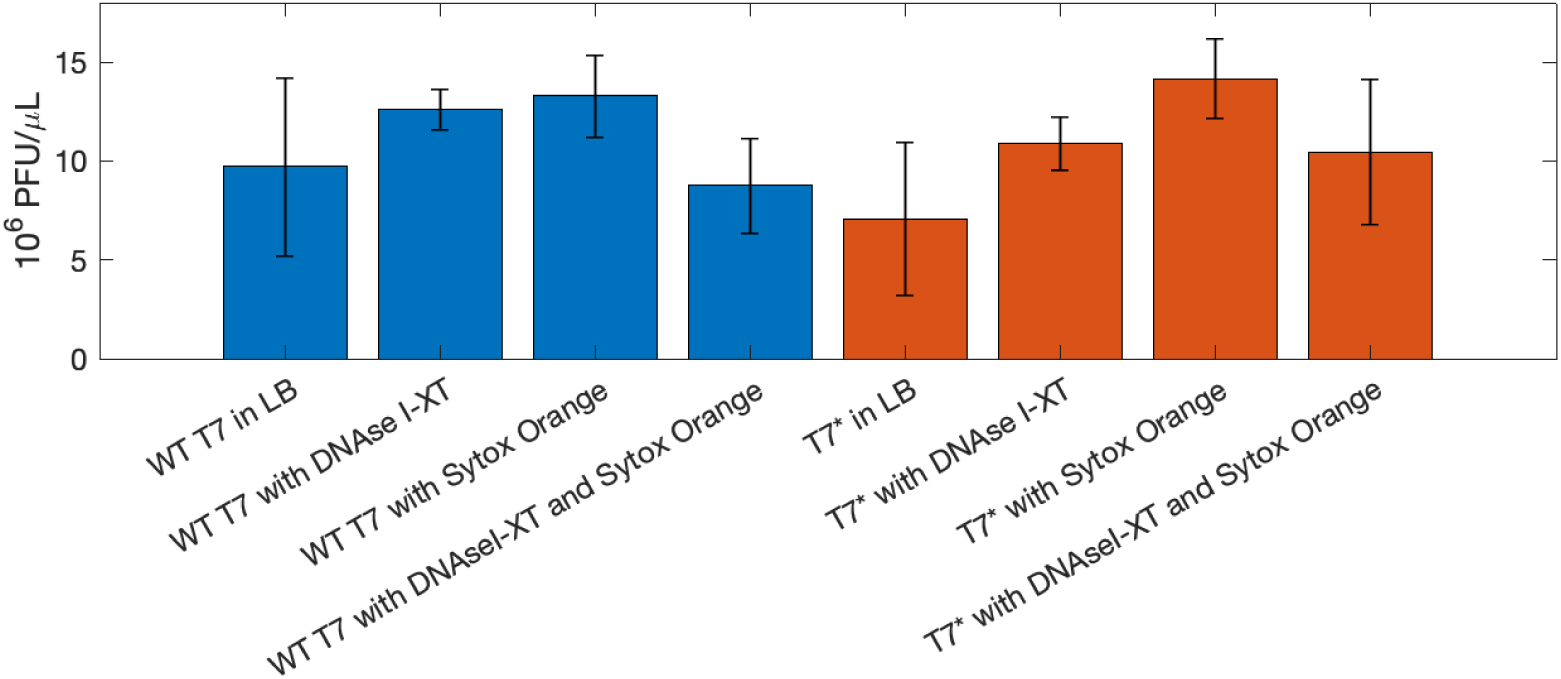
Effect of DNase I-XT and SYTOX Orange on the viability of T7 virions. The WT T7 (blue) and T7* (red) lysates were subjected to DNAse I-XT treatment, SYTOX Orange treatment and both treatments. Plaque assay experiments were performed to compare PFU concentration in the treated samples and the untreated controls (LB). No significant reduction of PFUs was observed after any treatment.

Slight variations of the lysate staining method outlined above were used when preparing phage samples for imaging. These variations, which compared to the optimised method outlined above included using a 5x lower concentration of the dye and fewer filtration steps, were made in attempts to improve the signal-to-noise ratio during imaging. Fig. S10 (Supplementary note 11) shows results from three experiments on T7*, where these variations were included, and there are no significant differences in the measured single-cell lysis times if outliers are excluded (as described in Fig. S10). This is consistent with the results shown in Fig. S1, where we find that SYTOX Orange staining does not affect PFU counts in stained lysates.

#### Supplementary note 3 Microfluidic device design optimisation

To investigate the effect of trench design and presence of side trenches on the access of phage to the cells, we carried out diffusion simulations and experimental infection assays.

We first set up a diffusion simulation with the aim of understanding how bacteriophage particles travel along a trench containing cells, and how likely they are to adsorb to cells in a given position. We simulate a bacteriophage virion as a random-walking particle, initialised at the end of a trench open to the feeding lane. The virion was allowed to take random steps in three dimensions, constrained to the geometry of the trench. The walls of the trench, as well as the closed end, were treated as hard wall boundary conditions, in that if the virion attempted to step into them it would bounce off. The cross section of the trench (Fig. S2c) was taken to be constant along its length, with the cell membrane acting as a stochastically “sticky” boundary condition: if the virion attempted to step into the cell it had a chance to adsorb, calculated from its bulk adsorption rate, or would otherwise bounce off. Virions that stepped backward into the feeding lane were considered to have washed out, and their random walk terminated.

**Supplementary table 4:** Parameters used in phage diffusion simulations.

| ***Simulation Parameters*** |  |
| --- | --- |
| Random walk step size ($dx$) | 50 nm |
| Timestep ($dt$) | Calculated from step size and diffusivity such that random walk matches diffusive behaviour: $dt = dx^2/2D$ |
| Trench length ($Z$) | 70 $\mu\text{m}$ |
| Trench cross section | Central trench 1.4 $\mu\text{m}$ x 1.4 $\mu\text{m}$ in both cases. Where present, side trenches were 0.4 $\mu\text{m}$ high, 2.8 $\mu\text{m}$ wide. See Fig. S2c. |
| Number of simulations ($N$) | 100,000 for each combination of cross section and adsorption probability |
| ***Phage Parameters*** |  |
| Virion diffusivity ($D$) | 4 $\mu\text{m}^2 \text{s}^{-1}$ |
| Virion radius ($R$) | 25 nm |
| Adsorption rate ($\alpha B_0$) | Bulk adsorption rate multiplied by bacteria density. In liquid culture, we expect the concentration of virions to decrease exponentially at this rate.<br>[3, 10, 30, 300, 3000] $\text{min}^{-1}$ |
| Adsorption probability ($a$) | Probability to adsorb on attempting to step into the cell. Calculated from adsorption rate: $a = 2 dt \alpha B_0$ |

For virions that successfully adsorbed to a cell, we recorded their trench penetration distance. This distance was converted to units of cell lengths, assuming 1 cell length = 2 μm, and the number of virions adsorbed “to each cell” was plotted against cell number, 1 being the closest to the feeding lane (Fig. S2a). We find that the presence of side trenches greatly increases the number of phages able to adsorb to cells further from the feeding lane. To examine the robustness of this result to the bacteriophage’s adsorption rate, we ran the simulation using a range of adsorption rates over three orders of magnitude (Supplementary table 4). For each combination of cross-section and adsorption rate, we calculated the characteristic penetration depth: the distance along the trench required for the number of adsorptions per cell to drop by a factor of *e* (Fig. S2b). The results show that the presence of side-trenches doubles the characteristic penetration length consistently across adsorption rates.

To test the effect of side trenches on phage access experimentally, we carried out a parallel infection assay across two trench designs, one with side trenches, and one without (Fig. S2d). The two types of trenches were in separate, isolated flow lanes, but the infection assays were performed simultaneously. The method used was similar to that described previously (Methods). SB7 cells (Table 1) were grown in the trenches of both devices. The growth media used was LB Miller (Invitrogen), with an addition of 0.8% v/v of sterile filtered 100 g L^−1^ pluronic F-108 (Sigma-Aldrich) solution. Cells were first grown in this media for 5.5 h at a flow rate of 5 μL min^−1^ before the media was switched to phage media at the same flow rate. Media containing phage was prepared to be the same as the growth media, but with the addition of 2 × 10^5^ PFU μL^−1^ of T7* phage. The cells were imaged in red fluorescence, yellow fluorescence and phase contrast every 2.5 min during the course of the infection using a Plan Apo λ 40x air Ph3 DM objective with 1.5x post-objective magnification (Methods). Images were segmented using a fluorescence Omnipose model^68^ trained on data generated using SyMBac^31^. For the avoidance of false positives, a cell was determined to be infected if its YFP signal surpassed a threshold value defined as the mean plus seven standard deviations of the YFP intensity distribution measured from over 285,000 observations of uninfected cells in early part of the data. We recorded the time an infected cell was first observed in each given trench, and the time at which a trench became completely cleared by phage lysis.

The results of this comparative infection assay are shown in Fig. S2e. Infections are quickly seen in trenches with sides, with at least one infection being seen in over 50% of trenches within 55.0 min of the first infection being observed. By 97.5 min, at least one infection is observed in over 90% of trenches with sides. By contrast, it takes 285.0 min before over 50% of trenches with no sides see at least one infection, indicating that phage access to cells in trenches without sides is significantly limited. The number of trenches completely cleared by phage lysis is also considerably greater in trenches with sides. 75.0 min after the first infection is observed in each given mother machine lane, all cells had been lysed by phage in 20% of trenches with sides, but in only 3% of trenches without sides. By 175.0 min, all cells had lysed in 90% of trenches with sides, but in only 16% of trenches without sides. These results, in combination with the diffusion simulations, demonstrate that cells are much more readily infected when side trenches are available to facilitate phage transport.

**Fig. S2:**
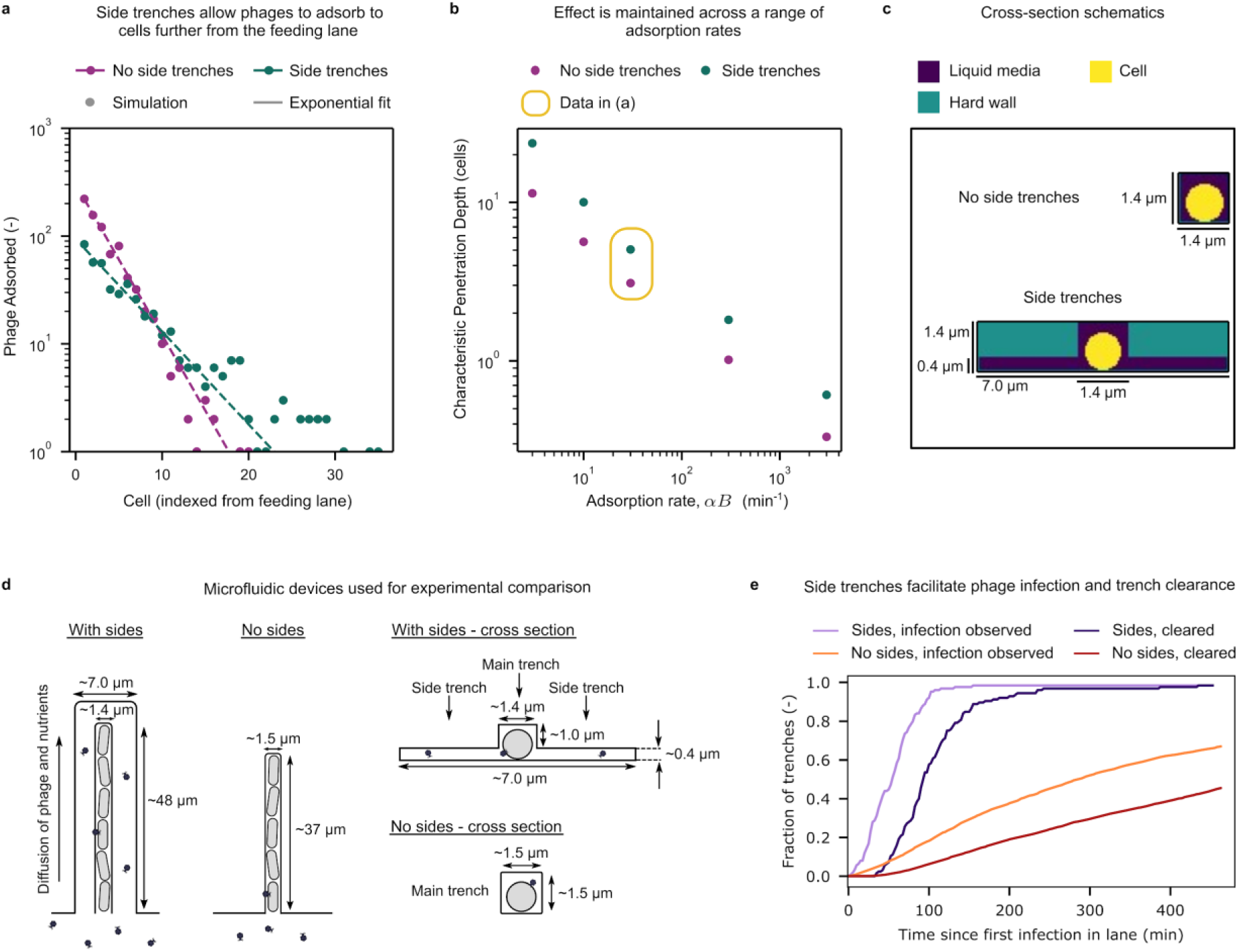
Side trenches increase phage access to cells in simulated and real experiments. **a.** Diffusion simulation results demonstrate that when side trenches are used, phage have an increased tendency to adsorb to cells deeper into the trench. **b.** The effect seen in (**a**) is maintained even when the adsorption rate of phage to host cell is varied. **c.** Device geometry and boundary conditions used in the diffusion simulations in (**a**) and (**b**). **d.** Schematics showing plan and cross-sectional views of the trenches used for the experimental comparison of phage infection in different trench designs. **e.** Experimental results of phage infection in the two trench designs described in (**d**). The light purple and orange lines show the cumulative fraction of the number of trenches of each type where at least one infection was observed. The dark purple and red lines show the cumulative fraction of the number of trenches of each type where all cells were lysed by phage. The time axis is offset by the time of the first observed infection in each respective lane. Both infection and trench clearance proceed much more rapidly in trenches with sides.

#### Supplementary note 4 Two-step machine-learning pipeline for cell image segmentation in phase contrast images

To address the additional features introduced by side trenches of the microfluidic devices in the phase contrast images, we implemented a two-step segmentation pipeline for analysing phase contrast images of SB8 in the mother machine microfluidics device. First, we trained a fluorescence segmentation model using SyMBac-generated synthetic images, which was then applied to segment fluorescence images from SB7 (*E. coli* cells tagged with a constitutively expressed red fluorescent protein marker). Since phase contrast images were captured simultaneously with fluorescence images in the SB7 experiment, we used the resulting fluorescence masks to train a phase contrast segmentation model. This model was then applied to segment phase contrast images of SB8 (lacking any fluorescent marker) from a separate experiment, using the same microfluidics device. The process is illustrated in Fig. S3.

**Fig. S3:**
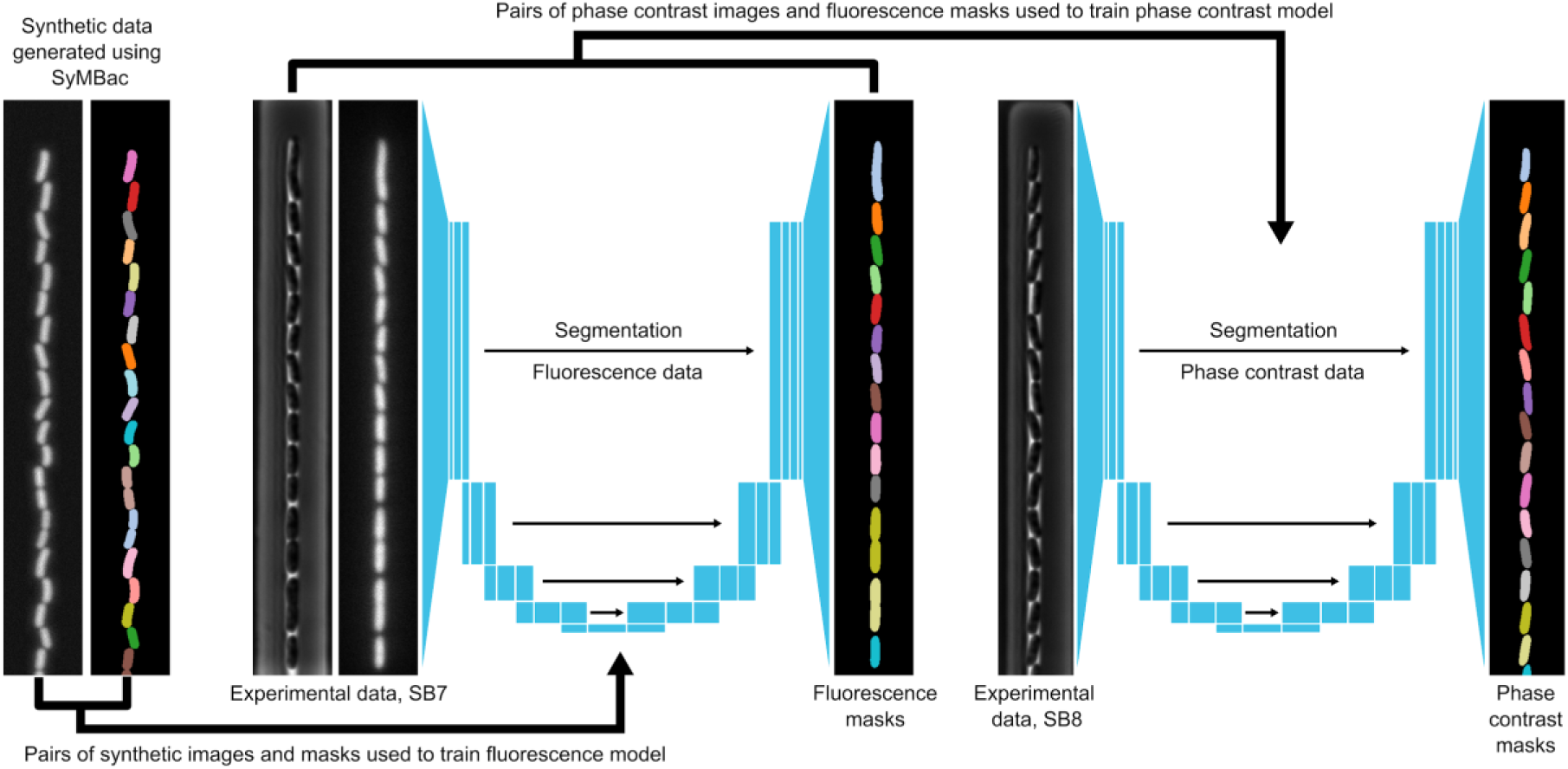
A schematic illustrating the “translational learning” segmentation process. Synthetic image pairs (fluorescence images and ground truth masks) generated by SyMBac^31^ were used to train an Omnipose^68^ fluorescence image segmentation model, which was then used to segment fluorescence image data of SB7 cells in the microfluidic device. The resulting masks were then paired with phase contrast data of SB7, as corresponding ground truth data, to train the Omnipose phase contrast image segmentation model which was used to segment phase contrast data from experiments using SB8.

#### Supplementary note 5 Algorithm for tracking individual cells during growth and lysis

The lineage tracking algorithm (https://github.com/erezli/MMLineageTracking) takes CSV files containing cell properties extracted from segmented channel images using the Scikit-image regionprops function^69^ and Zernike moments extracted from cell masks using the Mahotas package^71^, and uses these as key predictors for tracking the cells. It extracts the statistics of several cell growth parameters (Supplementary table 5) from the mother cells in each trench which do not require tracking.

**Supplementary table 5:** Parameters for lineage tracking.

| Growth parameters | Definitions |
| --- | --- |
| $\lambda$ | Cell elongation rate, such that $L(t + \tau) = L(t)2^{\lambda\tau}$ |
| $\theta_a$ | Adder model parameters (measured mean, variance, and skewness of the added length before a cell divides) |
| $\theta_s$ | Sizer model parameters (measured mean, variance, and skewness of the cell length before a cell divides) |
| Cell parameters | Definitions |
| **L** | Array of all cell lengths in a frame |
| $\delta L$ | Array of added cell lengths since each of their birth in a frame |
| **A** | Array of all cell areas in a frame |
| $\alpha$ | Array of all cell orientations in a frame |
| **P<sub>y</sub>** | Array of all cell vertical positions in a frame |
| **z** | Array of all 1-norm of Zernike moments in a frame |

In each frame, the physical properties of cells are simulated to change under the constraints of the mother machine using the properties estimated from the mother cell track. Each cell is assigned a probability of division, which depends on the cell size regulation model employed, such as the adder and sizer models^72^. The division probability for each cell is calculated as follows:

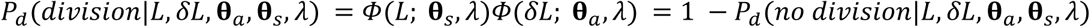

Where ***Φ*** is the skewed normal bivariate cumulative distribution function (CDF) with parameters fitted from the adder and sizer model data. The efficient estimation of the skewed normal CDF is achieved by Owen-T approximation:

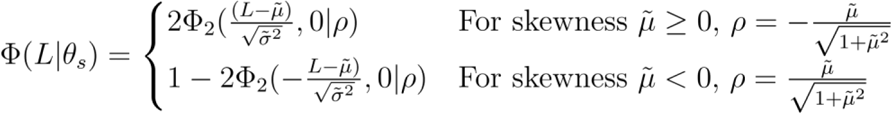

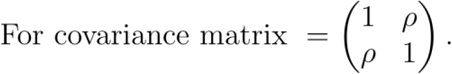

Many potential scenarios are generated using a binary permutation of 0 and 1 in an array, ***div***, indicating division of each cell in each frame. Therefore, the a priori probability of the k-th scenario considering all cells in a frame becomes:

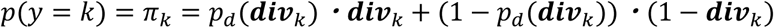

Where *Σ_k_π_k_* = 1.

From the cell parameters and the growth rate, the algorithm generates predictions for cell lengths, areas, positions and Zernike moments across all simulated scenarios by assuming symmetric cell division, effectively creating a high-dimensional feature space. The tracking problem is then regarded as classifying the next frame, *x*, into one of the simulations, *y*. After matching the high-dimensional array in sequential order (allowing skips since cells can lyse) for each simulation, the k-th simulation will return a minimum Euclidean distance *d*_*k*,*min*_. This gives a likelihood probability of each scenario, represented in softmax form:

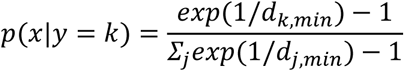

From this, we can classify the next frame into one of the simulations by maximising the posterior:

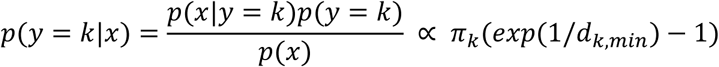

The prior can be given an arbitrary offset to improve robustness, and then the final decision is made by considering the score:

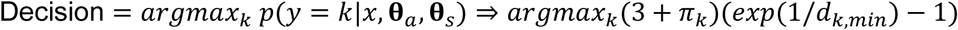

Given that the number of simulations scales exponentially with the number of cells that need to be tracked simultaneously, and the simulation noise increases proportionally to the number of cells (as positional changes accumulate in one direction), we choose to concurrently track only a restricted number of cells and retain the lineage results before the next iteration. This strategy allows us to track more new cells in the subsequent iterations, with updated parameters and saved lineages from previous iterations. The tracking results can be visualised by drawing the connections between cells over frames as illustrated in Fig. S4.

**Fig. S4:**
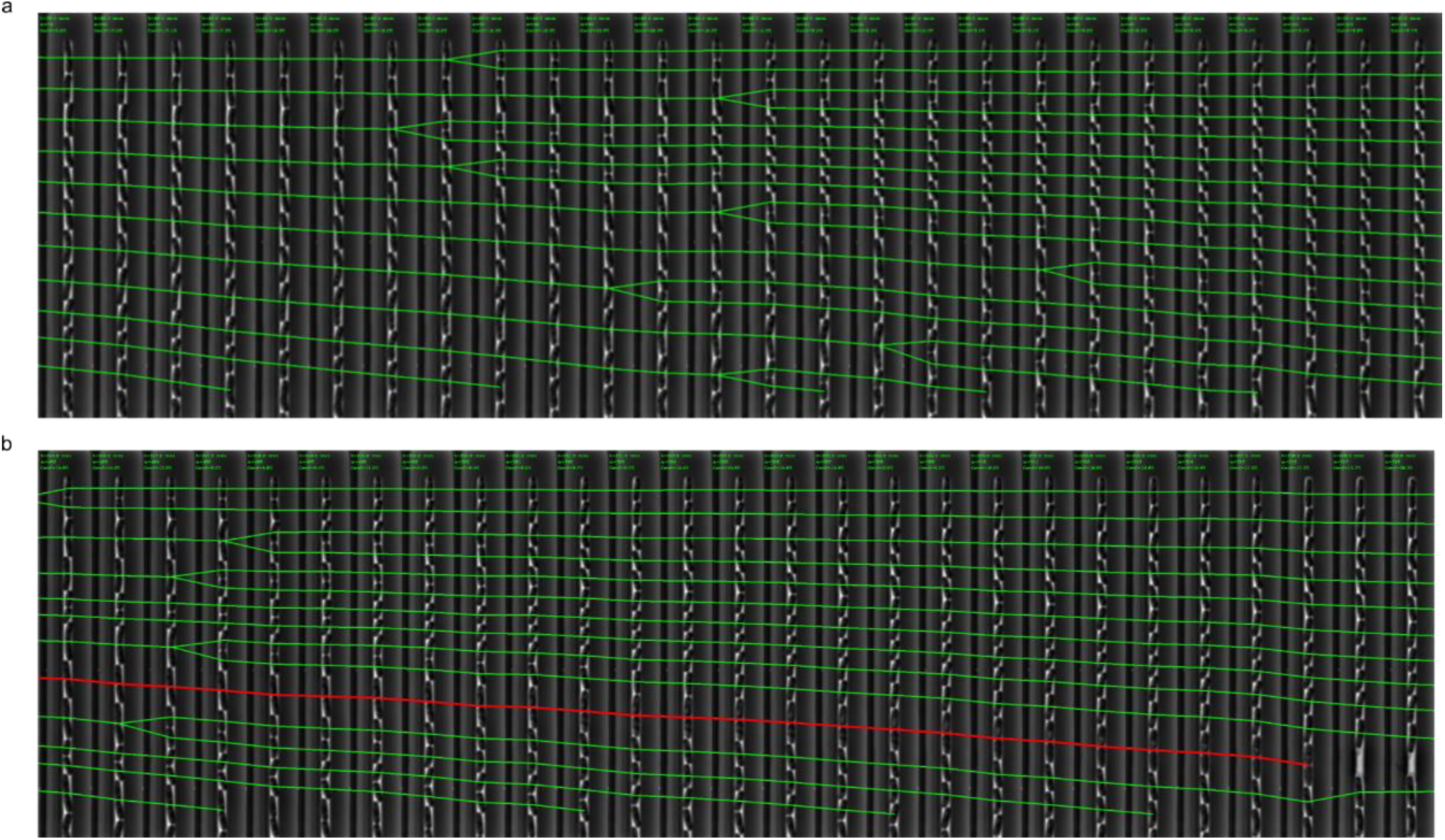
Example kymographs illustrating tracking results. **a.** A kymograph showing *E. coli* cells in a single trench undergoing balanced growth and divisions before phage infection, where the cell tracking result is shown by the green lines. **b.** A kymograph from the same trench at a different time window, showing a single cell’s growth arrest and lysis caused by phage infection, indicated by the red line, while the rest of the cells continue to grow exponentially and divide.

#### Supplementary note 6 Bandpass filtering for bleedthrough correction

When imaging T7* infection, there is a small amount of YFP bleed through into the orange channel images, due to a spectral overlap. In order to clarify Fig. 1c, we applied a band pass filter to the image in order to remove this bleed through artefact. A description of the band pass filtering process used is given in Fig. S5. Note that this bleed through correction is for visualisation purposes only, and is separate from the bleed through correction used in the genome injection analysis described in Supplementary note 8.

**Fig. S5:**
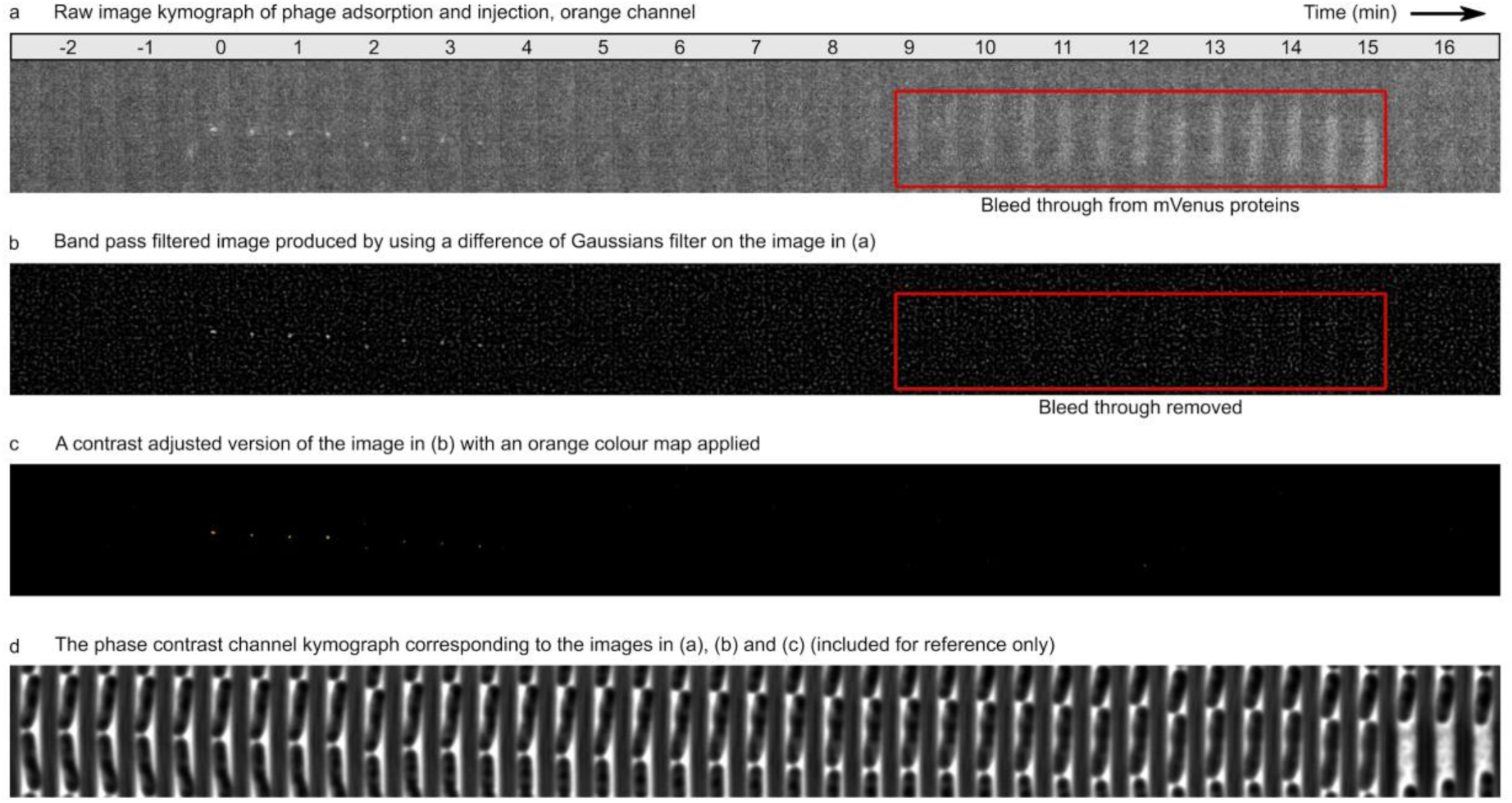
Bandpass filtering to remove YFP bleed through from Fig. 1c. The orange channel image in Fig. 1c has been filtered to remove a bleed through artefact resulting from a small amount of orange channel fluorescence produced by the mVenus expression from the phage genome (as highlighted in Fig. S5a, removed in Fig. S5b). The filtering was implemented using a difference of Gaussians band pass filter, which removes features which are not of a given size. Specifically, we used the Scikit-Image difference of Gaussians filter with low and high Gaussian standard deviations of 1 and 2 respectively^69^. The resulting image, once contrast adjusted and with an orange colour map applied (Fig. S5c), is used in Fig. 1c. The positions of the cells are shown in Fig. S5d.

#### Supplementary note 7 Example quantification of an infection event

**Fig. S6:**
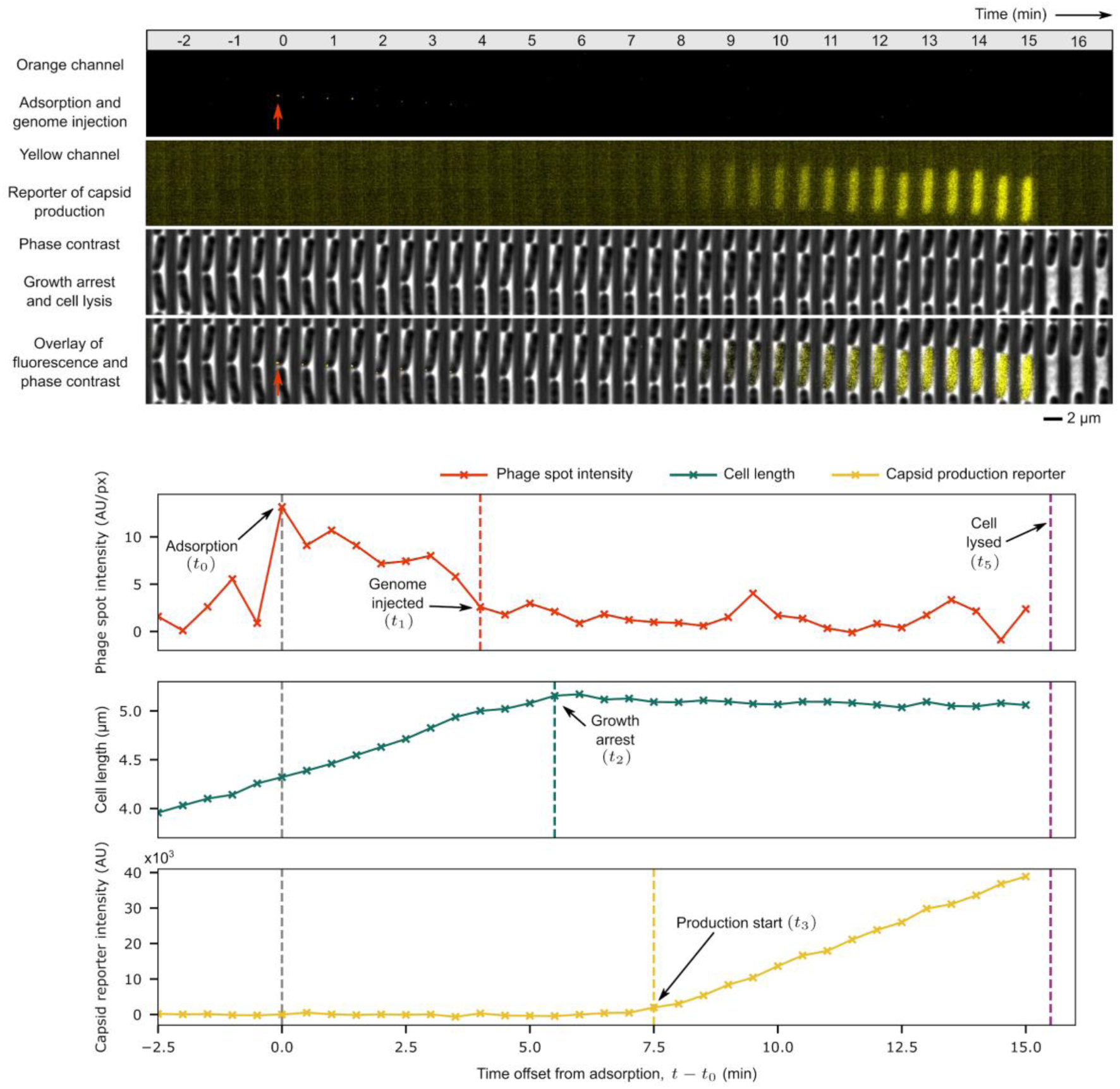
Example infection event showing quantification of genome injection, growth arrest and production start. This figure is a version of Figs. 1c and 1d, where the panels of Fig. 1d have been separated for clarity. The kymographs show the separate colour channels which are quantified to produce the time series data. The top panel of the time series data (orange line) shows the intensity of the stained bacteriophage, and how the intensity decreases as the genome is injected. The central panel shows the cell length (green line) and the subsequent growth arrest. The bottom panel shows how the capsid reporter (yellow line) expression starts shortly after the growth arrest and then increases until lysis. The vertical dashed lines indicate specific timepoints within the viral life cycle. The data points in each panel are vertically aligned with the corresponding images in the kymographs.

#### Supplementary note 8 Phage genome injection kinetics analysis

To track the injection of the phage genome, the intensity of the bright spot, indicating the location of the SYTOX Orange stained phage, needed to be tracked over consecutive frames until the genome injection was complete. For the data in Figs. 1, 2 and 6, an automated method was developed to find phage bound to cells using the Python package Trackpy^73^. Trackpy could efficiently find phage bound to cells when the phage signal was bright, but towards the end of an injection when the phage signal became dim, the track was often lost. To measure a baseline intensity prior to phage binding, and to measure the intensity in frames where Trackpy has lost the track, additional phage spot coordinates were estimated for the frames both before and after the segmented trajectory found by Trackpy. Analysis methods used for Figs. 1 and 6 compared to those for Fig. 2 differ slightly due to adjustments made to account for differences in the data and the objectives of the analysis, but the overall structure of the analysis is the same in both cases.

For Figs. 1 and 6, the spot position needed to be tracked until cell lysis, as it was necessary to measure the YFP intensity to estimate the bleed through to the orange channel for a bleed through correction. The position of the spot before and after the Trackpy trajectory was estimated by using a model of cell growth in the microfluidic device. Some of the estimated positions were then manually adjusted to ensure accuracy. For Fig. 2, we only required the intensity in the period during and adjacent to injection, so we manually found the coordinates of the 10 frames before and after the Trackpy trajectory for each injection event.

To measure the spot intensity for Figs. 1 and 6, the mean of the brightest 14 pixels in a rectangular box centred on the spot position was computed for each frame (Fig. S6a). Using 14 pixels was found to be sensitive in detecting the phage signal even when the signal to noise ratio was low. The box was made sufficiently long in the direction of the spot motion to account for any inaccuracies in the position estimation. Additionally, the mean of the brightest 14 pixels in a box alongside the spot box was computed to act as a control region. The measurements (along the short axis of the trench) of the spot and control box widths, along with the distance between the inside edges of these two boxes (the control distance), are listed in Supplementary table 6.

For Fig. 2, a similar method was used, only that the spot and control boxes measured 9×9 pixels each, and the signal intensity at each frame was measured by summing the brightest 28 pixels in the control box, and subtracting this from the sum of the brightest 28 pixels in the spot box. The smaller spot and control boxes were used as the phage locations were found more accurately, and the use of a smaller box removed some background noise. 28 pixels were used as this is the number of pixels in a circle of diameter six pixels, which was the approximate size of a phage in the image.

Due to a high SYTOX Orange background in the T7* experiment for Fig. 2, a small amount of dye often entered the living cells. Because of this, following the background subtraction using the spot and control boxes, the background subtracted baseline intensity prior to the phage binding was offset from zero. To correct for this, a constant offset was applied to the whole time series to shift the baseline intensities to approximately zero. This allowed the phage signal to be evaluated fairly regardless of any dye ingress, and for the photobleaching to be assessed accurately. The baseline offset was applied to both the T7 and T7* experiments for Fig. 2, but the difference made to the T7 experiments was negligible as the baseline was already approximately zero.

**Supplementary table 6:** Parameters used in stained phage image acquisition and in the extraction of genome injection time series data.

|  | Experiments |  |
| --- | --- | --- |
| Parameter | Figs. 1 and 6 | Fig. 2 |
| Imaging interval (s) | 30 | 30 |
| Laser power at 561 nm (%) | 100 | 100 |
| Orange channel exposure time (ms) | 50 | 100 |
| Main trench width ($\mu\text{m}$) | 1.4 | 1.4 |
| Pixel to micron ($\mu\text{m}/\text{px}$) | 0.065 | 0.104 |
| Box width (px) | 22 | 9 |
| Box length (px) | 30 | 9 |
| Gap between spot and control box (px) | 14 | 10 |
| Number of pixels used for brightest pixels (-) | 14 | 28 |
| Processing of brightest pixels | Mean | Sum |
| YFP bleed through correction | Yes | No |

In some cases, bright spots in the orange channel separate to the bound phage are observed passing through the spot and control boxes. When these are brighter than the phage and appear in the spot box, or if they appear in the control box, they will obscure the estimation of the phage spot intensity. In any frames which featured other bright spots in either the spot or control box, linear interpolation between the preceding and subsequent frames was used to estimate the true intensity.

To correct for any potential YFP bleed through, the following method was used. This was only done for the injection data in Figs. 1 and 6 which used T7* phage. First, the starting time of YFP production was found by detecting when the derivative of the background subtracted YFP signal increased significantly above zero. Then, linear regression slopes of the YFP and orange signals between the production start time and lysis were calculated. The ratio of these two slopes was calculated as a correction factor. Then, the bleed through corrected orange signal was calculated by subtracting the YFP signal divided by the correction factor from the orange signal at each time point. An example trace is shown in Fig. S6b. For the T7* injection events in Fig. 2, no bleed through correction was applied. This was because for that experiment, the SYTOX Orange background signal was high, and any bleed through from YFP expression was negligible by comparison. Moreover, the exclusion criteria for event curation for Fig. 2 excluded any events where the injection finished after YFP expression had begun (Supplementary note 11).

**Fig. S7:**
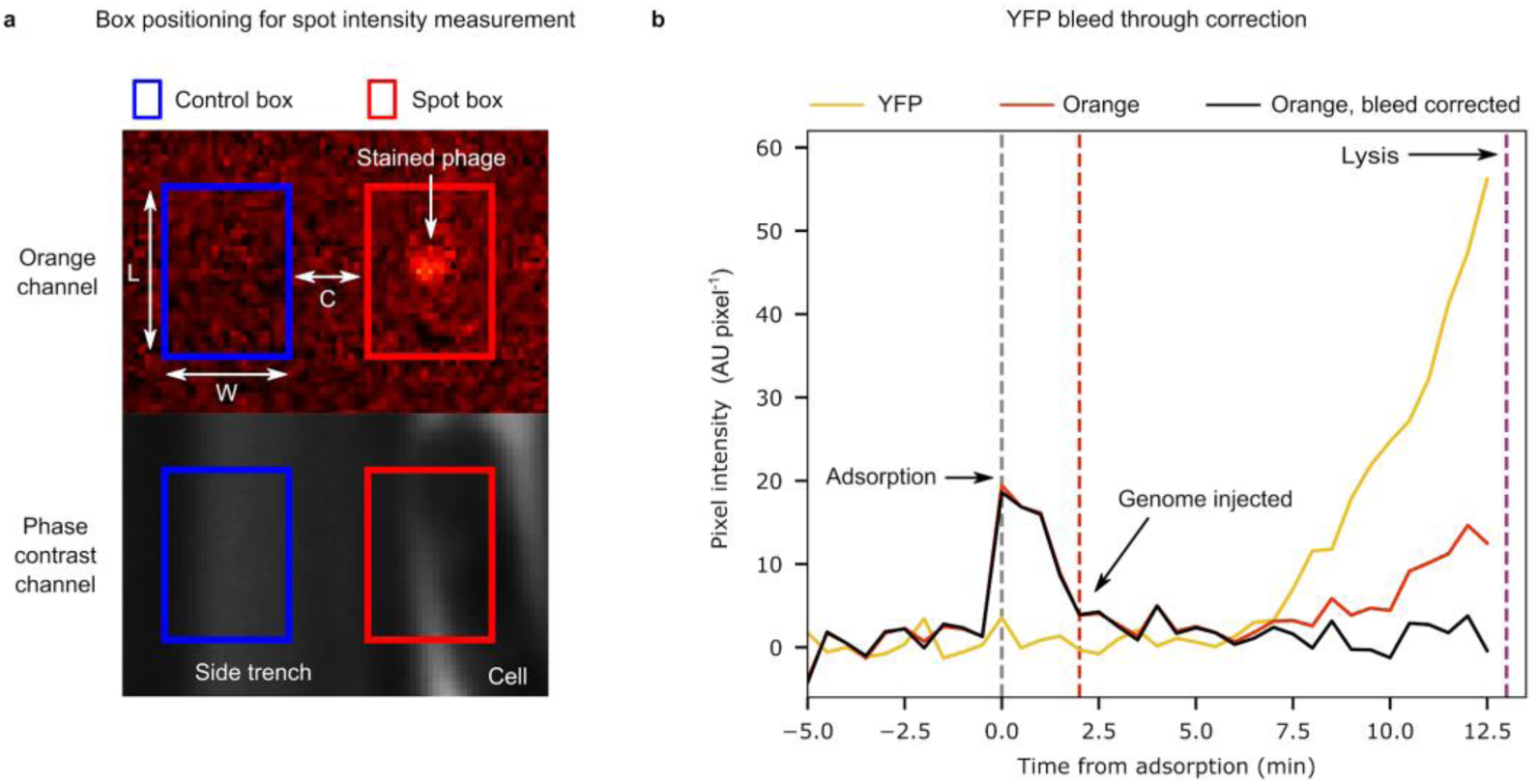
Examples illustrating the box positioning and bleed through correction used during genome injection analysis. **a.** An example of the box positioning used for the spot intensity measurement. The two images show the same scene, in both the orange channel (top) and phase contrast channel (bottom). The length (L) and width (W) of the spot and control boxes are annotated onto the drawing, along with the control distance (C) between them. **b.** An example of the bleed through correction applied to the orange signal when tracking the injection of the T7* genome for the data in Fig. 6. The YFP signal (yellow line), is the time series of the mean of the brightest 14 pixels in the yellow channel in the spot box minus the corresponding mean from the control box. The orange signal (orange line) is calculated similarly, and shows a peak as a SYTOX Orange stained phage adsorbs to the cell and subsequently injects its genome. The bleed through corrected orange signal (black line) shows how the correction method removes the second rise of orange signal (from YFP bleed through), but has a negligible effect on the early part of the data.

For Fig. 6, the duration of the genome injection was estimated by measuring the time between adsorption and the spot intensity returning to the intensity measured in the control box. The bleed through corrected signal was used, and the threshold below which the signal had to drop (for three consecutive time points) in order for injection to be considered complete was calculated as the mean plus three standard deviations of the signal in a five time point window, ending five time points prior to the starting time point as detected by the tracking algorithm. For Fig. 2, the threshold below which the signal intensity had to drop (for three consecutive time points) in order for injection to be considered complete was 20% of the initial signal intensity. This threshold was used for the study of injection kinetics as it reflects a consistent fraction of the genome between different events. The noise-based threshold used for Fig. 6 was chosen as many of the events had a low signal to noise ratio, meaning that a fixed threshold could be within the range of background noise for low signal events.

For the data in Fig. 6, the accuracy of the genome injection time was ensured by comparing the algorithmically detected injection end to an injection end time determined by inspecting the images. If the two times agreed within 2 min of each other, the algorithmic detection was considered to be accurate and used for Fig. 6. Of the 23 infection events which were selected based on the criteria in Supplementary note 11, 20 algorithmically determined injection times were in agreement with the inspection based injection time (Supplementary note 11, Supplementary table 9).

#### Supplementary note 9 Stained phage photobleaching kinetics

This supplementary note describes the control experiment and analysis to understand the photobleaching kinetics present when imaging SYTOX Orange stained phage in the absence of cells. To image the phage, we flowed SYTOX Orange stained T7 phage at a concentration of 4.4 × 10^5^ PFU/μL in LB miller into a microfluidic device (Fig. S8a, left) containing large chambers. The chambers had a width of 38 µm and lengths between >50 µm, meaning they are much larger than typical microfluidic trenches used in this study, and therefore provide a large glass surface area for phage binding. The experimental conditions, imaging settings and the objective lens were kept consistent with the experiments in Fig. 2.

The positions of any bright spots within the trenches were located using Trackpy^73^. To quantify the signal of any tracked feature (Fig 8a, right), the sum of the brightest 28 pixels in a pixel array centred on the feature was first calculated. A similar sum from a corresponding background array was then subtracted from the feature array sum. Performing this calculation at each timepoint generates a time series of the background subtracted signal. For the initial analysis, a 10×10 pixel array was used, but for the final signals used in part c onwards of Fig. S8, a 9×9 pixel array was used to reduce unnecessary background, and so that the bright spot was centred in the array. The next challenge is to identify which bright spots correspond to stained phages.

Initial analysis of the images revealed three distinct classes of fluorescent features. (1) Glass-bound features showed steadily decreasing signal intensity consistent with photobleaching; (2) Ceiling-bound features exhibited a similar decay and were identified by an Airy disk halo, likely due to defocus; (3) Additional glass-bound features displayed diverse signal profiles but retained measurable intensity throughout imaging and never appeared to bind or unbind, suggesting that they represent pre-existing SYTOX-binding impurities on the glass surface. In contrast, features in the first two classes frequently appeared and disappeared, implying transient interactions. The shorter residence times of these binding and unbinding features indicate that they originated in the medium and are therefore likely to be phage particles. In total, 32 features of the first class, 6 of the second and 49 of the third class were observed.

Because the Airy disk artefacts associated with ceiling-bound features complicated image processing, only the first class, glass-bound features showing photobleaching, was analysed (Fig. S8). An example of a glass-bound phage is presented as a “wrapped” timelapse in Fig. S8b. This example was seen to bind and then unbind during the imaging, remaining bound for 14.5 min. During that time, the signal decreases due to photobleaching. To increase our confidence that the signals used in the analysis are due to stained phage, only particles which are seen to bind or unbind (a total of 24) are taken forward. Of these, one had a signal with a sudden drop which could be explained by partial genome ejection. This particle was also excluded, leaving 23 to be used in Fig. S8c. The left panel in Fig. S8c shows the natural log-transforms of the background subtracted time series signal intensities of 23 glass-bound phage. In log space, the signals show an approximately linear decrease down to a threshold indicated by the dot-dashed line. Beyond this threshold, the signals tend to plateau and become considerably more noisy as the signal intensity approaches that of the background. As the signal kinetics in this region are difficult to discern from noise, the signals were truncated at the point where the signal first drops below the noise threshold for a minimum of three consecutive time points. All signals which were at least 10 time points long following this truncation (a total of 22 signals) were carried forward into the next part of the analysis in the right-hand panel of Fig. S8c.

The slopes in log space of each time series were then calculated, and the median slope found. A linear guide line (black dashed line) was then constructed using the median slope, with the vertical axis intercept at the maximum initial signal intensity across the 22 signals. Each signal was then shifted horizontally such that its initial value sits on the guide line. Grouping the signals in this manner (Fig. S8c, right hand panel) demonstrates that the photobleaching kinetics approximately follow a single exponential decay (linear in log space), and that the kinetics are consistent over a range of intensities.

With each signal in its new position (as plotted in the right-hand plot of Fig. S8c), the datapoints from all signals were collated and a linear regression line was calculated (blue dashed line). The residual of each datapoint relative to this central fit line was then calculated. The 2.5th and 97.5th percentile residual values were then used as lower and upper offset values respectively to construct the lower and upper bound lines (green dashed and green dotted lines respectively). A decaying signal bounded by these lines can be explained purely by photobleaching, but a signal which transgresses these bounds requires further explanation. In the case of a cell-bound phage, a signal deviating from the bounds could be explained by genome injection. Fig. S8d shows the signals of two cell-bound phage aligned to the guide line. *An injecting phage (orange line) decays rapidly, but in steps, below the lower bound and to the level of the background noise. In contrast, a different phage (purple line) binds, remains close to the guide line, and then drops in a single step to the level of the background noise*. This signal is likely explained by the phage binding, bleaching slightly while attached, and then detaching from the cell. The injecting phage shown is a wild type T7 phage used for the genome injection duration distribution in Fig. 2f, and the detaching phage was a wild type T7 phage excluded from the Fig. 2f data (Supplementary note 11, Supplementary table 8). Values of intensity in the cell-bound phage intensity time series below 50 AU were set to 50 AU before the log transform, for numerical stability.

*A further way to add confidence that the glass-bound particles used in the analysis are indeed phage, is to estimate the shape and size of the point spread function (PSF) and compare them to the PSF of cell-bound phage.* Fig. S8e shows measurements of the full width at half maximum (FWHM), a common measure of PSF size, for glass-bound and cell-bound phage. The FWHM was estimated for each observation of the glass-bound phage used to construct the time series in the left panel of Fig. S8c, with the exception of those timepoints which required interpolation due to passing particles (of a total of 1061 observations, 14 were excluded due to interpolations, leaving 1,047 observations to be plotted). The cell-bound phage (pink dots) are from the infection events used to construct Fig. 2f. A total of 104 observations of the cell-bound phage are plotted. The FWHM estimation was performed by fitting an isotropic two-dimensional Gaussian to a 7×7 pixel array at the location of the phage, centred on the brightest pixel. Curve fitting was done using Scipy^74^. The FWHM was then calculated as 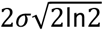, where *σ* is the standard deviation of the fitted Gaussian.

The FWHM distribution of glass-bound phage above the noise threshold (blue dots) reveals two broad groupings. The first consists of points with low amplitude and a wide range of FWHM values, largely attributable to noise when particles become very dim due to photobleaching. Under low signal-to-noise conditions, noise in the pixel array can distort the fitted Gaussian, producing unpredictable FWHM estimates. As indicated by the grey dots, most of these observations correspond to particles below the noise threshold shown in Fig. S8c. The second grouping forms a vertical band at FWHM values of 300–350 nm, which broadens slightly as amplitude decreases, again consistent with reduced signal-to-noise giving more weight to the distribution tails. The origin of the small amplitude gap between 300 nm and 400 nm FWHM values remains unclear. Importantly, the FWHM values for cell-bound phage predominantly fall into the same range of FWHM values as the glass-bound phage, forming a column between 280 nm and 330 nm. The high degree of overlap in the particle size suggests they result from the same kind of particle, implying they are both phage. *This supports the validity of this experiment as a control for photobleaching in the absence of cells*.

Fig. S8f shows an estimated genome injection profile for T7 phage, based on the results of two studies^37,38^ (see Supplementary note 10 for calculation), providing us with an estimation of the biological timescale of genome injection. If this is considerably less than the timescale of photobleaching, then it suggests that measurements of the genome injection kinetics will not be significantly affected by photobleaching.

Fig. S8g shows measurements of the stained phage intensity half-life due to photobleaching. The left-hand plot shows a comparison of the actual photobleaching half-life, determined by measuring when the intensity drops below half of its initial value for a minimum of three consecutive timepoints, and the expected photobleaching half-life. The expected half-life is calculated by fitting an exponential decay model to the data, and using the decay constant to calculate the half-life. Due to binding and unbinding, only nine actual half-lives could be measured, as most time series did not fall to half their initial value before ending. For all nine of these, half-initial intensity was below the noise threshold in Fig. S8c. There is some disagreement between the actual and expected half-lives, with six of the nine measurements showing the expected half-lives between zero and 10 min longer than the actual half-lives. The reason for the discrepancy is unclear, but it could suggest that the single exponential model of photobleaching does not capture all salient details of the photobleaching kinetics at intensities below the noise threshold. Despite this, no actual half-lives are shorter than 12 min, suggesting the timescale of photobleaching is considerably longer than the timescale of full genome injection (indicated by the red shaded area).

The distribution of expected half-lives in the right-hand plot also shows that the half-life assuming exponential decay is considerably longer than the timescale of genome injection (shaded red box). The shortest expected half-life is 8.4 min, and mean is 21.9 min.

In summary, while bleaching is significant, its timescale is considerably longer than the biological process of T7 genome injection, meaning that genome injection should account for most of the signal loss during stained phage genome injection. Moreover, in Fig. S8c and d, we provide a method to distinguish between signal loss caused solely by photobleaching and signal loss which cannot be explained by photobleaching using the results of this control experiment.

**Fig. S8:**
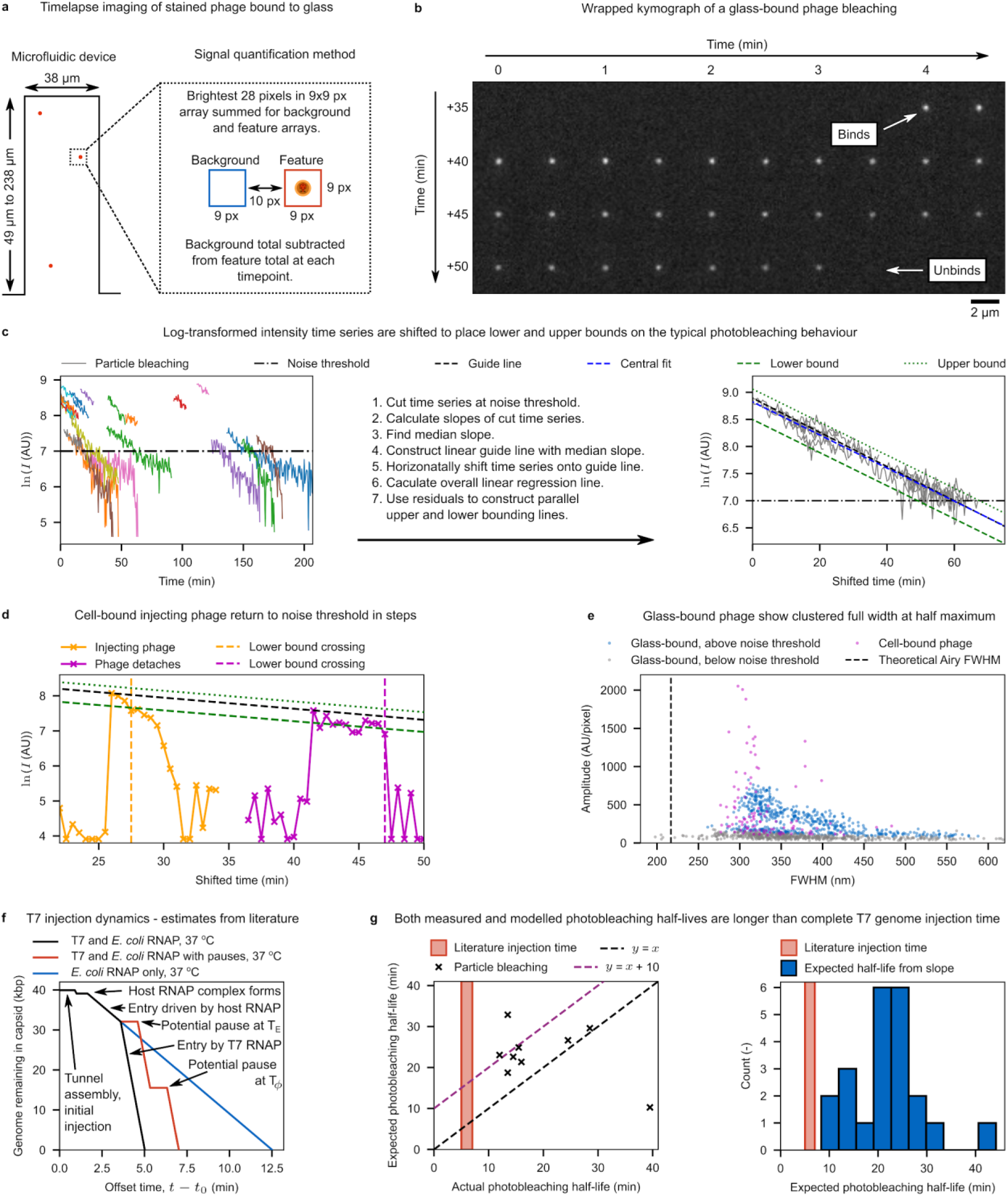
Stained phage photobleaching kinetics in the absence of cells. **a**. A diagram showing the microfluidic device used to image stained phage bound to glass. Large microfluidic chambers were used to maximise the glass surface area for phage to bind. The expanded view indicates the signal quantification method used. For each observation we sum the brightest 28 pixels in a 9×9 pixel array centred on the bright particle, and take a corresponding pixel sum from an offset 9×9 background array to perform a background subtraction. **b**. A “wrapped” kymograph of a bleaching phage bound to glass. The timepoints of binding and unbinding are indicated. The image is uniformly contrast adjusted for clarity. **c**. Natural-log transformed intensity time series of glass-bound stained phage (left panel) are subject to several operations to characterise the typical photobleaching kinetic. The decay of the log transformed signals is approximately linear, and their slopes are similar, so horizontally shifting them such that the initial timepoint falls onto the guide line (right panel) allows the typical deviation from the median slope to be assessed across particles with different intensities. The deviations from the central line are used to construct upper and lower bounding lines. Signals which start on the guide and remain within the bounds can likely be explained by photobleaching, but observations beyond the bounds can be assumed to have an additional cause. The left panel has 23 time series and the right panel has 22 time series. **d**. An example where data from phage bound to live cells are aligned with the guide line. A phage which likely injects its genome is shown by the orange line, which drops below the lower bound and then returns to the level of background noise in steps. In contrast to this is a cell-bound phage which likely detaches from the cell (pink line), as shown by a signal which adheres closely to the guide line before immediately dropping to the level of the background noise. **e**. Measurements of the full width at half maximum (FWHM) for glass-bound (blue dots, n = 540) and cell-bound (pink dots, n = 104) phage. Grey dots (n = 507) represent glass-bound phage where the total signal intensity falls below the noise threshold indicated in part (**c**). Each dot represents a FWHM estimation (obtained by fitting an isotropic two-dimensional Gaussian) for one phage at one time, collated from 23 different glass-bound phage and 12 cell-bound phage. **f**. The course of T7 genome injection as estimated from literature sources^37,38,75^ which measured the rate of T7 genome internalisation using a DNA adenine methylase based assay. The estimated time to complete injection from adsorption at 37 °C is between 5 min and 7 min. **g**. The measured and modelled photobleaching half-lives are longer than the complete T7 genome injection time. For photobleaching time series long enough to directly measure the half-life, the actual half-life is compared to the expected half-life estimated from an exponential decay model (left panel, n = 9). The black and purple dashed lines are visual guides to help estimate the difference between the actual and expected half-life. A histogram (right panel, n = 22) of all expected half-lives shows the timescale of photobleaching is typically much longer than the timescale of genome injection.

#### Supplementary note 10 Genome injection dynamics show different categories of behaviour

In this supplementary note, we compare the genome injection dynamics of single-cell single-phage injection events to the expected dynamics based on data from experiments found in the literature^37,38^. Fig. S8f (Supplementary note 9) shows an annotated genome injection profile for T7 phage based on this estimation, and they are also plotted for comparison throughout Fig. S9. These studies measured the rate of genome internalisation by measuring the timing of methylation of certain sites on the T7 genome by the enzyme DNA adenine methylase (Dam). Their experiments used LB media, with some experiments at 37 °C, using a modified *E. coli* K-12 strain and modified T7 phage variants, meaning the conditions are similar to those used in this study.

These studies measured rates of genome internalisation for each of the three phases of T7 genome entry, each likely driven by a different enzyme^33,35,37,76^. The first phase, likely driven at least in part by gp16^33,76^, was measured to proceed at 141 bp s^−1^ at 37 °C. The second phase, driven by the *E. coli* RNAP, has a measured speed of 60 bp s^−1^ at 37 °C. The final phase, driven by the T7 RNAP, has not been explicitly measured at 37 °C. *However, the internalisation speed was calculated at 30 °C*^38^ *as between 200 bp s^−^*^1^ *and 300 bp s^−^*^1^*. To estimate the speed at 37 °C, we assume that the proportional speed increase is the same as that of the E. coli RNAP over the same temperature increase, a factor of 1.446.* Therefore, we assume the speed of genome internalisation by T7 RNAP to be 361 bp s^−1^ at 37 °C.

To estimate the total time of genome internalisation, any potential pauses in the injection must be accounted for. The first is an estimated pause of 54 s at 37 °C, which accounts for the trans-envelope tunnel assembly. The second is the elongation complex formation of the host RNAP^37^, estimated as 1.1 to 2.2 min. Taking the midpoint of this range, and accounting for the time of tunnel formation and the gp16 mediated delivery, we estimate this step to take 39 s. There are then two further potential pauses, one at the early terminator for host RNAP, T_E_, and another at the T7 RNAP terminator, T_φ_. While the authors do not provide estimates for these pauses, they do note that a pause of less than 60 s at T_E_ would not be detected by their assay^37^. Therefore, we conservatively include a scenario with a pause of 60 s at each of the terminators.

Taking into account all these estimates, and using the genome length and terminator positions published by Dunn and Studier^75^, we provide estimates of the T7 genome delivery times under different scenarios in Supplementary table 7, and provide a corresponding estimated injection profile in Fig. S8f above.

**Supplementary table 7:** Literature based estimates of T7 genome injection time.

| Step | Injection time -<br>No terminator<br>pauses (s) | Injection time -<br>Terminator<br>pauses (s) | Injection time<br>- Host RNAP<br>only (s) |
| --- | --- | --- | --- |
| Tunnel assembly (0 bp) | 54 | 54 | 54 |
| Transcription independent delivery (850 bp) | 6 | 6 | 6 |
| Elongation complex formation (0 bp) | 39 | 39 | 39 |
| Host RNAP delivery (6738 bp) | 112 | 112 | 112 |
| Pause at $T_E$ (0 bp) | 0 | 60 | 0 |
| T7 RNAP delivery – first stage (16621 bp) | 46 | 46 | 277 (delivery by host RNAP) |
| Pause at $T_\phi$ (0 bp) | 0 | 60 | 0 |
| T7 RNAP delivery – second stage (15727 bp) | 44 | 44 | 262 (delivery by host RNAP) |
| **Total** | 301 | 421 | 750 |

We now look to compare our own results to these estimates. To account for the signal decay caused by photobleaching, we apply a simple correction to estimate the underlying fraction of the genome remaining in the capsid. If we assume that the SYTOX Orange dye is evenly distributed along the length of the genome, then the observed signal at a given time (*S*(*t*)) is the product of the fraction of the genome remaining in the capsid (*G*(*t*)), and the expected signal magnitude following photobleaching if the entire genome remained in the capsid (*B*(*t*)). If *G*(*t*) and *B*(*t*) are normalised between one and zero, then *S*(*t*) is given below, where *S*_0_ is the initial signal magnitude:

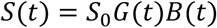

The function *B*(*t*) approximates an exponential decay, as described in Supplementary note 9, with a decay constant of 0.032 min^−1^. Therefore, the fraction of the genome remaining in the capsid at any given time can be approximated by *G*(*t*), where *t* starts at zero at the time of adsorption:

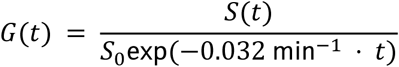

The length of the genome remaining in the capsid is then found by multiplying *G*(*t*) by the length of the genome of the given phage. The observed signals used to calculate the genome injection durations in Fig. 2f have been converted into genome lengths and divided into *five categories* in Fig. S9. One T7* injection has been excluded from Fig. S9, as there were transient fluorescent artefacts distorting the injection dynamics. Each category has the expected injection dynamics calculated from literature sources (as in Fig. 2g, calculated in Supplementary table 7) plotted for comparison. While the parallels to existing theories are interesting, further experiments to test specific and alternative hypotheses would be necessary, along with larger sample sizes, to support or reject any of the potential explanations.

*The first category* (Fig. S9a) shows a slow rate of genome entry for the first 1.5 min to 2 min, before transitioning to a more rapid genome entry. The transition we observe to a rapid genome entry happens sooner than would be expected from the literature estimate. However, if it is assumed that the *E. coli* RNAP begins translocating the genome almost immediately, then our observations could be well-explained by the speed of the enzymes. The initial speed is comparable to that observed in the second category (Fig. S9b), where entry proceeds at approximately 3.6 kbp min^−1^ to 4.0 kbp min^−1^. At this rate of entry, approximately 6 kbp of the viral genome would enter the cell in the first 1.5 min. We observe the transition to a faster entry speed when approximately 6 kbp to 12 kbp of the genome has entered the cell. Gene 1, encoding the T7 RNAP, ends at genomic location 5,820 bp and the second T7 promoter, *ϕ*_1.1A_, is located at position 5,848 bp^75^. The first category therefore has good consistency with the hypothesis that translocation of the first part of the T7 genome is carried out by the *E. coli* RNAP, and the rest by the much faster T7 RNAP.

While the above is a broadly accurate description of the entry kinetics for the first category, there is some noise in the measurements which could potentially obscure other biological phenomena within the injection process. For instance, while the initial entry rates for some category 1 and 2 injections are close to 3.6 kbp min^−1^, others are closer to 8.5 kbp min^−1^, which is the expected speed of the gp16 driven transcription-independent genome injection at 37 °C. At this stage, we cannot say whether this is due to measurement noise or whether transcription-independent delivery plays a larger role at the beginning of genome injection for some events. Conversely, some noise in the measurements may be caused by the biological process. The entry rates during the fast entry stage of category 1 injections fluctuate considerably, which may be caused by intermittent transcriptional pausing by the T7 RNAP enzyme.

The discrepancy between our observations and the literature-based estimate in Supplementary table 7 for the timing of the switch to faster entry could potentially be explained by a limitation of the Dam methylase assay, the method used in the studies forming our literature estimates. This limitation is caused by the way in which T7 controls the access of host proteins to its entering genome. For instance, one study^77^ has shown that the restriction enzymes EcoB, EcoK and EcoP1 cannot access the entering T7 genome until 6 min to 7 min after infection starts at 30 °C. This transcription-independent^78^ inaccessibility of host proteins to the early genome includes Dam, which is why the methylation of sites in the early regions of the genome does not occur until after the early gene products have been translated^38^. This creates a major limitation for the Dam methylation assay in determining the kinetics of genome entry for the early genome. However, this limitation does not seem to exist for our genome staining assay. Therefore, our results may reflect a more accurate representation of how the early genome exits the capsid.

*The second category* (Fig. S9b) shows genome entry at an approximately constant rate throughout the injection period, and the rate is comparable to that which would be expected by *E. coli* RNAP driven internalisation at 37 °C (3.6 kbp min^−1^). Events of the third and fourth categories have a high initial rate of genome entry, which differentiates them from the first and second categories. *The third category* (Fig. S9c) shows approximately linear kinetics, but at a rate approaching the expected T7 RNAP driven internalisation speed at 37 °C (measured rates are approximately 15 kbp min^−1^ to 18 kbp min^−1^ for wild type T7, compared to the estimated expectation of 22 kbp min^−1^). However, unlike the second category, some events show short pauses with between 15 kbp and 25 kbp remaining in the capsid. A potential explanation for the initially high rate of genome entry could be that the T7 RNAP enzyme is present from a previous unstained infection, whether live or defeated by host defence mechanisms, and uses the *ϕ*_OL_ promoter at genome location 405 bp to rapidly pull the genome into the cell^38,75^. *The fourth category* (Fig. S9d) shows events which generally start with a high rate of genome entry, which then decays as the capsid empties. The reasons for the dynamics of the fourth category are unclear, but they may also start with the T7 RNAP present from a previous unrelated infection, which is then slowed by an unknown mechanism. *The fifth category* encompasses events which show little to no entry initially, then a sudden drop with an instantaneous delivery speed of approximately 60 kbp min^−1^, far in excess of the speed of the T7 RNAP enzyme. These events could represent some malfunction of the T7 ejectosome which causes the genome to be delivered in an uncontrolled manner.

**Fig. S9:**
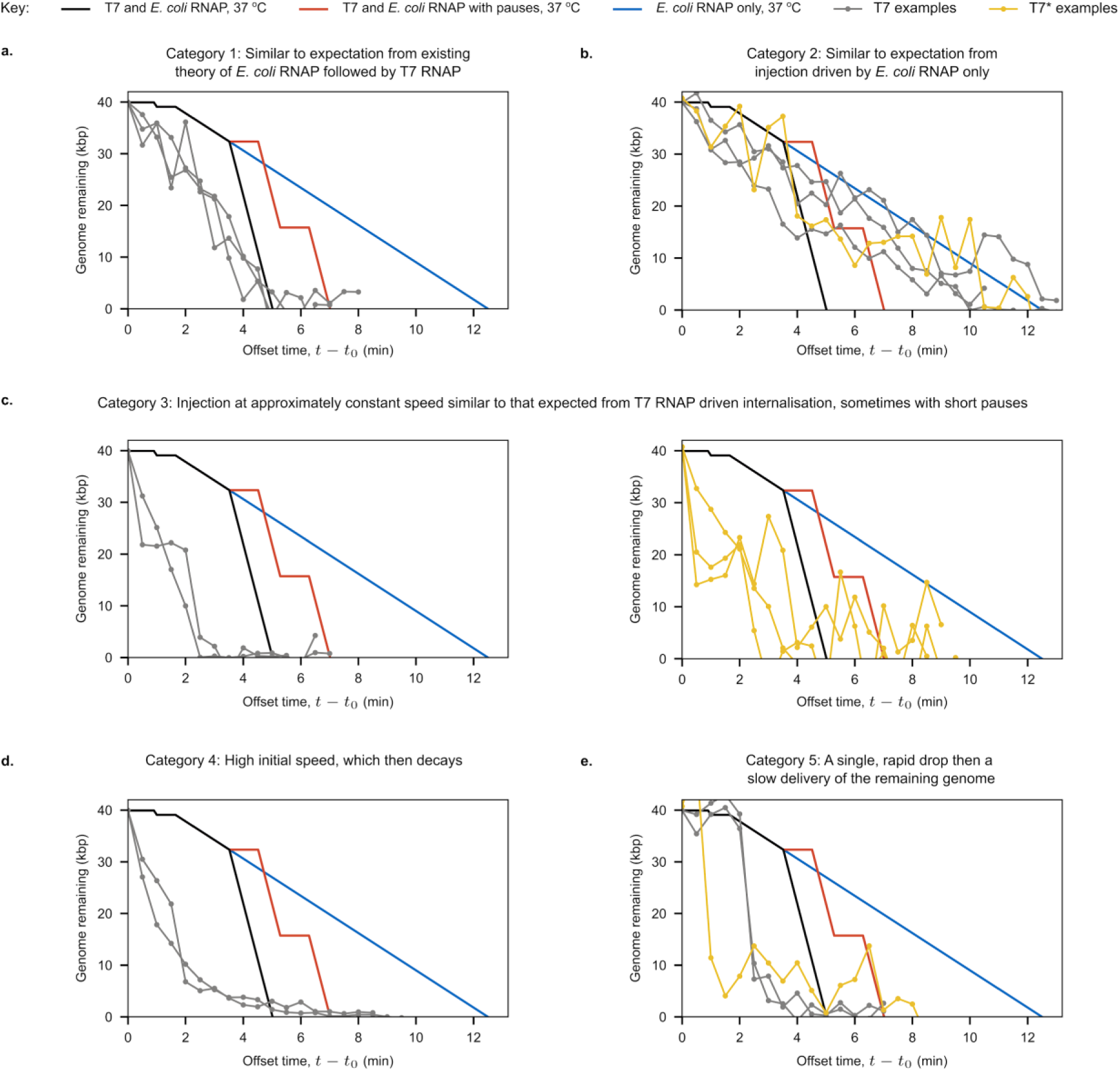
Categories of genome injection dynamics. The genome injection dynamics from the underlying events from Fig. 2f can be approximately grouped into five categories. For each category, an estimation of the genome injection dynamics based on literature sources is presented for comparison. **a**. The first category shows an initially slow injection and transitions to a faster rate after approximately 2 min. **b**. The second category shows injection at a constant, slow rate. **c.** The third category shows injection at a faster constant rate, sometimes with pauses when between 15 kbp and 25 kbp of the genome remain in the capsid. **d.** The fourth category generally shows a fast rate of initial injection, which then decays in rate. **e.** The fifth category shows little to no injection initially, followed by a sudden drop of approximately 30 kbp over 30 s.

#### Supplementary note 11 Data curation for stained phage genome injection analysis

In attempting to track genome injection events in the mother machine, it is not always the case that a single, bright, SYTOX stained phage binds to a host cell and injects its DNA, causing a subsequent infection and lysis of the host. In fact, such an initiating event paired with an observed lysis outcome, without obvious confounding factors, is relatively rare. When we observe a phage binding to a cell, it will typically either inject its DNA, or detach from the cell without any significant loss of fluorescence signal. Additionally, different phage particles can bind to the same cell over time. There are also differences in outcome. If a phage appears to inject DNA into a cell, it may lead to a subsequent lysis, but it may also leave the trench before the infection cycle is completed. Often, no apparent infection follows from an injection event, possibly from the presence of anti-phage defense mechanisms within the host cell. Even if an injection and a corresponding lysis are observed, other confounding events such as the lysis and release of phage from a separate cell can occur during the infection cycle. Therefore, to select clean examples of infection events (in order to exclude as much interference from factors separate to the one-to-one phage to host interaction we wish to study as possible), we curated data for analysis in accordance with strict criteria.

Two sets of experiments were conducted in this study where the genome injection has been analysed. These are discussed in Fig. 2 and Fig. 6, and the primary scientific objectives of these two sets of experiments were different. For the experiments in Fig. 2, the primary aim was to study genome injection dynamics. For Fig. 6, the primary aim was to study the adsorption to lysis time distribution. Due to the different objectives, different experimental parameters were used, which have subsequently led to the nature of the data being different, and the need for different selection criteria.

Two experiments (one with T7, one with T7*), both using a phage titre of 1.0 × 10^6^ PFU μL^−1^, were analysed for Fig. 2. A higher titre was used in these experiments in order to increase the number of stained phage-host interactions (Supplementary note 20, Supplementary note 21). To increase our confidence that any given observed genome injection led to a productive infection, we restricted our search for events to the first cell to lyse in any given trench, checking each of these lysing cells for interactions with stained phage. Due to the higher phage titre used, phage-host interactions were numerous in these datasets, meaning that many of the lysing cells had more than one phage interact with the cell prior to lysis. The selection criteria for Fig. 2 therefore aimed to isolate one-to-one phage-bacteria interactions which lead to a productive infection. To systematically sort the data, we defined a single-cell metric we call the multiplicity of adsorption (MOA), a measure which describes how many stained phage adsorb to a cell in a given time period. This is related to the multiplicity of infection (MOI), but it accounts for the observation that not all adsorbing phage lead to successful infections. The MOA does not make any assumptions about the nature of the interaction between the phage and the bacteria, only that an interaction occurs. For the first lysing cell in each trench, we manually screened the images and noted the value of the following metrics:

- The MOA in the time period starting 30 min before lysis, and ending with lysis. We were searching for genome injections, so we only considered adsorptions which lasted a minimum of three consecutive frames (images were taken at 30 s intervals), unless a two-frame adsorption appeared to strongly show genome injection (this was very rare). If an adsorption started before the 30 min window but crossed over into the window, this was included. The MOA for a given cell was equal to the number of phage adsorption events meeting these criteria.
- The MOA for the injection period (MOAI, MOA for Injection), is calculated using the same criteria as the MOA, but only considering the time period where the injection causing the infection could reasonably have occurred. For T7*, this was defined as the period starting 30 min before lysis, and ending when the YFP signal from capsid production became visible. For T7, no yellow signal is available, so it is more difficult to judge when a cell is infected. Therefore, the period for MOAI calculation for T7 was defined as starting 30 min before lysis and ending 10 min before lysis.

After the first cell in a trench has lysed, it becomes very likely that other cells in the trench become infected with unstained phage at an MOI greater than one, which is why the analysis is restricted to the first lysing cell.

For all events with an MOAI equal to one, the phage signal was quantified as described in Supplementary note 8. The signal is compared to the photobleaching decay (described in Supplementary note 9), and any signal deviating beyond the bounds of expected photobleaching is assumed to be driven by genome injection. If the signal and its comparison to photobleaching meet any of the following criteria, the event is excluded:

1. If a signal decays in line with the expected rate of photobleaching (Supplementary note 9), and then immediately falls into the range of the background noise, we assume it has detached without injecting and exclude the event.
2. If the decay of the phage signal appears to be completely driven by photobleaching but does not detach.
3. If, once the timepoint at the end of injection has been quantitatively determined, we find that the injection ends beyond the MOAI time period (for example, after YFP expression has started).
4. If the initial signal magnitude is less than the noise threshold used in the photobleaching analysis (1097 AU), the signal is likely too dim to accurately quantify the injection kinetics.
5. If the bound phage is obscured by a fluorescent artefact for several frames such that the injection dynamics are unclear.
6. If an injection stalls for an excessively long time (more than 10 min), we assume this is not representative and exclude the event.

If the event does not meet any of the above criteria, it is included in the statistics for Fig. 2f. Supplementary table 8 shows how the above criteria are applied to the wild type T7 experiment used for Fig. 2 to find cells with isolated genome injection events which can be used for analysis.

**Supplementary table 8:** Data curation for the T7 experiment used in Fig. 2.

| **Totals** | **Exclusion category** | **Number of exclusions** | **Subtotal following exclusions** |
| --- | --- | --- | --- |
| **Total trenches** |  |  | **120** |
|  | Trench has cells filling the side trenches | 2 | 118 |
|  | Trench is empty | 6 | 112 |
| **Trenches with cells** |  |  | **112** |
|  | No lysis in trench | 4 | 108 |
|  | It is unclear which cell lyses, or information prior to imaging start or beyond the crop near trench exit prevents categorisation | 4 | 104 |
| **Total lysis events where MOAI can be evaluated** |  |  | **104** |
|  | MOAI = 0 | 70 | 34 |
|  | MOAI >= 2 | 17 | 17 |
| **Total events with MOAI = 1** |  |  | **17** |
|  | Phage detaches | 1 | 16 |
|  | Phage has no injection, only photobleaching | 0 | 16 |
|  | Phage-host interaction ends beyond the MOAI period | 3 | 13 |
|  | Phage signal is dim | 1 | 12 |
|  | Phage is obscured by fluorescence artefact for multiple frames | 0 | 12 |
|  | Genome injection has an excessive stall of >10 min | 0 | 12 |
| **Total events to be included in Fig. 2f** |  |  | **12** |

Following this screening process in Supplementary table 8, 12 events were isolated for T7 and used in the genome injection duration distribution in Fig. 2f. Following the same set of criteria for the T7* experiment, from 135 trenches 12 cells had an MOAI of one, of which two detached from the cell, one was obscured by a fluorescent artefact, one showed only photobleaching and two finished their injection after the end of the MOAI period, leading to a total of six events being used for Fig. 2f.

Supplementary table 8 shows that 70 of 120 events in the T7 experiment had an MOAI of zero, implying that at least 58% of infections were caused by unstained phage, which could have been released by the lysis of cells in other trenches or were present in the supplied media due to incomplete staining. It is also possible that some of the infections apparently caused by stained phage may have been caused by an underlying unstained phage infection. However, a current limitation of our method is that we cannot yet estimate the fraction of observed infections with a stained genome injection which were caused by unstained phage. Calculating this estimation from the relevant conditional probabilities would require careful accounting of all stained phage-host interactions and the resulting cell fates, and all observed infections, and whether they were preceded by a stained phage-host interaction in an experiment. Such a task is too labour intensive to complete manually, and we do not yet have an automated pipeline to accurately record the details of all infections and interactions as described.

Supplementary table 8 demonstrates the rarity of one-to-one stained phage-host infections in this experiment (which uses a high phage titre) and the necessity of careful screening. In the same experiment, of 120 trenches, only 20 had an MOA of one, and only five had an MOA and MOAI of one. Of these five, only two passed through the final set of criteria and were included in the statistics for Fig. 2f. This demonstrates the need to use lower phage titres to observe enough isolated events to construct an adsorption to lysis time distribution.

The set of four experiments for Fig. 6 (three with T7*, one with T7) used lower phage titres (Supplementary note 20), meaning infections were less frequent, but more likely to be isolated when they did occur. For Fig. 6, infections were selected using the following criteria:

1. We exclude all events where an injection is suspected, but the SYTOX stained phage spot, if any, is too dim to be confidently differentiated from background noise. These events are excluded to give confidence that all injections recorded are genuine, and to ensure there is sufficient signal to analyse the injection kinetics.
2. We exclude all events where an injection is suspected, but the SYTOX stained phage spot shows no sign of dimming over two or more frames, before the spot completely disappears in one frame without sign of injection. It is difficult to confirm that such events are not simply the phage leaving the cell, so they are excluded.
3. In any cases where a cell lyses following a successful injection, but either the lysing cell or its mother also had a second phage bind and appear to inject, it is impossible to know which phage, or if both, led to the subsequent infection and lysis. All such events are excluded.
4. We exclude any events where an infected cell is about to lyse, but before it does a second cell lyses. In these cases, it is difficult to know if the release of new phage or phage lysis proteins influences the course of the infection (such as by hastening the onset of lysis), and therefore we exclude these events.
5. We exclude any events with an excessively long lysis time of >1 hour. These events are unlikely to be representative of a typical infection, and could indicate a defective phage or a partially resistant bacteria. Furthermore, when lysis times are excessively long, it becomes increasingly difficult to be confident that the observed injection was responsible for the subsequent lysis, as an unstained phage has more time to interact with the host cell.
6. Any other uncommon or unusual features of the injection and the subsequent infection profile may lead to its exclusion from the data set. For example, an injection profile which fluctuates in intensity would likely be excluded.

Data for Fig. 6 were curated according to these criteria. All single-cell lysis times for each experiment (performed on different days), curated following the above criteria, are shown in Fig. S10. We observed an unusual distribution of lysis times in the data for wild type T7, where three of the events happened to have lysis time close to twice the mean of the rest of the population. By excluding all events with a lysis time of 30 min or above from our single-cell measurements, the estimate for single-cell mean lysis time became similar to the population level measurements of lysis time. We believe this to be a justifiable way to compare the two methods, as in a one step lysis curve (Supplementary note 18, Fig. S14), infections with a lysis time of 30 min or more would not influence the first “step” used to estimate lysis time.

**Fig. S10:**
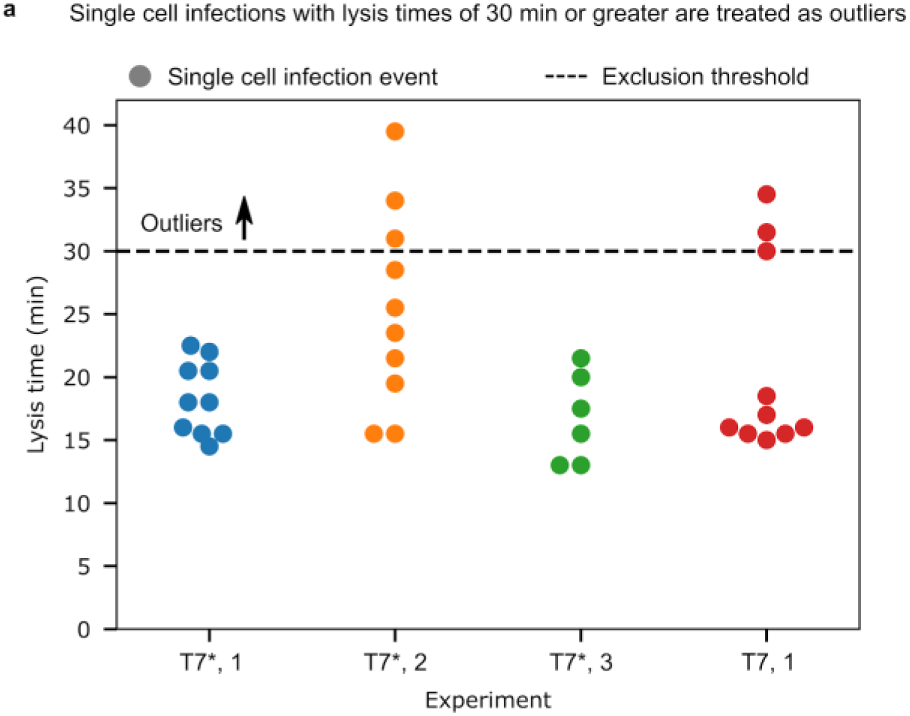
Single-cell lysis times with outliers indicated. **a.** Single-cell lysis times for the three T7* experiments and the one T7 experiment used for Fig. 6. The data for wild type T7 (red circles) shows an unusual bimodal distribution. The dashed black line at 30 min is used to show that all events with a lysis time of 30 min or greater are treated as outliers and excluded from the data used for Fig. 6. Differences between the means of T7* 1 to 3 are significant when outliers are included, but are not significant when the outliers are excluded (one-way ANOVA).

The final two steps of the data curation are summarised in Supplementary table 9. The first is using only those events with a lysis time of less than 30 min, as discussed in Fig. S10, and is used for the parts of Fig. 6 presenting the adsorption to lysis time. The second is the requirement for the algorithmic detection of genome injection to match the estimation from inspecting the images, as described in Supplementary note 8. The event numbers described in this row of Supplementary table 9 are used for the parts of Fig. 6 presenting genome injection time.

**Supplementary table 9:**
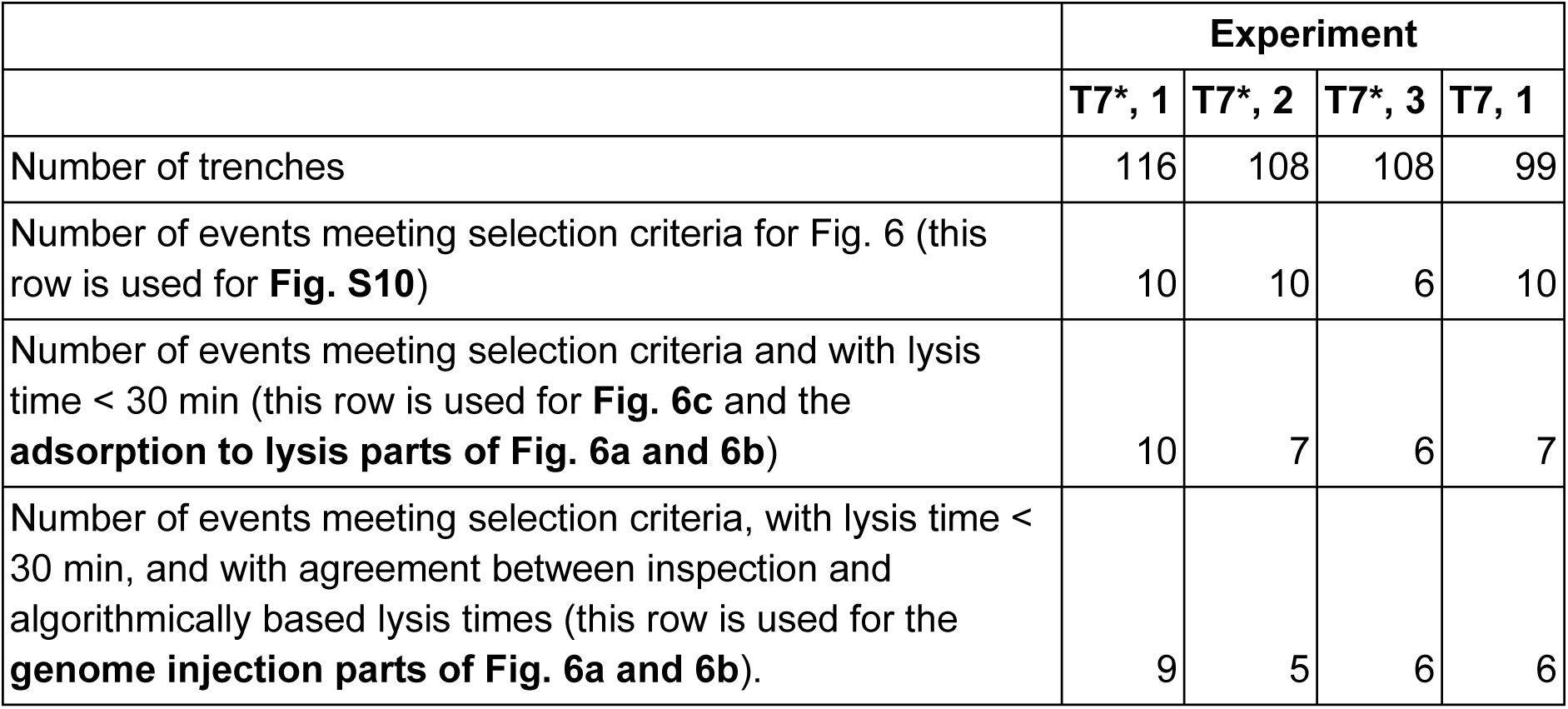
Data curation for Fig. 6.

|  | Experiment |  |  |  |
| --- | --- | --- | --- | --- |
|  | T7*, 1 | T7*, 2 | T7*, 3 | T7, 1 |
| Number of trenches | 116 | 108 | 108 | 99 |
| Number of events meeting selection criteria for Fig. 6 (this row is used for **Fig. S10**) | 10 | 10 | 6 | 10 |
| Number of events meeting selection criteria and with lysis time < 30 min (this row is used for **Fig. 6c** and the **adsorption to lysis parts of Fig. 6a and 6b**) | 10 | 7 | 6 | 7 |
| Number of events meeting selection criteria, with lysis time < 30 min, and with agreement between inspection and algorithmically based lysis times (this row is used for the **genome injection parts of Fig. 6a and 6b**). | 9 | 5 | 6 | 6 |

#### Supplementary note 12 Estimation of growth rate and determination of point of growth arrest

To calculate the cell growth rate, *λ*(*t*), for a given cell, we first calculated the natural logarithm of the cell length in μm. Then, we used a Savitzky-Golay filter^79^ with a third-order polynomial and a window size of eight data points to smooth the signal. The growth rate was calculated as the first derivative of this smoothed signal (the difference between sequential data points divided by the imaging interval).

To calculate the time of growth arrest, an average, uninfected growth rate was calculated from 75 uninfected SB8 cells (0.0304 min^−1^). A threshold growth rate for growth arrest was then set as half this value. We declare growth arrest (*t*_2_) as the first point where the growth rate drops below this threshold, if the growth rate remains below this threshold for a minimum of four consecutive time points.

The growth arrested cell length, *L*_*GA*_, is calculated as the mean cell length between growth arrest, *t*_2_, and the penultimate length measurement before lysis. The final length measurement prior to lysis is not included in the calculation, as the final measurement is susceptible to sudden length changes which may occur due to segmentation error if the image is captured just prior to lysis. Any such changes would not be representative of the cell length throughout the infection period, which otherwise remains approximately constant.

#### Supplementary note 13 YFP signal loss before lysis is caused by phage induced perforation of the cell envelope

The concluding stage of the infection process involves the lysis of the host cell. As evidenced by the data presented in Fig. 3d and Fig. 5, cells undergo a phase of perforation, characterised by a partial loss of YFP fluorescence signal and phase contrast, before complete lysis ensues.

When imaging T7* infection with a 30 s imaging interval (Fig. 3), we observed a fraction of cells (23.3%) to show a drop in YFP signal in the final observation before lysis. We hypothesised that this drop was due to YFP molecules leaving the cell due to the perforation of the cell envelope. If this were to be the case, we would expect the YFP signal to drop as the phase contrast signal increases (indicating a loss of material from the cell). Example time series data from two infected cells, one showing a drop in YFP at the final observation, and one which does not exhibit a drop, are compared in Fig. S11a to illustrate this difference. The perforation-induced anticorrelation between the YFP intensity and phase contrast pixel intensity is more pronounced in the rate of change of these variables. The rate of change of YFP intensity is plotted as a function of the rate of change of phase contrast intensity in Fig. S11b. This demonstrates how in the penultimate observation before lysis, the two derivatives are uncorrelated (left, top panel), but in the final observation before lysis (left, bottom panel), a fraction of cells show a negative YFP intensity derivative and a large positive phase contrast intensity derivative. This generates a strong negative correlation, implying that cells which lose more material (greater phase contrast intensity derivative) also show greater losses of YFP intensity. By checking the value of the correlation coefficient over time (Fig. S11b, right panel), we see that these two variables are uncorrelated until perforation begins.

**Fig. S11:**
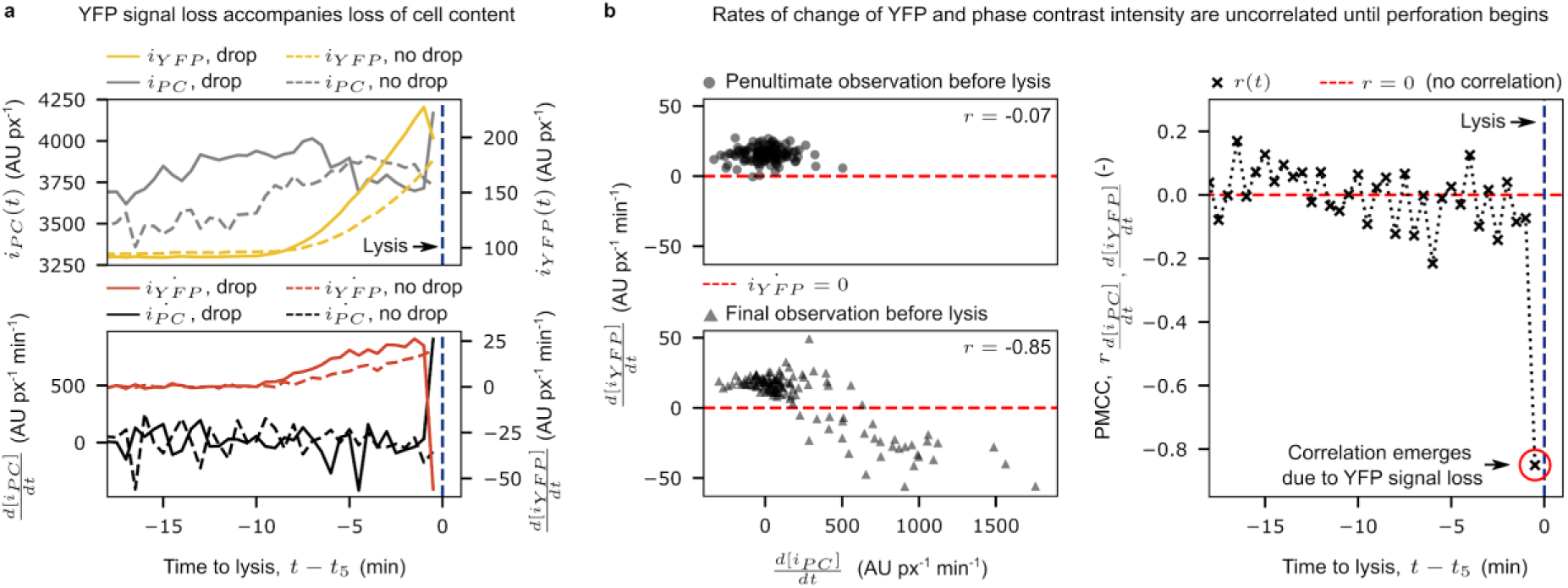
YFP signal loss prior to lysis is caused by the perforation of the cell envelope. **a.** When imaging infection at 30 s intervals, a fraction of cells (23.3%, n = 133) are shown to exhibit a loss of YFP in the final observation before lysis. This loss of YFP is accompanied by a corresponding increase in phase contrast intensity. Example time series data of YFP and phase contrast intensity (top panel) and the corresponding derivatives (bottom panel) are shown for two cells, one in which a drop in YFP is observed, and one where no drop is observed. **b.** The correlation between the rate of change of phase contrast intensity and the rate of change of YFP intensity at the penultimate observation before lysis (left, top panel) and the final observation before lysis (left, bottom panel). In the final observation before lysis, a fraction of cells show a large, positive rate of change of phase contrast intensity and a concomitant negative rate of change of YFP intensity, indicating the cells are becoming brighter in phase contrast while losing YFP signal. Analysis of the product moment correlation coefficient (PMCC, *r*) over time (right panel) demonstrates that such a correlation only exists for the final observation before lysis.

As a further check that perforation is causing the YFP drop seen in a fraction of cells, we can compare the fraction of infections in which a YFP drop is seen (23.3%) to the fraction of the final imaging interval which we expect to be occupied by perforation. The mean time between perforation and the onset of lysis is 6.56 s, and the imaging interval is 30 s, meaning that we would expect approximately 21.9% of the final observations before lysis to be of cells undergoing perforation. This closely matches the fraction of cells observed to have a drop in YFP in the final observation before lysis.

#### Supplementary note 14 Data curation for growth arrest and capsid production analysis

In Figs. 3 and 4, we aimed to use isolated infections resulting from one-to-one bacteria-phage interactions. In curating the data, we used only the first lysis in a given trench, as after the first lysis the probability of infections with an MOI greater than one is very high. Despite this, in high-throughput bacterial imaging studies such as this there can be many technical and biological factors which can affect the data quality and the relevance of the data to the scientific objective. Furthermore, there are sometimes technical reasons why the calculation of a specific quantity is not possible. To ensure that the data was as free from interfering factors as possible, we curated the data in accordance with specific criteria. These are listed under the column “Exclusion category” in Supplementary table 10. Where these are not self-explanatory, they are explained below.

- Simultaneous infections: Low phage titres were used for these experiments (Supplementary note 20), giving a low probability per unit time that a cell in a given trench would become infected. In some trenches, other cells in the same trench would be infected at the same time as the first lysing cell. While this can happen by chance, it can be indicative of a transiently high phage concentration in the trench, potentially caused by the transfer of phage progeny released by lysis from an upstream trench. As this could reflect an increased chance of the first lysing cell being infected with an MOI greater than one, we exclude these events.
- Criteria relating to the cell being in a “halo”: In the phase contrast images, there is a strip near the trench exit (in the direction of the short axis of the trench) with high pixel intensity. We refer to this strip as the halo, and it is caused by the diffraction of the illumination light off the edge of the feeding lane structure. We initially attempted to segment the whole phase contrast image, but the segmentation was unreliable in the halo. For this reason, we segmented only a cropped version of the trench with the halo removed to achieve accurate segmentation. For this reason, we were unable to obtain accurate quantitative information from cells in the halo, but could accurately determine the timepoint of lysis by inspecting the images.
- Twin daughter cells are infected and lyse: A subset of infected cells would finish division during infection, leading to two infected daughter cells. However, this creates ambiguity regarding which cell, or both, should be quantified and included in the statistics. To avoid this ambiguity, all such events were excluded.

For Figs. 3b, 3c and 3d, 23 example events are used to illustrate the nature of the data and to highlight some key features. These events are from the experiment “T7*, 1” and result from an earlier version of the data selection criteria and processing code. Despite this, they are highly representative of the data used for statistical analysis in Figs. 3e, 4 and S11 (Supplementary note 13). Of the 23 cells used in Figs. 3c and 3d, 20 were considered for inclusion in the updated data set described in Supplementary table 10. Of these, four were filtered by the introduction of the requirement to to exclude twin cells. Of the remaining 16, 16 were used for growth arrest and capsid production start measurements, and 14 were used for the full expression profile, meaning there is a high degree of overlap between these examples and the data used to generate the statistics. The fraction of cells showing a drop in YFP prior to lysis is also representative of the final statistics (Supplementary note 13). The main change in the data processing for the capsid production data between the example plots and the processing used for the statistical analysis is a subtle change to the background subtraction method, which improved the consistency and accuracy of the production start measurement. However, plotted on the scale of Fig. 3d, the changes in magnitude are negligible. In summary, the example plots used are representative of the final statistics and are sufficient to fully illustrate the methods. The correlation coefficient between *L*_*GA*_ and *I*_*max*_ quoted in Fig. 3d is alculated from the final data used for modelling (n = 130), as displayed in the correlation matrix in Fig. S12 (Supplementary note 15).

**Supplementary table 10:**
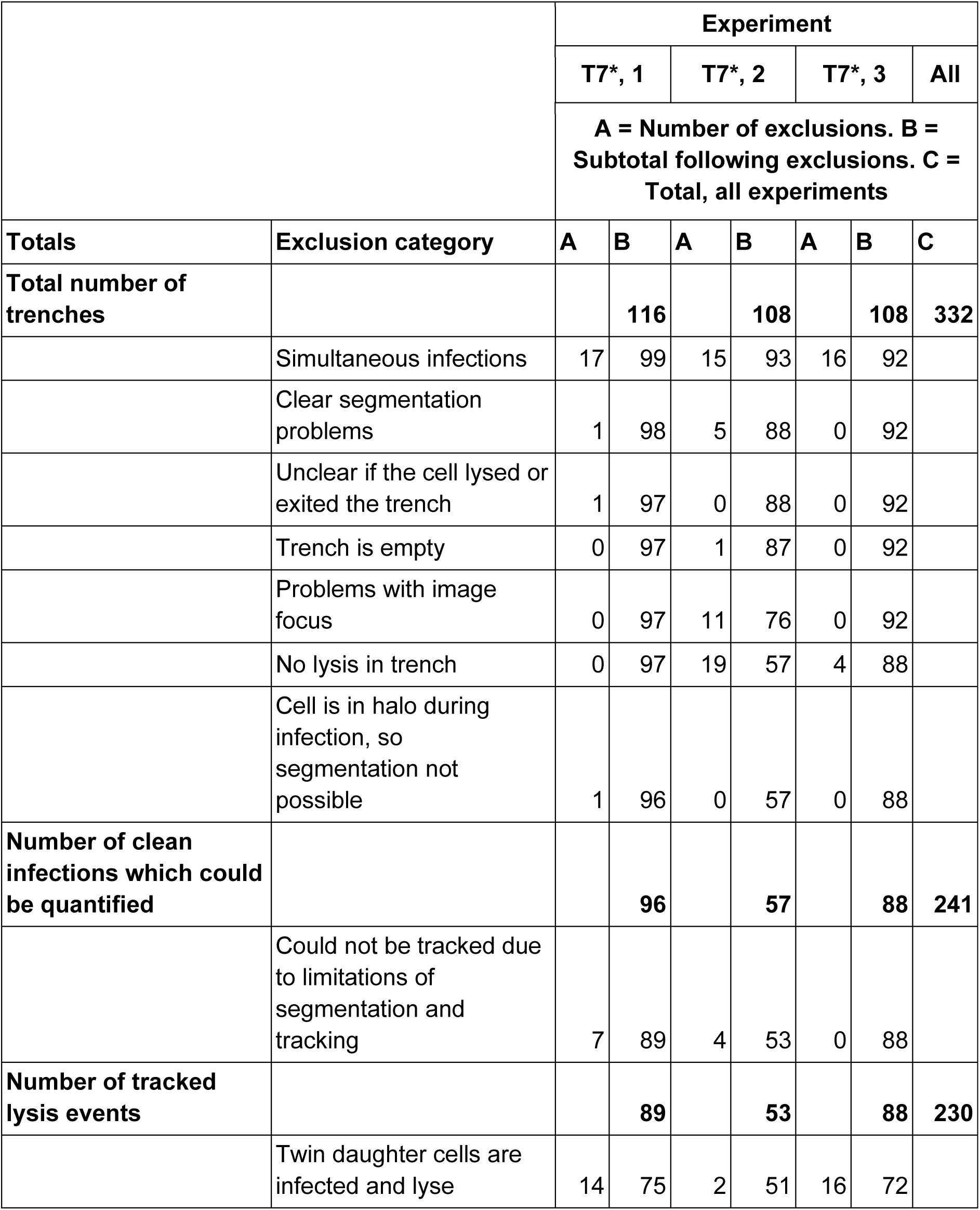
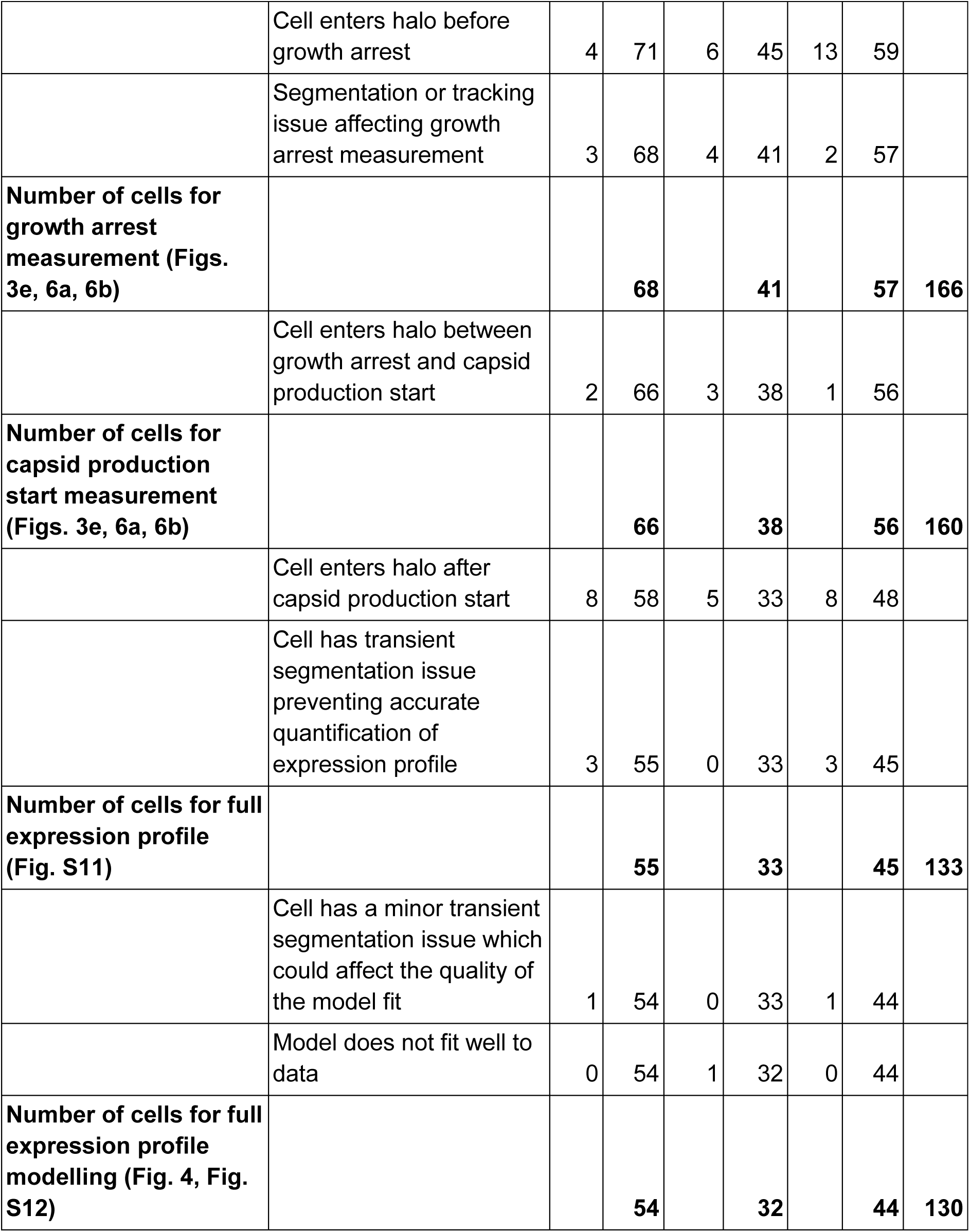
Data curation for Figs. 3 and 4.

|  |  | Experiment |  |  |  |  |  |  |
| --- | --- | --- | --- | --- | --- | --- | --- | --- |
|  |  | T7*, 1 |  | T7*, 2 |  | T7*, 3 |  | All |
|  |  | A = Number of exclusions. B = Subtotal following exclusions. C = Total, all experiments |  |  |  |  |  |  |
| Totals | Exclusion category | A | B | A | B | A | B | C |
| Total number of trenches |  |  | 116 |  | 108 |  | 108 | 332 |
|  | Simultaneous infections | 17 | 99 | 15 | 93 | 16 | 92 |  |
|  | Clear segmentation problems | 1 | 98 | 5 | 88 | 0 | 92 |  |
|  | Unclear if the cell lysed or exited the trench | 1 | 97 | 0 | 88 | 0 | 92 |  |
|  | Trench is empty | 0 | 97 | 1 | 87 | 0 | 92 |  |
|  | Problems with image focus | 0 | 97 | 11 | 76 | 0 | 92 |  |
|  | No lysis in trench | 0 | 97 | 19 | 57 | 4 | 88 |  |
|  | Cell is in halo during infection, so segmentation not possible | 1 | 96 | 0 | 57 | 0 | 88 |  |
| Number of clean infections which could be quantified |  |  | 96 |  | 57 |  | 88 | 241 |
|  | Could not be tracked due to limitations of segmentation and tracking | 7 | 89 | 4 | 53 | 0 | 88 |  |
| Number of tracked lysis events |  |  | 89 |  | 53 |  | 88 | 230 |
|  | Twin daughter cells are infected and lyse | 14 | 75 | 2 | 51 | 16 | 72 |  |
|  | Cell enters halo before growth arrest | 4 | 71 | 6 | 45 | 13 | 59 |  |
|  | Segmentation or tracking issue affecting growth arrest measurement | 3 | 68 | 4 | 41 | 2 | 57 |  |
| **Number of cells for growth arrest measurement (Figs. 3e, 6a, 6b)** |  |  | **68** |  | **41** |  | **57** | **166** |
|  | Cell enters halo between growth arrest and capsid production start | 2 | 66 | 3 | 38 | 1 | 56 |  |
| **Number of cells for capsid production start measurement (Figs. 3e, 6a, 6b)** |  |  | **66** |  | **38** |  | **56** | **160** |
|  | Cell enters halo after capsid production start | 8 | 58 | 5 | 33 | 8 | 48 |  |
|  | Cell has transient segmentation issue preventing accurate quantification of expression profile | 3 | 55 | 0 | 33 | 3 | 45 |  |
| **Number of cells for full expression profile (Fig. S11)** |  |  | **55** |  | **33** |  | **45** | **133** |
|  | Cell has a minor transient segmentation issue which could affect the quality of the model fit | 1 | 54 | 0 | 33 | 1 | 44 |  |
|  | Model does not fit well to data | 0 | 54 | 1 | 32 | 0 | 44 |  |
| **Number of cells for full expression profile modelling (Fig. 4, Fig. S12)** |  |  | **54** |  | **32** |  | **44** | **130** |

#### Supplementary note 15 Mathematical model for YFP expression data and fit to experimental time series

To model the YFP fluorescence signal from single-cell experiments, we develop a simple model that captures the initial exponential growth of mRNA, followed by the ribosome-limited translation activity of the cell. The model assumes that: (i) the total fluorescence signal is proportional to the number of YFP proteins, *P*, (ii) the viral genome, *g*, is growing exponentially at a rate *λ*_1_, (iii) the T7 RNAP is highly abundant/efficient, so that it is not the limiting factor in transcription, which proceeds at maximum rate *λ*_m_, (iv) ribosomes are limited resulting in a first order Hill function for protein production with a maximum translation rate *λ*_2_, where *K* is the mRNA concentration at half the maximum translation rate. We are neglecting any delay in protein production due to folding time. The model can absorb any potential mRNA degradation rate in the exponential growth rate, *λ*_1_.

Following the assumptions of the model, we find that:

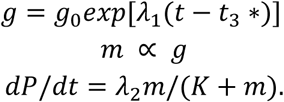

The system of equations can be solved analytically, leading to an exponentially growing number of mRNA molecules, *m* = *m*_0_*exp*[*λ*_1_(*t* − *t*_3_ ∗)], where *m*_0_ is a constant, and two useful functional forms for *dP*/*dt* as a function of *P* and of *P* as a function of time:

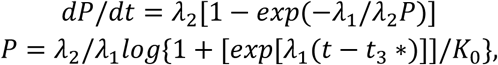

where *K*_0_ = *K*/*m*_0_.

We fit these expressions to the experimental time-series of infected cells. More specifically, the equation for *dP*/*dt* is first used to estimate *λ*_1_ and *λ*_2_, as these are the only two free parameters (Fig. 4a). These estimated values are then used to fit the experimental time series of YFP intensity with the equation for *P*(*t*) to infer *t*_3_ ∗ and *K*_0_ (Fig. 4a). From the value of *t*_3_ ∗ we then calculate the production time as the period of time between *t*_3_ ∗ and cell lysis (*t*_5_). Fig. S12 shows the correlation plots and distribution of all experimental observables and model parameters, a subset of which are reported in Fig. 4b-c.

**Fig. S12:**
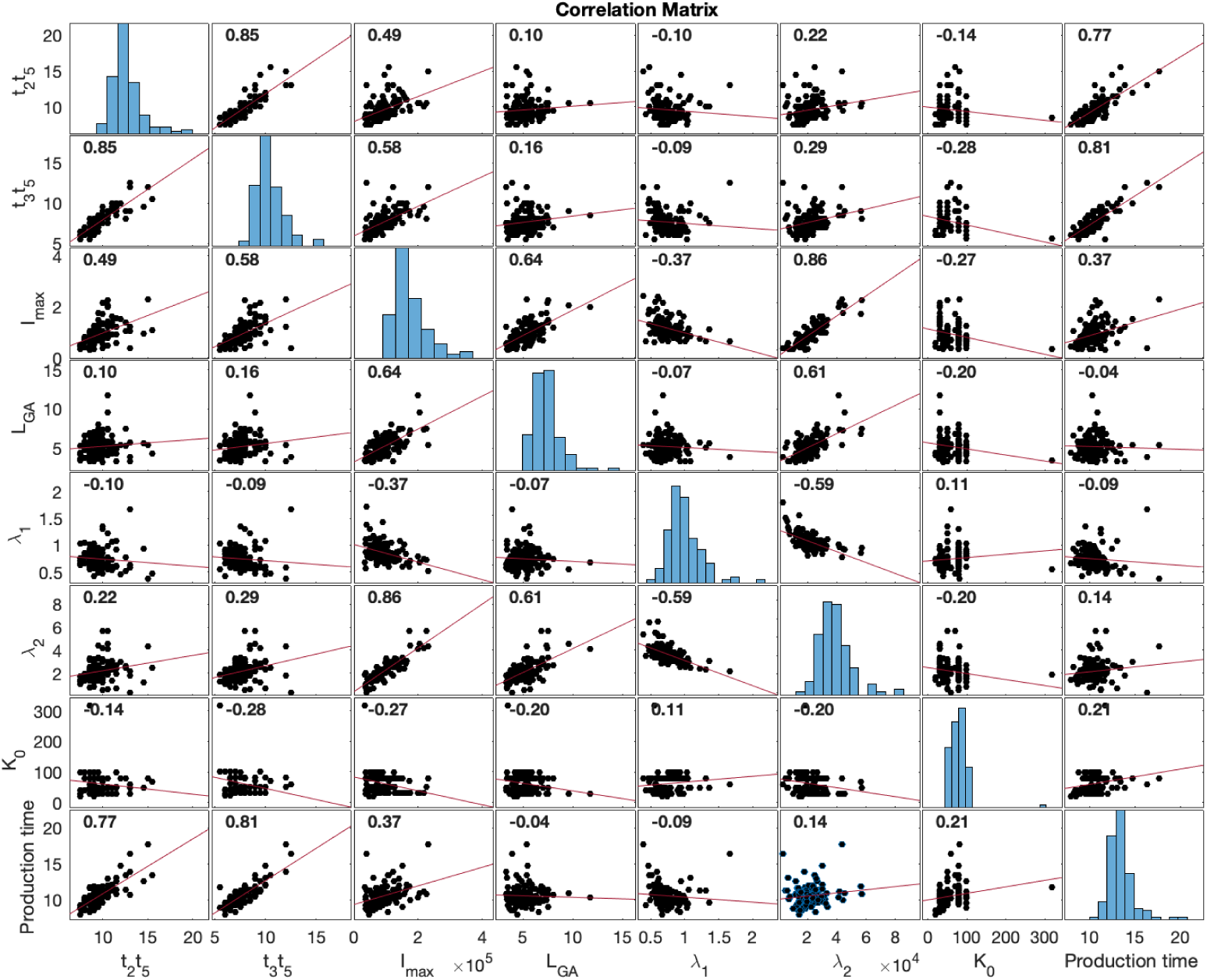
Correlation plots between experimental observables and inferred model parameters. A total of 130 expression profiles are fit as described above to infer the model parameters *λ*_1_, *λ*_2_, *t*_3_ ∗ and *K*_0_. Correlations between each of these parameters and the experimental observables *t*_2_*t*_5_ (time from growth arrest to lysis), *t*_3_*t*_5_(time between empirical start of protein production and lysis), *I*_*max*_ (maximum YFP) and *L*_*GA*_ (cell length at growth arrest), are reported together with their corresponding Pearson correlation coefficients *r*.Histograms summarising the distribution of each quantity are also shown on the diagonal.

#### Supplementary note 16 Perforation occurs in all observed lysis events

Fig. S13 shows the time series of mean phase contrast intensity in 36 lysing cells, demonstrating how in all cells, phase brightness is constant until the point of perforation at time *t*_4_. Following perforation, there is commonly a small initial surge in material loss, and then a period of steady material loss. After an average of 6.56 s has passed after perforation, lysis begins, leading to a rapid and complete breakdown of the cell envelope.

Only 36 events are used to produce Fig. S13, rather than the 42 used for Fig. 5e. The difference is because for six of the events used to plot Fig. 5e, a neighbouring cell moved into the position of the static mask as the cell lysed. This did not affect the calculation of perforation duration for those six events, but they have been omitted from Fig. S13 as the intensity data following lysis contains errors due to cell movement as described. Further details are provided in Supplementary note 17.

**Fig. S13:**
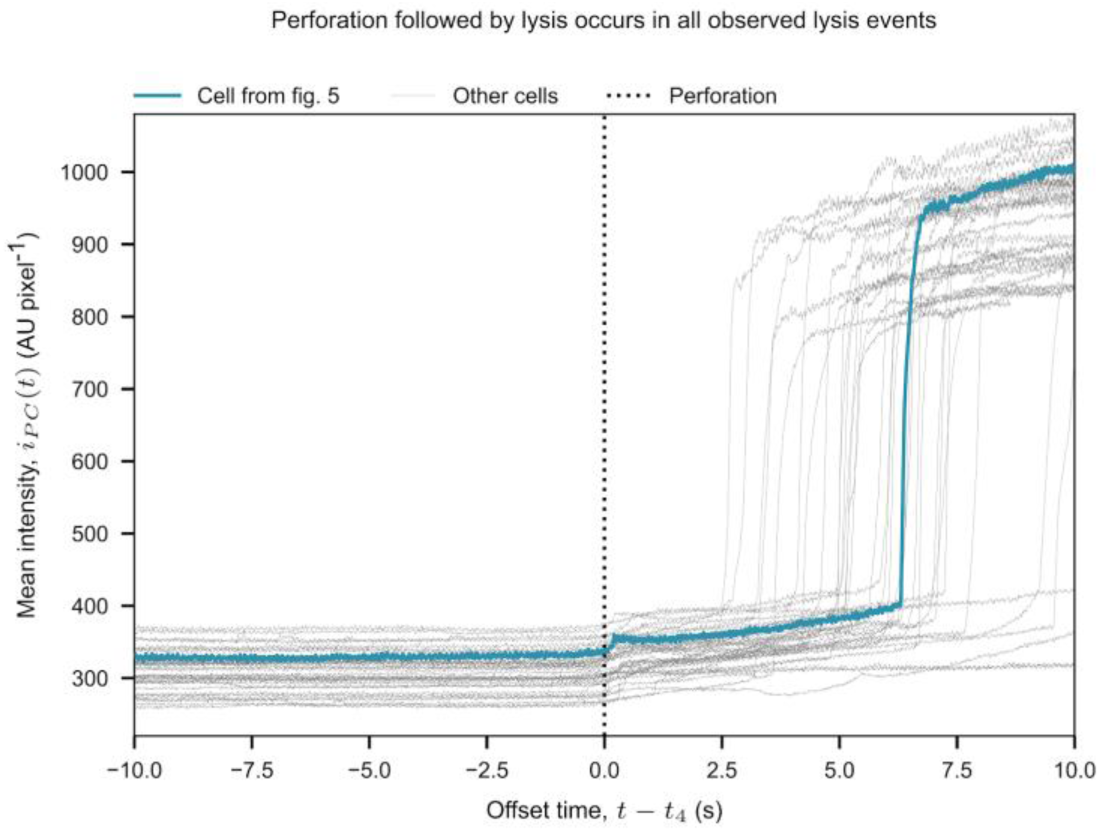
Perforation precedes all observed lysis events. A total of 36 time series showing the mean phase contrast intensity within each cell before and during lysis. All time series have been offset by the detected perforation time, *t*_4_. The blue line corresponds to the event in Fig. 5d, and the grey lines are 35 other cells.

#### Supplementary note 17 Data curation for perforation analysis

The criteria and event numbers used to curate data for the perforation analysis are summarised in Supplementary table 11. The analysis of perforation data uses static, hand drawn masks which do not move between frames. This is why the movement of the cell or of adjacent cells can lead to exclusion from the data set, because in these cases, the mask is no longer aligned with the cell of interest.

**Supplementary table 11:** Data curation for Fig. 5.

| Totals | Exclusion category | Number of exclusions | Subtotal following exclusions |
| --- | --- | --- | --- |
| **Total number of lysis events imaged** |  |  | **48** |
|  | The nearby lysis of another cell during perforation interferes with the measured signals | 1 | 47 |
|  | Part of cell cropped from image by the border of the camera region of interest | 2 | 45 |
|  | The cell moves during perforation, corrupting the signal | 3 | 42 |
| **Total number of perforation durations measured (Fig. 5e)** |  |  | **42** |
|  | Adjacent cell moves into the position of the lysing cell as it lyses, distorting the phase contrast signal during cell envelope breakdown | 6 | 36 |
| **Total number of full phase contrast signals (Fig. S13)** |  |  | **36** |

#### Supplementary note 18 Bulk one-step lysis curve experiments

Population level measurements of the mean lysis time of wild type T7 and T7* were carried out according to the ‘one-step growth curve’ or ‘lysis curve’ protocol^67^. An overnight culture of SB8 (Table 1) was diluted 1:100 into 1 ml of fresh LB Miller (Invitrogen) and placed in a shaking incubator at 37 °C, 250 rpm. After 2 h this culture was inoculated with 1 µl of bacteriophage stock, kept at 5 °C. At the same time, a second 1:100 dilution of the overnight culture was made, and both cultures were incubated for 2 h, yielding one dense bacterial culture (order 10^8^ cells ml^−1^, OD_600_ of 0.8), still in exponential phase, and one clear culture with few remaining cells but a very high density of bacteriophage (order 10^10^ PFUs ml^−1^). Both cultures were transferred to centrifuge tubes and spun at 6,000 g for 5 min. In the bacterial culture, the supernatant was removed and the pellet of cells was resuspended in 1 ml of fresh LB. In the phage culture, the supernatant was transferred to a new tube and any pellet of cell debris was discarded.

At time t = 0, the cell culture was inoculated with 10 µl of bacteriophage culture (approximate multiplicity of infection (MOI) of 1), vortexed, and placed in a bench-top shaking incubator at 37 °C, 300 rpm. The remaining bacteriophage culture was discarded. 1 min was allowed for adsorption to take place, after which the culture was serially diluted in LB down to two ‘active’ cultures ‘D6’, and ‘D7’, corresponding to dilution factors of 10^6^ and 10^7^ respectively. D6 and D7 were kept in the bench-top incubator (37 °C, 300 rpm), with 20 µl samples taken of each every 2.5 min and the bacteriophage density determined by plaque assay (Methods).

The population-level mean lysis time was determined by fitting a Gaussian cumulative distribution function (sigmoid) to the data. In some experiments, the number of PFUs was observed to decay sharply after the step, which may be due to multiple virions adsorbing to the same bacterial cell, thus concealing their true number. In these cases, the dataset was truncated after the step. Three replicates were carried out for each bacteriophage, each on a different day starting from a different clonal population of bacteria.

The three lysis times measured for wild-type T7 infecting SB8 were 16.7 min, 13.8 min, and 13.9 min, giving a grand mean and standard deviation of 14.8 ± 1.3 min. The three lysis times measured for T7* infecting SB8 were 18.6 min, 22.9 min, and 18.8 min, giving a grand mean and standard deviation of 20.1 ± 2.0 min. The measured values for both phages agree to within the experiment’s 2.5 min resolution.

**Fig. S14:**
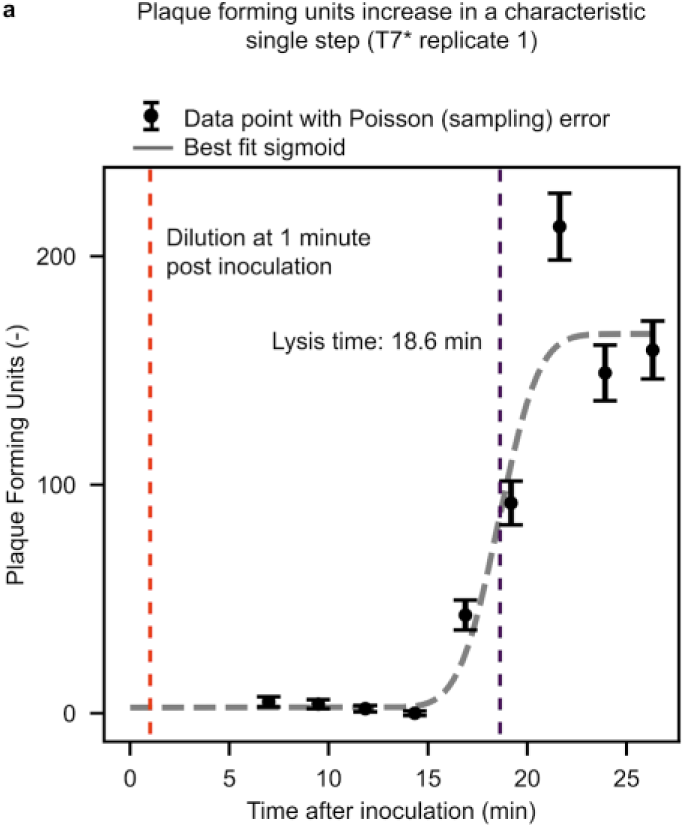
One-step growth curve for population level measurements of lysis time. **a.** Example one-step growth curve (lysis curve) used for population-level lysis time measurement. Three such datasets were obtained for each of T7 and T7*, and a sigmoid was fitted to each. The centre of the sigmoid is taken to represent the population mean lysis time, in this case 18.6 min after inoculation. Error bars represent the Poisson error arising from sampling active cultures. Note that this method cannot distinguish between time-to-adsorb and lysis time post adsorption. In theory the underlying variation in lysis time can be inferred from the slope of the sigmoid, however in practice this is complicated by the one minute adsorption window, and the experiment’s 2.5 min resolution.

Fig. S15 shows population level lysis time measurements from one-step growth curves for four phage-bacteria combinations. The two phage are T7 and T7*, as used throughout this study. One bacterial strain is SB8, used in all experiments in this study. The other is *E. coli* MG1655 7740, the wild type background strain of SB8.

**Fig. S15:**
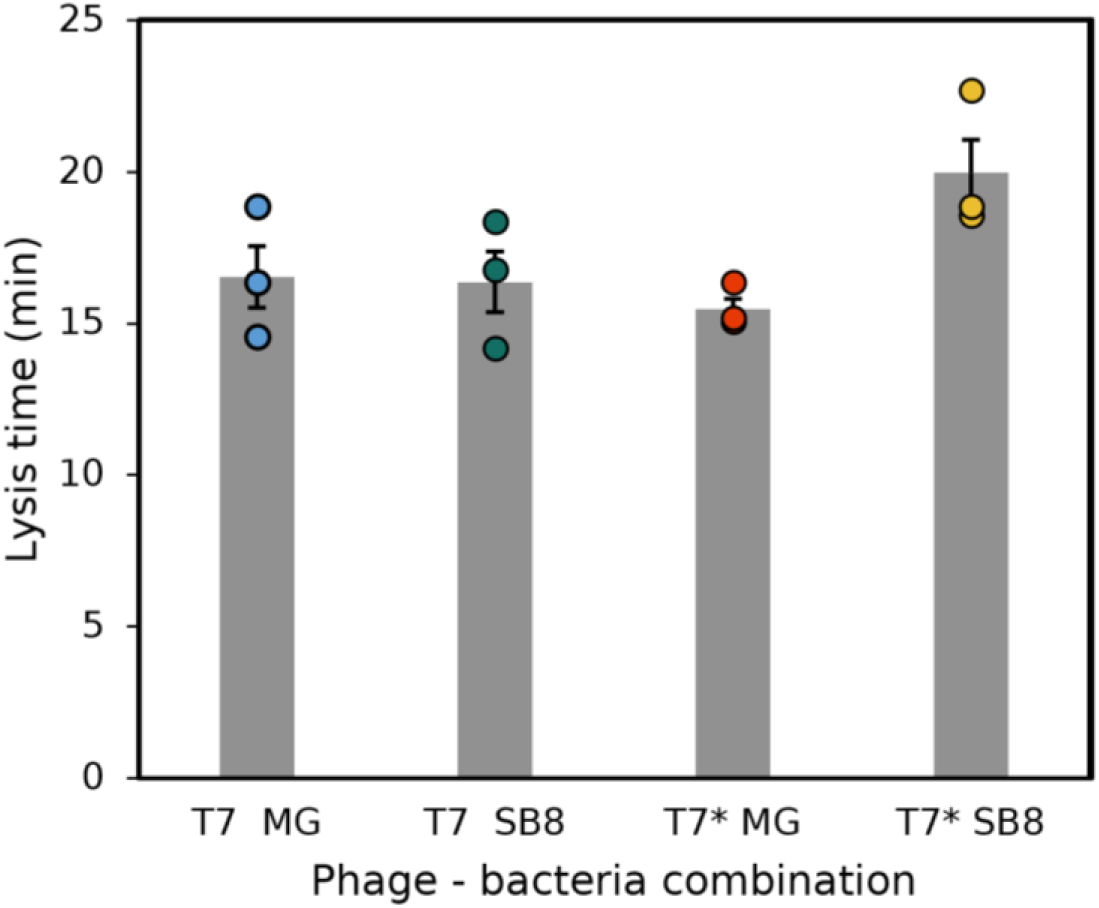
Lysis times observed in one-step growth curves. Three biological replicates are shown for each phage-bacteria combination. The error bars show ± 1 standard error of the mean. The experimental method is the same as described for Fig. S14.

#### Supplementary note 19 Serial passage simulation of phage competition

The stochastic, agent-based simulation was written in Python 3, based on earlier computational work^54^, and carried out on the high performance computing cluster at University of Cambridge (CSD3). The simulation’s dynamics are derived from the following infection kinetic equations^52^:

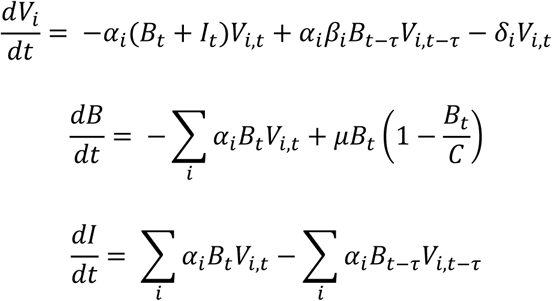

In which *B* is the concentration of uninfected bacteria, *I* the concentration of infected bacteria, *V*_*i*_ the concentration of free phage of type *i*, *C* the bacterial carrying capacity, *μ* the growth rate of healthy bacteria, *X*_*t*_ quantity *X* evaluated at time *t*, *⍺*_*i*_ the adsorption rate of phage *i*, *β*_*i*_ the burst size of phage *i*, *τ*_*i*_ lysis time of phage *i*, and *δ*_*i*_ the free virion decay rate of phage *i*.

The simulation is initialised with 2 pools of 100 phage each and 10,000 susceptible cells, in a simulated well mixed volume of 10^−5^ ml. Each bacterium is a Python object and maintains two internal clocks, one representing its own cycle of growth and division, and the other representing a count down to lysis. All bacteria begin the simulation in the ‘uninfected’ state, at random points in their cell cycle, with their ‘lysis clock’ stopped.

In each simulation time-step, ‘bacterial growth’, ‘adsorption’, ‘infection’, ‘lysis’, and ‘decay’ substeps occur. First, in the growth step, uninfected bacteria reduce their remaining time-to-fission by one increment of time, *Δt*. If time-to-fission falls below zero, a new bacterium is added to the simulation and both bacteria reset their division clocks, drawing their doubling time from a normal distribution with mean 1/*μ*.

Next, in the adsorption step, a number of infecting phage of each type, *ΔV*_*i*_ is drawn from a Poisson distribution with mean corresponding to the expected value *⍺*_*i*_(*B*_*t*_ + *I*_*t*_)*V*_*i*,*t*_*Δt* in a well-mixed population. The infecting phage are removed from the pool of free phage, and *ΔV* bacteria, whether infected or uninfected, are chosen randomly, uniformly and with replacement to which the phage adsorb. Each bacterium tracks how many phages of each type have adsorbed to it.

In the infection step, any bacterium which is still ‘uninfected’, but has at least one phage adsorbed to it changes state to ‘infected (wild type)’ or ‘infected (mutant)’ according to the nature of the adsorbed phages. In the case that a cell is infected by phages of both types, it chooses randomly in proportion to the number of adsorbed phages of each type. When a cell becomes infected it sets its internal countdown-to-lysis to a time drawn from a normal (or skew-normal) distribution with mean *τ*. The standard deviation and skew of this distribution depend on the phage strain. The wild type strain had a fixed standard deviation and zero skew throughout all simulations, mutant 1 varied the standard deviation across simulations and had zero skew, and mutant 2 had the same standard deviation as wild type but varied in skew across simulations. Supplementary table 12 lists all parameters used.

In the lysis step, infected cells reduce their time-to-lysis by the time increment *Δt*. If time-to-lysis falls below zero, the cell is deleted from the simulation, and a number of phage (wild type or mutant, according to which type of infection the cell had suffered) are released, drawn from a normal distribution with mean *β*.

Finally, in the decay step, each phage strain loses a number of free phage drawn from a Poisson distribution with mean *V*_*i*_*δ*_*i*_*Δt*.

Once the number of cells has dropped below 1% of its starting value, a ‘bottleneck’ is carried out, simulating the transference of a small quantity of the culture to a fresh bacterial stock. The bottleneck is implemented by a 1000-fold dilution, with each phage and cell in the simulation having a 99.9% chance to be deleted, followed by adding 10,000 new susceptible cells and the stochastic process above repeats. The whole procedure is repeated until, immediately following a bottleneck, one phage pool outnumbers the other 70:30, at which point it is declared the winner and the simulation stops. If neither phage is declared the winner after 600 min of simulated time, the simulation ends in a tie.

The simulation is carried out using the same parameter values for the wild type phage in every instance, and a range of values for the mutant. Values used are presented in Supplementary table 12. For each set of values, 25 independent runs are performed and the results are tallied to arrive at a fixation rate for the mutant phage, as the proportion of runs in which the mutant won.

**Supplementary table 12:** Serial passage simulation parameters.

|  |  |
| --- | --- |
| ***Simulation parameters*** |  |
| Maximum time | 600 min |
| Volume | $10^{-5}$ ml |
| ***Bacteria parameters*** |  |
| Initial number | 10,000, corresponding to a cell density of $10^9$ ml<sup>-1</sup> |
| Carrying capacity | 100,000, corresponding to a cell density of $10^{10}$ ml<sup>-1</sup> |
| Doubling time | Drawn from a normal distribution with mean 20 min, standard deviation 2.5 min. Positive definite. |
| Steric limit | Maximum number of phages that can adsorb to a single bacterial cell: 100. |
| ***Phage parameters*** |  |
| Initial number | 100 wild type, 100 mutant |
| Adsorption rate | $3 \times 10^{-9}$ ml min<sup>-1</sup> |
| Burst size | Drawn from a normal distribution with mean 150, standard deviation 50. Positive semidefinite. |
| Lysis time (wild type) | Drawn from a normal distribution with mean 17, standard deviation 2.5. Positive definite. |
| Lysis time (mutant 1) | Drawn from a normal distribution with mean 17, standard deviation varied from 2.0 to 3.0. Positive definite. |
| Lysis time (mutant 2) | Drawn from a skew-normal distribution with mean 17, standard deviation 2.5, skew varied from -0.88 to +0.88. Positive definite. |
| Decay rate | 0.00013 min<sup>-1</sup> |

#### Supplementary note 20 Experimental parameters for single-cell infection assay

Supplementary table 13 summarises the seven experiments which were used for Figs. 1 to 6. The titres of phage used in each experiment are such that the effective MOI is less than one, and therefore it is likely that any given infection is caused by a single phage.

**Supplementary table 13:** Parameters used for each experiment.

| Experiment | Figures used in | Phage | Titre in media (PFU/μL) | Media flow rate (μL/min) |
| --- | --- | --- | --- | --- |
| T7, injection | 2 | T7 | $1 \times 10^6$ | 0 |
| T7*, injection | 2 | T7* | $1 \times 10^6$ | 0 |
| T7, perforation | 5 | T7 | $1 \times 10^6$ | 5 |
| T7*, 1 | 1, 3, 4, 6a, 6b, 6c | T7* | $1 \times 10^5$ | 5 |
| T7*, 2 | 3, 4, 6a, 6b, 6c | T7* | $2 \times 10^5$ | 5 |
| T7*, 3 | 3, 4, 6a, 6b, 6c | T7* | $5 \times 10^4$ | 5 |
| T7, 1 | 6c | T7 | $1 \times 10^5$ | 5 |

#### Supplementary note 21 Experimental protocol adjustments for genome injection dynamics experiments in Fig. 2

The microfluidic experiments used in Fig. 2 followed a similar form to that outlined in the Methods section, but some changes were made in an attempt to improve throughput and the signal to noise ratio. The same cell strain as all other experiments (SB8) was used. Microfluidic trenches with the same main trench and side trench widths were used, but with a 48 μm main trench length. The shorter trenches contained fewer cells, meaning that cells closer to the trench exit had a longer residence time in the trench when compared to longer trenches containing more cells. This increased the chance of observing lysis before infected cells were pushed out of the trench. For the same reason of increasing cell residence time, we aimed to sparsely load the trenches with cells by only leaving the dense cell solution in the feeding lane for a short period when loading the device.

Several changes were made to improve the signal-to-noise ratio. One was the use of a brightfield objective, used with the aim of allowing more of the emission light to pass through than would be the case with a phase contrast objective. The new objective was also of 60x rather than 100x magnification, which had the benefit of increasing throughput. Furthermore, a longer exposure time of 100 ms was used for the orange channel imaging. For the stained phage experiments used for Fig. 6, the time from the creation and SYTOX Orange staining of the phage lysate to the experiment ranged between seven and 101 days. We observed that the biological activity of the phage lysates had good stability over several months when stored in LB Miller at 4 °C. Over the four experiments there was no clear pattern between the gap from staining to experiment and the quality of the staining. Despite this, we aimed to exercise tighter control over this gap. The times between staining and experiment for Fig. 2 were five and six days for the T7 and T7* experiments respectively.

Other changes were made in an attempt to increase throughput. A higher phage titre of 1.0 × 10^6^ PFU μL^−1^ was used to increase the infection rate per unit time. Furthermore, a different flow regime was used. After sparse loading, a flow of growth media was supplied for 2 h, before switching to a phage media flow at 15 μL min^−1^. When phage media reached the trenches and the SYTOX orange signal observed in the media had reached steady state, the flow was switched off. The primary aim of doing this was to reduce the transfer of phage progeny released by lysis to other trenches, to maximise the ratio of stained to unstained phage infections, but it is unclear if this was successful as the effect is difficult to measure. The time between the flow being switched off and the end of the experiment was at most 1.7 h. During this time, the growth of cells appeared to be normal, with media availability aided by a low cell count due to sparse loading and phage lysis. Furthermore, the tubing, with a 0.5 mm internal diameter, has a volume per unit length that is 65 times larger than the internal volume of the microfluidic device per unit length, providing a substantial reservoir of media which could diffuse to replace any consumed nutrients.

#### Supplementary note 22 High frame rate image acquisition and nucleoid compaction in T7 infected cells

To capture high frame rate images of the lysis process for Fig. 5, we found it was necessary to use a small region of interest (ROI) for the camera, the size of a single trench, otherwise the imaging process was limited to lower frame rates. In order to gain sufficient throughput, we found trenches where multiple infected cells were due to lyse as a consequence of the phages released from a first-round lysis event in the trench, and started the imaging on these cells to capture images of multiple lysis events. We were able to spot infected cells in the phase contrast images due to the appearance of phase bright foci within the cells. We explain more about the cause of the phase bright foci below. After most cells had lysed, we would find another trench with multiple cells about to lyse, and begin imaging there. Five trenches were imaged in this manner over one microfluidic experiment, capturing the lysis of 48 cells. For this reason, cells imaged for the perforation analysis were likely infected at an MOI greater than one. More information regarding how the 48 lysis events have been used can be found in Supplementary note 17.

In the high frame rate movies (Supplementary movie 4), we can see that infected cells develop a phase-bright feature at their centre. Fig. S16 shows how phase bright foci develop in T7 infected cells during the late stage of infection. The phase bright foci are clearly visible using different objective lenses (Fig. S16a, 100x oil, S16b, 40x air) and across different experiments, strongly indicating the structures are real and not an artefact. An example data from such an experiment, Fig. S16b, shows cells with a nucleoid-associated protein with an RFP fusion infected by a genetically modified T7 phage (TY001) similar to T7*. TY001 was a modified wild type T7 phage, where an mEYFP cassette under the control of an *E. coli* promoter was inserted downstream of gene 12. We thank Alberto Scarampi for providing us with materials for constructing this phage. The reasons for the modification were separate to the observation presented here, but this phage effectively lysed cells similar to T7* and the wild type T7. The cell strain visualised in Fig. S16b is LY119, which is an MG1655 E. coli strain (MS388) with a hupA-mCherry-FRT-kan-FRT modification^80^, as described by Cayron et al. This strain was kindly provided by the Lesterlin lab. The appearance of the phase-bright foci correspond to the coalescence and brightening of the regions showing the nucleoid label in the RFP channel.

**Fig. S16:**
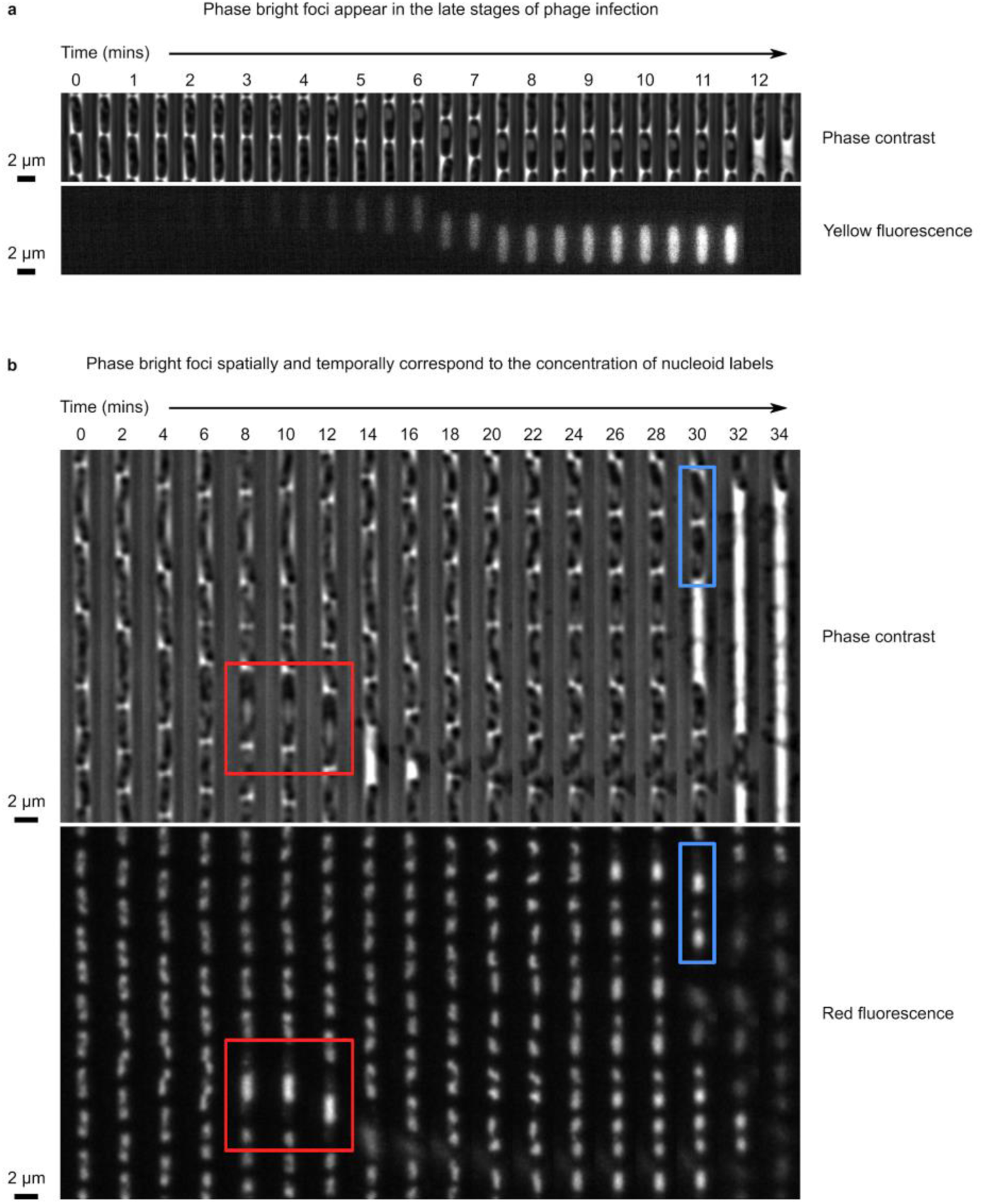
Phase bright foci appear during phage infection. **a**. A kymograph of phase contrast images (top panel), showing the appearance of a phase-bright focus in a cell infected with T7*. The corresponding yellow fluorescence image (bottom panel) shows the expression of YFP from the phage genome. There is an approximate correspondence between the appearance of the phase-bright focus and the appearance of the YFP signal, which matches our current experience of the phase-bright focus appearing in the later stages of infection. **b**. Phase contrast (top panel) and red fluorescence (bottom panel) kymographs showing an MG1655 strain with an RFP-tagged nucleoid protein being infected by a T7 phage with similar modifications to T7*. In uninfected cells, the nucleoid label appears as two lobes, reflecting the healthy spatial distribution of the nucleoid^80^. In the late stages of T7 phage infection, these lobes become brighter, and often coalesce into a single region or become asymmetric in their intensity. The red box highlights a particularly pronounced example. Following the lysis of the cell in the red box, secondary infections in a number of cells show phase bright foci and their corresponding nucleoid labels. Two of these cells are highlighted in the blue box, although phase bright foci and concentrated nucleoid labels can be seen in all the cells infected by phage released from the cell in the red box.

#### Supplementary note 23 List of single-cell features from image analysis

**Supplementary table 14:** Properties of infected cells.

| Property | Unit | Description |
| --- | --- | --- |
| Time, $t$ | min | Time from the start of imaging. |
| Interval between frames, $\Delta t$ | min | Time between consecutive frames. The values for each experiment are defined in Table 4. |
| Cell length, $L(t)$ | $\mu\text{m}$ | Cell length from scikit-image regionprops function. |
| Cell area, $A(t)$ | pixel | Cell area from scikit-image regionprops function. |
| Raw total intensity of YFP, $I_{raw}(t)$ | Arbitrary units (AU) | Sum of YFP pixel intensities over the cell area, from scikit-image regionprops function. |
| Raw mean intensity of YFP, $i_{raw}(t)$ | AU pixel<sup>-1</sup> | Mean of YFP pixel intensities over the cell area, from scikit-image regionprops function. |
| $\ln(L(t))$ | - | Natural log transform of the cell length. |
| $[\ln(L(t))]'$ | - | Result of applying the Savitzky-Golay filter<sup>79</sup> to $\ln(L(t))$ using a third-order polynomial and a window size of eight data points. |
| Growth rate, $\lambda(t)$ | min<sup>-1</sup> | $\lambda(t) = \frac{d}{dt} [\ln(L(t))]'$ |
| Growth arrested cell length, $L_{GA}$ | $\mu\text{m}$ | $L_{GA} = \frac{\Delta t}{t_5 - 2\Delta t - t_2} \sum_{t=t_2}^{t_5-2\Delta t} L(t)$ |
| Mean background intensity of YFP, $i_{bg}$ | AU pixel<sup>-1</sup> | The mean background intensity is a position dependent quantity. The background YFP profile is calculated along the length of the trench approximately 5 min before YFP expression begins. The value of $i_{bg}$ for a cell in a given position is then taken from this background profile. |
| Total background YFP intensity, $I_{bg}(t)$ | AU | $I_{bg}(t) = i_{bg} \cdot A(t)$ |
| Total intensity, background corrected, $I(t)$ | AU | $I(t) = I_{raw}(t) - I_{bg}(t)$ |
| Mean phase contrast intensity in a cell, $i_{PC}(t)$ | AU pixel<sup>-1</sup> | Mean of the phase contrast pixel intensities over the cell area, from the Fiji (ImageJ) function "Plot Z-axis Profile" over a hand drawn segmentation of the lysing cell. |

#### Description of supplementary movies

##### Supplementary movie 1 Phage Infection and subsequent cell lysis in the microfluidic device

Supplementary movie 1 shows five trenches of the microfluidic device (mother machine), loaded with *E. coli* strain SB8 cells, as T7* phages infect and lyse them over time. Phage-containing growth media flows from left to right through the feeding lane at the bottom, diffusing into the trenches and allowing phages to infect individual cells. When a cell becomes infected, YFP is expressed from a gene in the phage genome, causing the cell to turn yellow, before they eventually lyse. The first infected cell is in the second trench from the right, displaying a yellow signal at approximately 00:47:00 and lysing at 00:53:00. Phages from this lysis event subsequently infect and clear the trench of cells by 01:50:00. Cells in the other trenches are infected and cleared similarly until all trenches are clear by 09:53:00. The time stamp is in hh:mm:ss, and the scale bar size is 5 μm. Images were taken every 30 s.

##### Supplementary movie 2 Phage infection from adsorption to lysis in a single cell

Supplementary movie 2 shows a single trench of a mother machine device, in which a SYTOX Orange stained T7* phage binds to an *E. coli* cell and injects its genome (indicated by the SYTOX Orange signal getting dimmer over time), causing a subsequent expression of YFP (appearance and increase of yellow signal) and eventually host cell lysis (loss of phase contrast and yellow signal). The position of the stained phage, which binds at 01:59:30, is indicated by an arrow. A YFP signal begins to form at 02:07:00, and increases in intensity until the cell lyses at 02:12:30. A sudden increase in SYTOX Orange intensity concurs with the lysis, likely due to residual SYTOX dye in the media binding to the released bacterial DNA. The time stamp is in units of hh:mm:ss, and the scale bar size is 2 μm. Images were taken every 30 s.

##### Supplementary movie 3 Adsorption and genome injection of a SYTOX Orange stained phage

Supplementary movie 3 shows a single trench of a mother machine device, in which a SYTOX Orange stained T7 phage binds to an *E. coli* cell and injects its genome. The position of the stained phage, which binds at 01:21:30, is indicated by an arrow. As the genome injection proceeds, the SYTOX Orange signal intensity decreases until the spot disappears entirely. The time stamp is in units of hh:mm:ss, and the scale bar size is 2 μm. Images were taken every 10 s.

##### Supplementary movie 4 High frequency imaging of phage-induced cell lysis

Supplementary movie 4 shows a single trench of a mother machine device, in which an infected *E. coli* cell undergoes T7 induced lysis. High frequency imaging (100 frames per second) was used to capture this process, which occurs on sub-second time scales. At 80.22 s, a significant rupture of the cell envelope occurs, which can be seen due to a plume of material (a darker region in the image) being ejected from the lower pole of the cell. The position of this rupture is indicated by an arrow. This rupture is promptly followed by the complete loss of material from the cell. The time stamp has units of seconds, and the scale bar size is 2 μm.

